# Variation in albumin glycation rates in birds suggests resistance to relative hyperglycaemia rather than conformity to the pace of life syndrome hypothesis

**DOI:** 10.1101/2024.07.02.600014

**Authors:** Adrián Moreno-Borrallo, Sarahi Jaramillo-Ortiz, Christine Schaeffer-Reiss, Benoît Quintard, Benjamin Rey, Pierre Bize, Vincent A. Viblanc, Thierry Boulinier, Olivier Chastel, Jorge S. Gutiérrez, Jose A. Masero, Fabrice Bertile, François Criscuolo

**Affiliations:** University of Strasbourg, CNRS, Institut Pluridisciplinaire Hubert Curien, UMR 7178, 67000 Strasbourg, France; National Proteomics Infrastructure, ProFi, FR2048 Strasbourg, France; Parc zoologique et botanique de Mulhouse, 68100 Mulhouse, France; Lyon University 1, UMR CNRS 5558, Laboratoire de Biométrie et Biologie Evolutive, 69622 Villeurbanne, France; Swiss Ornithological Institute, CH-6207 Sempach, Switzerland; CEFE, Montpellier University, CNRS, EPHE, IRD, Montpellier; Center of Biological Studies of Chizé (CEBC), UMR 7372 CNRS - La Rochelle University, 79360 Villiers-en-Bois, France; Ecology in the Anthropocene, Associated Unit CSIC-UEX, Faculty of Sciences, University of Extremadura, Badajoz, Spain

**Keywords:** birds, comparative method, glucose, glycation, life history, longevity

## Abstract

The pace of life syndrome hypothesis (POLS) suggests that organisms’ life history, physiological and behavioural traits should co-evolve. In this framework, how glycaemia (i.e., blood glucose levels) and its reaction with proteins and other compounds (i.e. glycation) covary with life history traits remain relatively under-investigated, despite the well documented consequences of glucose and glycation on ageing, and therefore potentially on life history evolution. Birds are particularly relevant in this context given that they have the highest blood glucose levels within vertebrates and still higher mass-adjusted longevity when compared to organisms with similar physiology as mammals. We thus performed a comparative analysis on glucose and albumin glycation rates of 88 bird species from 22 orders, in relation to life history traits (body mass, clutch mass, maximum lifespan and developmental time) and diet. Glucose levels correlated positively with albumin glycation rates in a non-linear fashion, suggesting resistance to glycation in species with higher glucose levels. Plasma glucose levels decreased with increasing body mass but, contrary to what is predicted to the POLS hypothesis, glucose levels increased with maximum lifespan before reaching a plateau. Finally, terrestrial carnivores showed higher albumin glycation compared to omnivores despite not showing higher glucose, which we discuss may be related to additional factors as differential antioxidant levels or dietary composition in terms of fibres or polyunsaturated fatty acids. These results increase our knowledge about the diversity of glycaemia and glycation patterns across birds, pointing towards the existence of glycation resistance mechanisms within comparatively high glycaemic birds.

## Introduction

The pace of life (POL) hypothesis [1–3] postulates that organisms behavioural/physiological characteristics and life histories have co-evolved in answer to specific ecological conditions forming pace of life syndromes (POLS) along a fast-slow continuum (but see [4] for the more recent consideration of other axes). According to the classical approach from life history theory, slower organisms would have, e.g. large body mass, late maturation, slow growth rate, small number of large offspring and high longevity, whit faster ones representing the opposite trait values (see [5–8]) within this continuum. Pace of life hypothesis added physiology to this continuum, with studies testing its predictions focusing often on metabolic rate (i.e. higher or lower metabolic rate corresponding to faster or slower pace of life, respectively; see e.g. [9–12]). However, key energy substrates related to metabolic performance, such as glucose concentrations in tissues, have been largely overlooked (but see [13–16]). Notably, glucose, a central energy source for many organisms, plays a pivotal role in metabolism and there are some indications that its circulating blood or plasma levels correlate positively with metabolic rate per gram of mass at the interspecific level [13,17], and with whole body metabolism at the intraspecific level ([18] for the case of non-diabetic humans).

Research focusing on plasma glucose becomes highly relevant as high glycaemia can entail costs that accelerate ageing. Along with other reducing sugars, glucose can react non-enzymatically with free amino groups of proteins, lipids or nucleic acids, a reaction known as glycation [19][20]. This reaction, after several molecular reorganizations, can lead to the formation of a plethora of molecules called Advanced Glycation End-products (AGEs) [21]. AGEs can form aggregates in tissues that are well known to contribute to several age-related diseases, such as diabetes (e.g., reviews by [22–25]). AGEs are also known to promote a proinflammatory state through their action on specific receptors called RAGEs [26–27].

Remarkably, birds show circulating glucose levels much higher than other vertebrate groups, on average almost twice as high as mammals [28]. These relatively high glucose levels might support the higher metabolic rates [29, 11] and body temperatures observed in birds compared to mammals [30–31]. In addition, elevated glycaemia is thought to be an adaptation to flight, providing birds with rapid access to an easily oxidizable fuel during intense bursts of aerobic exercise (see [32–33]), such as during take-off and short-term flapping flights [34–35]). However, the presence of high glycaemia in birds is also paradoxical, given their remarkable longevity compared to their mammalian counterparts — living up to three times longer than mammals of equivalent body mass [36, 11].

Several non-excluding hypotheses have been proposed to explain how birds apparently resist the pernicious effects of a high glycaemic state. One possibility is that birds might resist protein glycation through a lower exposure of lysine residues at the surface of their proteins [37]. Another hypothesis suggests that increased protein turnover may play a role [38–39]. Additionally, birds might benefit from more effective antioxidant defences [40–42] (although within Passeriformes, [16] shows no coevolution between glycaemia and antioxidant defences) or even the presence of “glycation scavengers” (i. e. molecules that bind glucose avoiding it to react with proteins) such as polyamines which circulate at high concentration [35]. Moreover, it is generally considered that RAGEs are not present in avian tissues, suggesting possible adaptive mechanisms that would allow birds to avoid the inflammatory consequences associated with RAGE activation [43–44]. However, a putative candidate for RAGE and a case of AGE-induced inflammation in birds were identified by Wein et al. [45], warranting further investigation on the occurrence of RAGEs in birds.

To date, although protein glycation levels have been measured in birds in several species, these results were mostly descriptive and concerned a small number of species [46–54]. Additionally, many of these studies employ commercial kits that were not specifically designed for avian species, making it challenging to interpret the results [53]. These limitations restrict our ability to draw general conclusions from these studies. Here, we performed a comparative analysis on primary data (for glycation and most of glucose values, see methods and ESM6) from 88 bird species belonging to 22 orders (see ESM6) and assessed whether and how bird glycaemia and glycation rates are linked to ecological and life-history traits (including body mass). While doing this, we also checked whether glycation levels are influenced by circulating plasma glucose concentrations. Finally, as diet can influence glycaemia (see e.g. [15, 55–56] but [14] found no significant effects) and glycation, we included it in our analyses. We hypothesized that carnivorous birds would exhibit higher glycaemia and glycation levels than omnivorous or herbivorous species, as diets with low-carbohydrates, high-proteins and high-fat are associated with comparatively high glycaemia and reduced insulin sensitivity in some vertebrates (e.g.; [57–59]). Accordingly, a recent comparative study in birds showed higher blood glucose levels in carnivorous species [56]. This may be attributed to high levels of constitutive gluconeogenesis, which has been confirmed in certain raptors [60–61]. Conversely, bird species with high sugar intake, such as frugivores or nectarivores, are expected to exhibit high glycaemia, as observed in hummingbirds [49] and Passeriformes with such diet [15] (although the opposite was found in [56]). In line with POLS hypothesis, we also hypothesized that species with a slower POL should exhibit a lower glycaemia and higher glycation resistance (glycation levels for a given glycaemia) compared to species with a faster POL. Hence, we included the following four parameters in our analyses of glycaemia and glycation levels: body mass, maximum lifespan, clutch mass and developmental time. Body mass is one of the main factors underlying life history, with higher body mass associated with “slower” strategies and vice versa [62]. Previous studies have reported a negative relationship between body mass and glycaemia [63, 34, 13–14, 56](but see [49, 64] with non-significant trends in birds, and [15] only for temperate species within Passeriformes). Therefore, we predicted that bird species that live longer, develop more slowly and invest less per reproductive event (see ESM6 for further justification of the chosen variables) should show lower plasma glucose levels and albumin glycation rates (after controlling for body mass, phylogeny, diet and glucose in the case of glycation; see methods).

## Materials and methods

### Species and sample collection

A total of 484 individuals from 88 species measured were included for this study (see **Table ESM6. 1**). A detailed list with the provenance of the samples, including zoos, a laboratory population, and both designated captures and samples provided by collaborators from wild populations, is provided in ESM3, with a textual description in ESM6.

Blood samples were collected from the brachial or tarsal vein, or from the feet in the case of swifts, using heparin-lithium capillaries or Microvette® (Sarstedt). As samples were collected mostly by different collaborators, handling times were not always recorded and could not be adjusted for, potentially rendering the results more conservative (for a more detailed discussion on potential stress effects on glucose, see ESM6), Samples were centrifuged at 4°C, 3500 g for 10 min and plasma was aliquoted when a large volume was available. Subsequently, they were transported on dry ice to the IPHC (Institut Pluridisciplinaire Hubert Curien) in Strasbourg and stored at -80° C until analysis. Glycaemia was measured in the laboratory on the remaining plasma after taking an aliquot for glycation assessment, using a Contour Plus^®^ glucometer (Ascensia diabetes solutions, Basel, Switzerland) and expressed in mg/dL. These of point-of-care devices have previously been assessed for its usage in birds [65–66], with several examples of its usage in recent literature related to the topic presented here (e.g. [14, 67–69]. We also performed an assay with this particular brand on a subset of 46 samples from this study, coming from nine species distributed across the whole range of glucose values (three species with “low” values, three with “medium” and three with “high”, from both captive and wild populations, with five individuals per species, except one including six), confirming a positive linear correlation (R^2^_marginal_=0.66; R^2^_conditional_=0.84, for a model including the species as random factor) of this device with Randox GLUC-PAP colourimetry kits (P-value<0.001, unpublished data). Due to technical issues, related to the incapability of the device to determine certain glucose values (not because of the glucose concentration, but perhaps the particular composition of the plasma samples from certain species), we could not determine glycaemia values of 95 individuals of those that we sampled and in which glycation levels were assessed (belonging to 40 species coming from different sources). For these species, if not a single individual had a glucose measurement (which was the case in 13 species), we obtained mean plasma glucose values reported for the species from the ZIMS database from Species360 (Species360 Zoological Information Management System (ZIMS) (2023), zims.Species360.org). This database provides plasma glucose data measured by colorimetric glucose kits on zoo specimens not neccessary corresponding to those in which we measured glycation values. The sample sizes for both glucose and glycation measurements, either from our measured individuals or from the ones from ZIMs, are reported on the ESM5.

### Glycation levels

Glycation levels for each individual were determined using liquid chromatography coupled to mass spectrometry (LC-MS), which is considered the gold standard for the assessment of protein glycation levels (see e.g. [70]) and has previously been used for birds [52–53, 71]. Given the relatively high time intrinsically taken by the employed methodology for analysing all the samples, linked to constraints in the access to the mass spectrometry devices, the whole set of samples of this study were analysed across several instances within a total timespan of less than 2 years (2021-2023). Only one sample from each individual was measured, given logistic limitations on the total number of samples that could be processed. Briefly, 3 µl of plasma were diluted with 22 µl of distilled water containing 0.1% of formic acid, followed by injection of 5 µl into the system. The glycation values used in the analyses represent the total percentage of glycated albumin, obtained by adding the percentages of singly and doubly glycated albumin. These percentages were calculated by dividing the areas of the peaks corresponding to each glycated molecule form (albumin plus one or two glucose) by the total albumin peak area (sum of the areas of glycated plus non-glycated molecules) observed in the spectrograms obtained from the mass spectrometry outcomes. These spectrograms represent the different intensities of signal for the components differentiated in the sample by their TOF (Time Of Flight), which depends on the mass to charge ratio (m/z) of the ionized molecules. More detailed information regarding functioning of the method and data processing can be found in [53]. In cases where albumin glycation values dropped below the limit of detection, resulting in missing data, these individuals (10 individuals from 4 species) were excluded from the statistical analyses, as outlined below.

### Data on ecology and life-history

Diets of individual species were extracted from the AVONET dataset ([72], coming from [73], and those adapted from [74]) and life history traits from the Amniote Database [75]. After determining that AVONET diet classifications did not align with our research needs, minor changes were made after consulting the original Wilman et al. [74] database (see ESM6).

For missing data on life history traits after this stage, we extracted values, in order of preference, from the AnAge database [76] or the Birds of the World encyclopaedia [77], by calculating mean values if an interval was given, and then averaging if male and female intervals were provided separately. For European species, maximum lifespan records have always been checked against the most recent Euring database [78], and from *Chionis minor* and *Eudyptes chrysolophus* (not available from the previous sources) they were extracted from the Centre d’Etudes Biologiques de Chizé, Centre National de la Recherche Scientifique (CNRS-CEBC) database. Several species from the zoo had maximum lifespan values available on the ZIMs from Species 360 (Species360 Zoological Information Management System (ZIMS) (2023), zims.Species360.org), so these were also compared with the data we had from other sources.

In the case of variables (other than maximum lifespan) for which two different sources provided different records, an average value was calculated. For the maximum lifespan value, available sources were cross-checked and the highest value was always used. In the absence of satisfactory support from another source, values for maximum lifespan indicated as being anecdotal and of poor quality have been excluded from the analyses. A table with the adequate citations for each value is provided as part of the online available data (see Data accessibility section). When the source is AVONET [72], this database is cited, and when it is Amniote Database [75], we cite the sources provided by them, so the references can be checked in [75]. A list with all the additional references not coming from any of these databases or not provided by their authors (and that are not already in the main text) is given as ESM4.

### Statistical analyses

All analyses were performed in R v.4.3.2 [79]. Alpha (α) ≤0.05 was reported as significant, and 0.05<α≤0.1 as trends. General Linear Mixed Models with a Bayesian approach (*MCMCglmm* function in R; [80]) were performed. All models were run with 6×10^6^ iterations, with a thinning interval of 100 and a burn-in of 1000. The models were simplified by eliminating quadratic terms where they were not significant and selecting models with lower AIC (Akaike Information Criterion) and BIC (Bayesian Information Criterion). Gaussian distribution of the response variables was assumed (after log_10_ conversion for glucose; see ESM6 for further discussions). The priors were established assuming variance proportions of 0.2 for the G matrix and 0.8 for the R matrix, with ν=2. Less informative priors with equal variance proportions for each partition (residual and random) gave similar results. Lower ν values (0.1, 0.2, 0.002) were also tested, without success (i.e. simulations aborted before the established number of iterations). For the phylogenetically controlled analyses, consensus trees (each with the species included in the model) were obtained by using the *consensus.edges* function (from the *phytools* package from R [81] from a total of 10,000 trees downloaded from Birdtree.org [82] (option “Hackett All species” [83]). For such purpose, a list of species with the names adapted to the synonyms available in such website was used (see ESM5), including the change of *Leucocarbo verrucosus* for *Phalacrocorax campbelli*, as the former was not available and it was the only species from the order Suliformes in our dataset, so that neither the position in the tree nor the branch length would be affected by this change. The consensus trees were included as *pedigree* in the models. We performed models with either glucose or glycation as dependent variables and with the diet, body mass and life history traits as predicting variables (and glucose in the models with glycation as a response to assess glycation resistance; see introduction; another set of models without glucose were performed to test if there was a covariation of glycation itself, independently of glucose levels, covaried with life-history; see ESM6). Generalized Variance Inflation Factors (GVIFs) were calculated for all the models with more than one predictor variable to assess the collinearity of them, as it may be expected for life history traits (see results on ESM1), considering values above 1.6 as slightly concerning, above 2.2 as moderately concerning and above 3.2 as severely concerning (i.e. indicating high collinearity; following [84]). Models testing the effects of age and sex on glucose and glycation levels and the number of exposed lysine residues on glycation were also carried out. Finally, we performed models on glucose and glycation values controlling for the taxonomic orders included in the dataset (see ESM5). The effects of phylogeny on all models was determined by calculating the ratio of the variance estimated for the “animal” variable, representing the *pedigree* (see ESM6) by the total variance (all random factors plus *units*). A thorough description of all of the models, including transformations of the variables and other details is given on the ESM6.

## Results

### Plasma glucose and albumin glycation variability across the birds’ tree

Plasma glucose and albumin glycation values varied considerably across species (**Figure 1**). Considerable within species repeatability (see ESM6) was observed (Glucose: R = 0.716, SE = 0.042, CI_95_ = [0.619, 0.785], P-value <0.0001; Glycation: R = 0.703, SE = 0.042, CI_95_ = [0.603, 0.768], P-value <0.0001).

**Figure 1.**
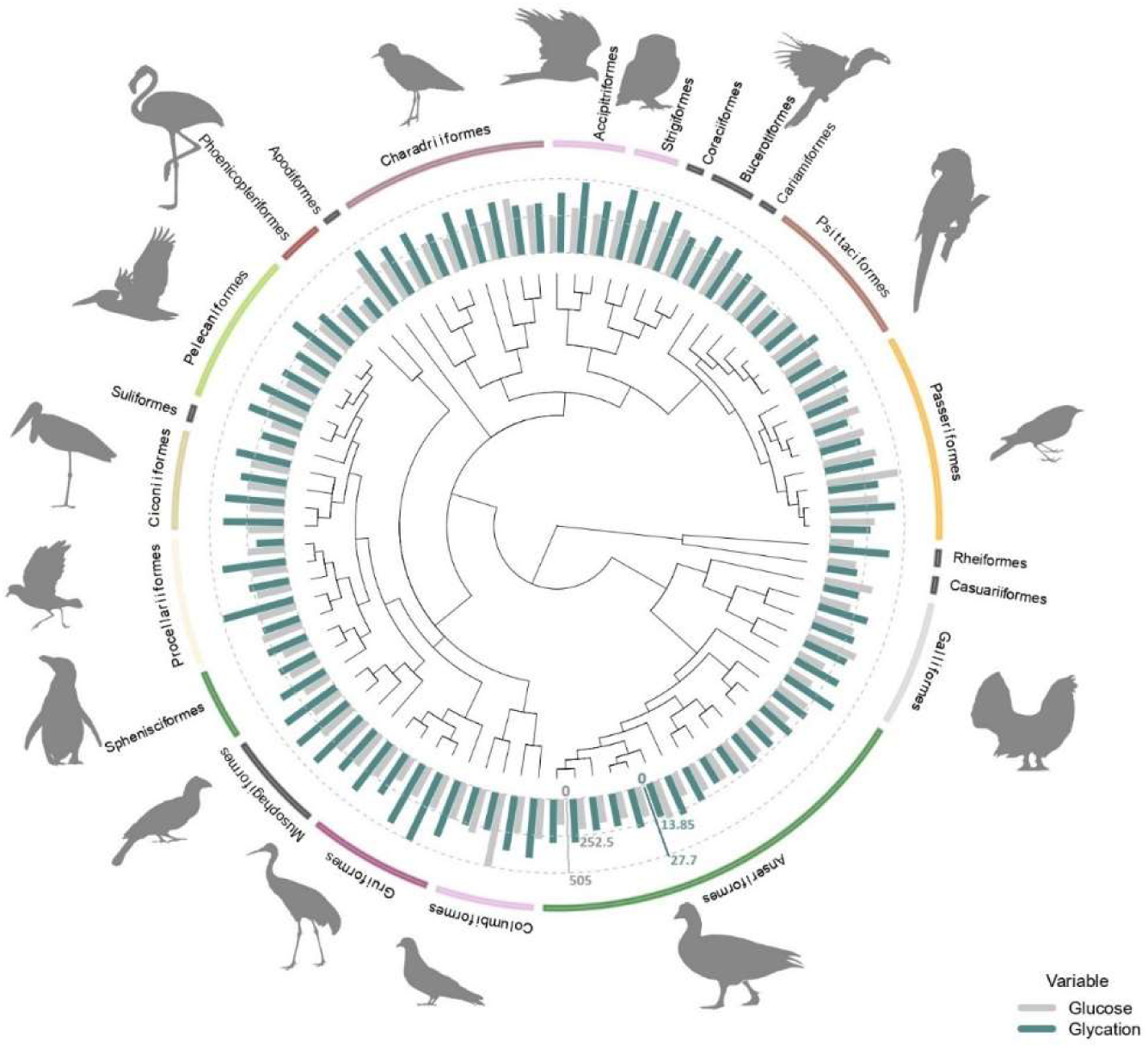
Average plasma glucose values in mg/dL (in grey) and average albumin glycation rate as a percentage of total albumin (in blue) from all the species used in this study (some of them with glucose values coming from ZIMs database; see methods) with the orders they belong to. Glucose and glycation values are standardized in order to be compared, with the dotted lines representing half the maximum and maximum values for each variable (as indicated by the axes in their corresponding colours), from inside out. Tree from a consensus on 10,000 trees obtained from “Hackett All species” on Birdtree.org, including 88 species from 22 orders (see methods).

We found significant differences between some orders for both glucose and albumin glycation. ESM1 shows the raw outcomes of the models, showing such significant differences to the intercept and the Credible Intervals to perform the pairwise comparisons across all groups. To explore some of the details, we can see for example how Apodiformes showed the highest average glucose values (species average model: Estimated mean=351.8 mg/dL, CI_95_[238.9, 514.2]; model with individuals: Estimated mean=349.8 mg/dL, CI_95_[251.8, 476.7], n=39 individuals, 1 species), followed by Passeriformes (species average model: Estimated mean=349.2 mg/dL, CI_95_[279.6, 438.6]; model with individuals: Estimated mean=337.6 mg/dL, CI_95_[274.5, 414.5], n=84 individuals, 8 species), while the Suliformes (species average model: Estimated mean=172 mg/dL, CI_95_[117.6, 252.9]; model with individuals: Estimated mean=169.5 mg/dL, CI_95_[120.9, 239.7], n=5 individuals, 1 species), Rheiformes (species average model: Estimated mean=173.7 mg/dL, CI_95_[118, 254.7]; model with individuals: Estimated mean=172 mg/dL, CI_95_[123.7, 239.8], n=9 individuals, 1 species) and Phoenicopteriformes (species average model: Estimated mean=176 mg/dL, CI_95_[130, 241.1]; model with individuals: Estimated mean=141.2 mg/dL, CI_95_[100.4, 198.2], n=5 individuals, 1 species) show the lowest average values (for the species average models, the pattern is Suliformes<Rheiformes<Phoenicopteriformes, while for the individual models is Phoenicopteriformes<Suliformes<Rheiformes).

For glycation, Strigiformes (species average model: Estimated mean=25.6 %, CI_95_[20, 31]; model with individuals: Estimated mean=25.3 %, CI_95_[19.1, 31.5], n=7 individuals, 2 species), Apodiformes (species average model: Estimated mean=25.5 %, CI_95_[18.6, 32.4]; model with individuals: Estimated mean=24.8 %, CI_95_[17.9, 32], n=40 individuals, 1 species) and Coraciiformes (species average model: Estimated mean=24.5 %, CI_95_[17.9, 31.6]; model with individuals: Estimated mean=23.9 %, CI_95_[15.8, 32], n=2 individuals, 1 species), all terrestrial carnivores as by our sampled species (see discussion below), had the highest average.

On the other hand, Casuariiformes (species average model: Estimated mean=10.8 %, CI_95_[3.7, 18.1]; model with individuals: Estimated mean=11.4 %, CI_95_[2.7, 20.6], n=1 individuals, 1 species), Phoenicopteriformes (species average model: Estimated mean=11.2 %, CI_95_[5.6, 16.7]; model with individuals: Estimated mean=10.5 %, CI_95_[4.6, 16.3], n=22 individuals, 2 species) and Suliformes (species average model: Estimated mean=13.8 %, CI_95_[7, 20.7]; model with individuals: Estimated mean=13.2 %, CI_95_[5.6, 20.5], n=5 individuals, 1 species) had the lowest average levels for the species average models, the pattern is Casuariiformes<Phoenicopteriformes<Suliformes, while for the individual models is Phoenicopteriformes<Casuariiformes<Suliformes). Graphs with raw data on species average and individual glucose and glycation values by order are shown on ESM2 (**Figure ESM2.3**). Estimates of phylogeny effects on the residuals of the models on glucose and glycation values differ if we consider the models with intraspecific variability or without it, but not so much between the models with and without life history traits (within the previous categories). In the case of species averages, the effect of the tree on residual glucose variation is lower and the estimation is less precise than for residual glycation variation, while for the models considering individual values, it is glycation residuals what shows lower levels of tree-related variance than glucose residuals (see ESM1).

### Bird glycemia appears related to body mass and maximum lifespan

After controlling for intraspecific variation in our analyses, we found that variation in glucose levels were significantly explained by variations in body mass and both the linear and quadratic component of residual maximum lifespan (i.e. mass and phylogeny-adjusted maximum lifespan), but not by variations in diet (**Figure 4.A** with predictions from the model without life history traits and **Figure ESM2.2.A** with raw individual data from the same dataset), clutch mass and developmental time (see **Table 1** for the model including life-history trait variables). Heavier species had lower glucose levels (see **Figure 2.A**, drawn with the estimates from the model without life history traits, which includes more species). Glucose levels increase with increasing mass-adjusted lifespan until reaching a plateau (**Figure 2.B**). Models that did not consider intraspecific variability show no significant effect of any of the aforementioned variables on glucose levels (see ESM1). The provenance of the samples (wild versus captive) only showed a trend to a higher glucose levels in the samples from captive individuals in the model without life history traits (Estimate = 0.058; CI_95_[-0.008, 0.125]; P-value=0.083).

**Figure 2.**
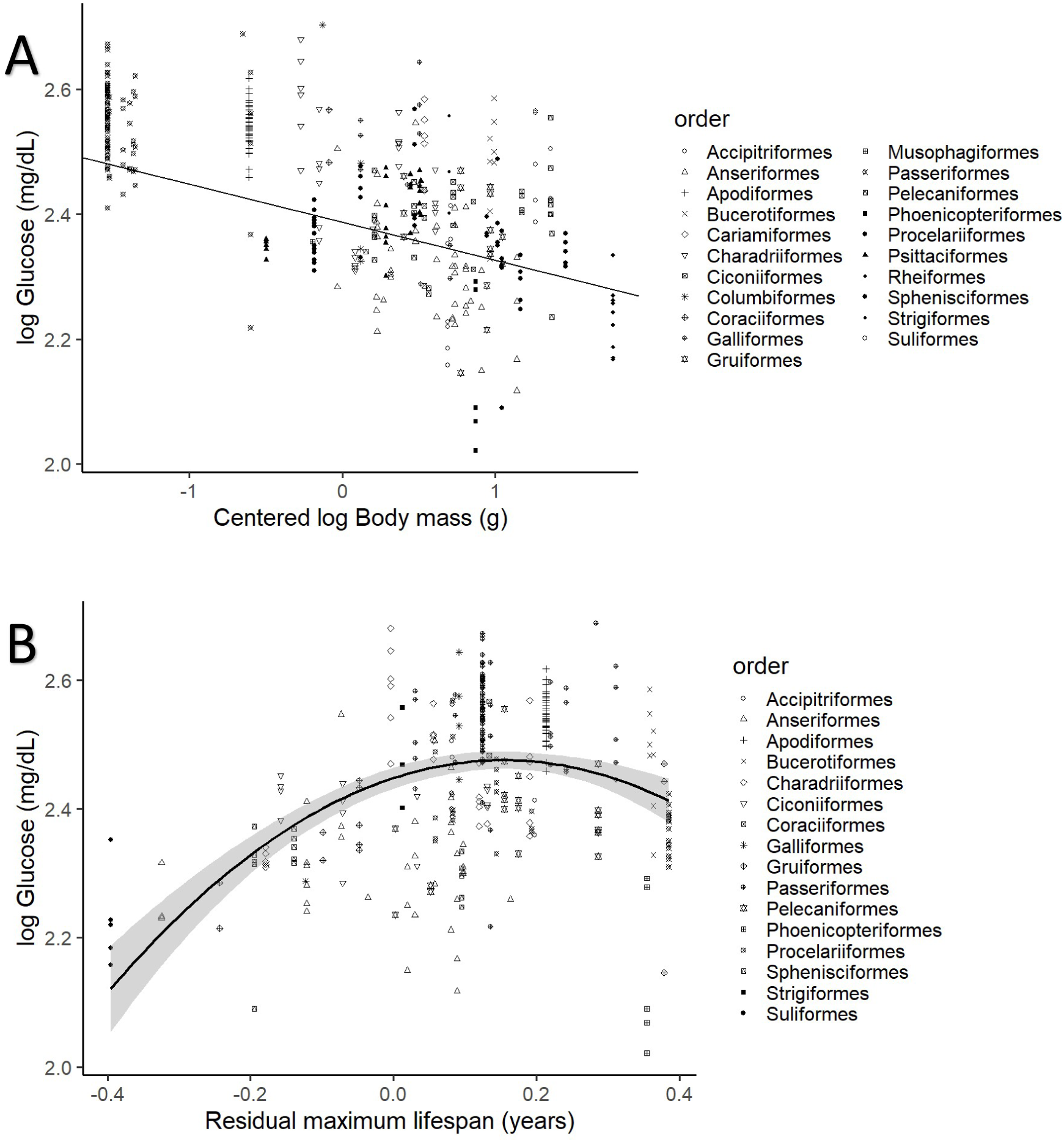
Plasma glucose levels (in mg/dL) variation as a function of **A** species mean centred body mass and **B** residual maximum lifespan. Both glucose and body mass are log transformed. Maximum lifespan (in years) is given as the residues of a pGLS model with body mass (in grams), both log_10_ transformed, so the effects of body mass on longevity are factored out (see ESM6). Different bird orders, are indicated by symbols, as specified on the legends at the right side of the graphs. Figure 2**.A** uses the values and estimates from the glucose model without life history traits (n=389 individuals from 75 species), while Figure 2**.B** uses only the data points employed on the complete model (n=326 individuals of 58 species).

**Table 1.** Final glucose model with diet, body mass, life history traits and sample provenance (wild versus captive; see ESM6) as explanatory variables, including the significant quadratic effect of maximum lifespan. Posterior means, CI_95_ and pMCMC from a phylogenetic MCMC GLMM model including n=326 individuals of 58 species. Both glucose and body mass are log_10_ transformed and life history traits are residuals from a pGLS model of log_10_ body mass and log_10_ of the trait in question (see ESM6). Body mass was also centred to better explain the intercept, as 0 body mass would make no biological sense. The intercept corresponds to the Omnivore diet, being used as the reference as it is considered the most diverse and “neutral” group for this purpose. Significant predictors are indicated in bold.

|  | Estimates | Lower<br>95% CI | Upper<br>95% CI | Sampling<br>effort | pMCMC |
| --- | --- | --- | --- | --- | --- |
| **Intercept (Omnivore)** | **2.387** | **2.268** | **2.5** | **59900** | **<0.001** |
| Diet: Carnivore terrestrial | 0.02 | -0.056 | 0.099 | 59900 | 0.592 |
| Diet: Aquatic predator | 0.034 | -0.047 | 0.112 | 59900 | 0.393 |
| Diet: Herbivore | -0.053 | -0.177 | 0.07 | 59900 | 0.393 |
| Diet: Frugivore/granivore | -0.055 | -0.239 | 0.135 | 59900 | 0.558 |
| **Centred Log<sub>10</sub>Body mass** | **-0.061** | **-0.106** | **-0.015** | **59900** | **0.009** |
| Maximum lifespan | 0.107 | -0.035 | 0.253 | 59900 | 0.142 |
| **Maximum lifespan<sup>2</sup>** | **-0.616** | **-1.166** | **-0.095** | **59900** | **0.026** |
| Clutch mass | -0.095 | -0.265 | 0.069 | 59900 | 0.258 |
| Developmental time | 0.011 | -0.185 | 0.212 | 59900 | 0.916 |
| Provenance: Captive | 0.034 | -0.039 | 0.106 | 59900 | 0.346 |

### Bird albumin glycation is related to glycemia and diet

After controlling for intraspecific variability, we found that diet was affecting variation in albumin glycation rates, with terrestrial carnivorous species having higher glycation levels than omnivorous species (**Table 2** for complete model; see **Figure 4.B** with predictions of the model without life history traits and **Figure ESM2.2.B** with raw individual data from the same dataset). However, for the models for species averages, there was only a trend on this pattern in the one including life history traits, and no effect in the other (see ESM1). The relation between glycation and glucose levels was positive and significant in all but the model that included life history traits but not intraspecific variation (see **Table 2** for the outcome of the model with individual values and life history traits and ESM1 for the rest; see **Figure 3** with estimates from the model including individual variation but no life history traits, as it contains more species and the estimates are similar). Given the logarithmic relationship between glycation and glucose (see ESM6), the slope lower than one (see **Table 2**) implies that birds with higher glucose levels have relatively lower albumin glycation rates for their glucose, fact that we would be referring to as higher glycation resistance. The glycation models excluding glucose levels, and therefore testing for covariates of life-history with glycation itself, without considering resistance, rendered similar results, with only the abovementioned dietary effects being significant (see ESM1).

**Figure 3.**
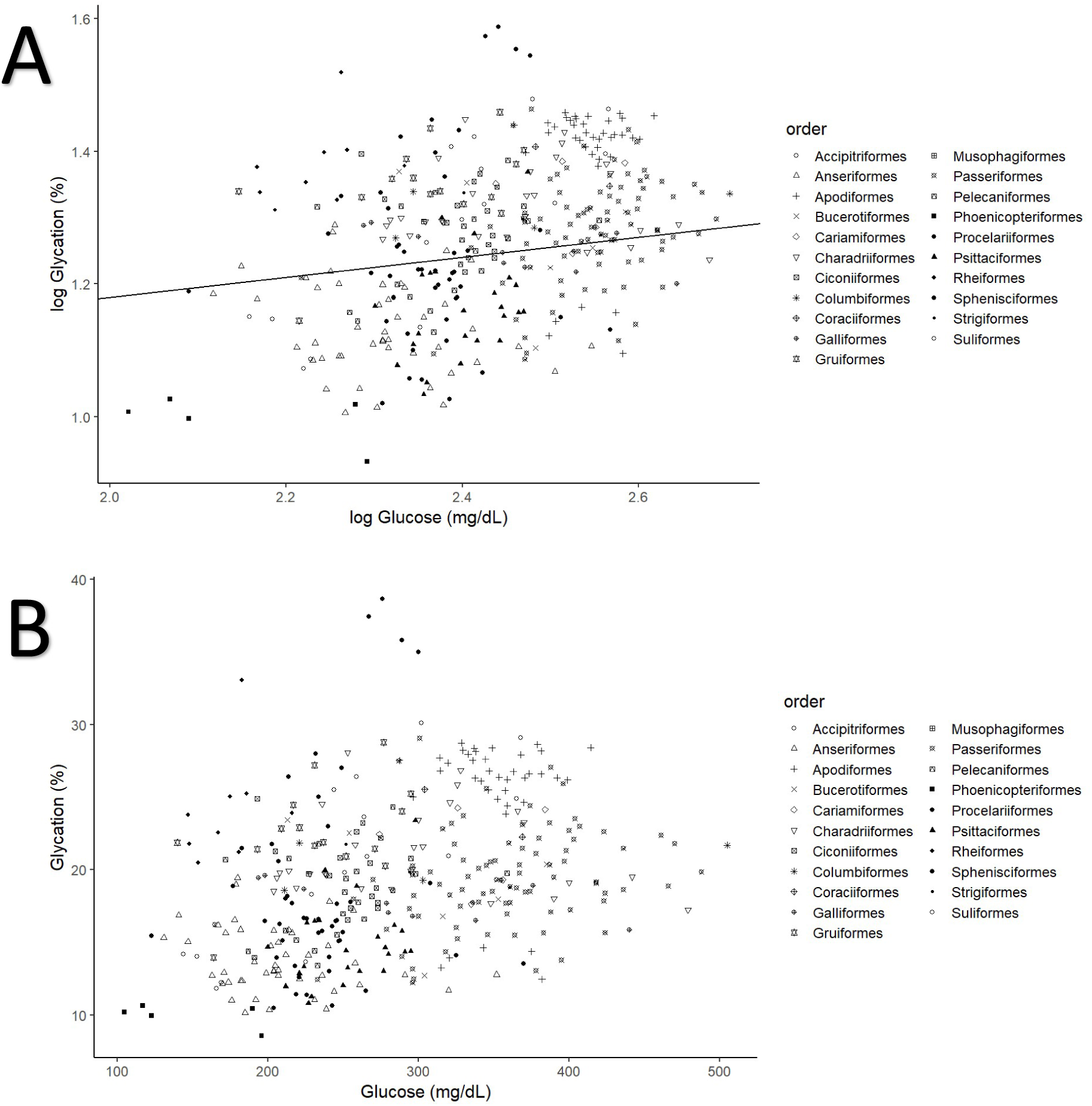
Individual albumin glycation rates (as a percentage of total albumin) variation as a function of individual plasma glucose values (mg/dL). **A**. Both variables log_10_ transformed, as in the model, including the line representing the predicted relationship). **B**. Both variables in a linear form, to more explicitly illustrate the phenomenon referred to as higher albumin glycation resistance in birds with higher plasma glucose levels, inferred from the faster increase in glucose than albumin glycation, i.e. the negative curvature of the relationship. Different bird orders are indicated by symbols, as specified on the legends at the right side of the graphs. The values and estimates used are from the glycation model without life history traits (n=379 individuals from 75 species).

**Figure 4.**
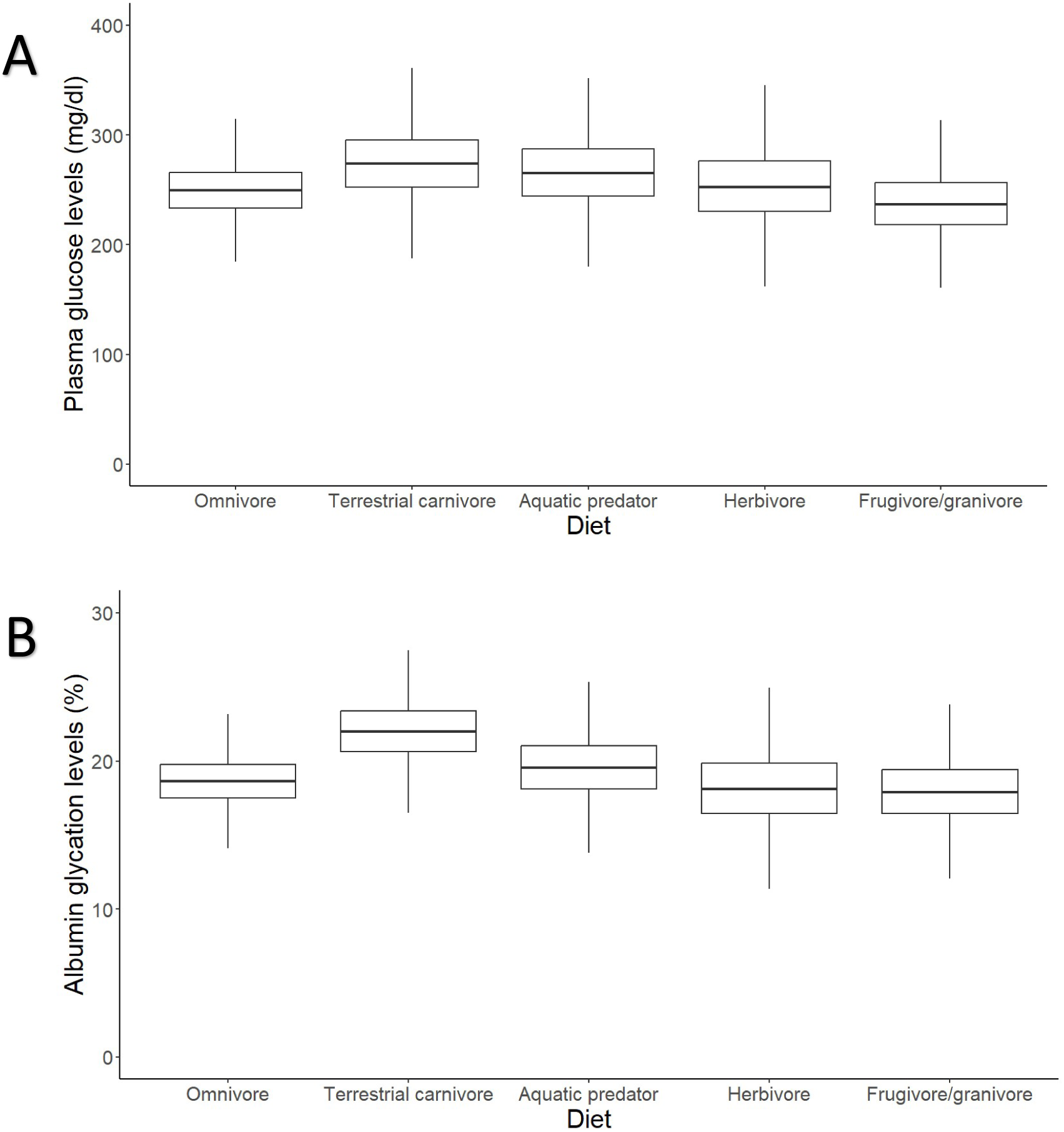
Outcomes of the models (estimates with interquartile ranges from the posterior distributions and whiskers representing credible intervals) on individual data on effects of diet on (**A**) plasma glucose levels and (**B**) albumin glycation in birds. Glucose levels are given in mg/dL, while glycation levels are a percentage of total plasma albumin which is found to be glycated. Terrestrial carnivores showed significantly higher glycation levels than omnivores (Estimate=21.62 %, CI_95_[18, 25.95], p_MCMC_=0.049). Models without life history traits, including more individuals, are represented, but the models with life history traits do not show differences in their qualitative predictions (i.e. higher albumin glycation in terrestrial carnivores than in omnivores; see ESM1).

**Table 2.** Final glycation model with diet, body mass, glucose and life history traits as explanatory variables. Posterior means, CI_95_ and pMCMC from a phylogenetic MCMC GLMM model including n=316 individuals of 58 species. Glycation, glucose and body mass are log_10_ transformed and life history traits are residuals from a linear model of log_10_ body mass and log_10_ of the trait in question (see ESM6). Body mass and glucose were also centred to better explain the intercept. The intercept corresponds to the Omnivore diet, being used as the reference as it is considered the most diverse and “neutral” group for this purpose. Significant predictors are indicated in bold and the credible intervals are considered for making pairwise comparisons between the groups.

|  | Estimates | Lower<br>95% CI | Upper<br>95% CI | Sampling<br>effort | pMCMC |
| --- | --- | --- | --- | --- | --- |
| **Intercept** | **1.232** | **1.112** | **1.351** | **59900** | **<0.001** |
| **Diet: Carnivore terrestrial** | **0.101** | **0.017** | **0.187** | **59900** | **0.021** |
| Diet: Aquatic predator | 0.027 | -0.062 | 0.118 | 59900 | 0.549 |
| Diet: Herbivore | -0.019 | -0.161 | 0.119 | 59936 | 0.781 |
| Diet: Frugivore/granivore | 0.095 | -0.098 | 0.292 | 59900 | 0.329 |
| Centred log<sub>10</sub>Body mass | 0.004 | -0.043 | 0.05 | 58765 | 0.876 |
| **Log<sub>10</sub>Glucose** | **0.137** | **0.012** | **0.255** | **59900** | **0.027** |
| Maximum lifespan | 0.037 | -0.122 | 0.195 | 59043 | 0.648 |
| Clutch mass | 0.151 | -0.03 | 0.346 | 59900 | 0.114 |
| Developmental time | 0.04 | -0.188 | 0.266 | 59303 | 0.725 |

Additional analyses looking at the number of exposed lysines in the albumin sequence of a species show no effect of this variable on albumin glycation rates (α=10.39: CI_95_ [-3.713, 24.993], β=0.246: CI_95_[-0.144, 0.629], P_MCMC_=0.196; ESM1).

We observed no significant effect of age relative to maximum lifespan nor sex on either glycaemia or glycation (see ESM1). In some models, GVIF for body mass and/or clutch mass were higher than 1.6, and in one case body mass is slightly above 2.2 (2.26, see ESM1). This indicates that there may be moderate collinearity for this variables, but the fact that we used life-history trait covariates that are excluding body mass associated variation, this small effects remain mysterious, but nevertheless probably not worrying.

## Discussion

### Plasma glucose level and albumin glycation rate (co)variation patterns suggests resistance mechanisms

Our findings show that glucose levels vary widely among different bird groups. Interspecific differences are partly explained by the allometric relationship of glycaemia with body mass (**Figure 2.A**), which has already been reported in previous studies [56, 63, 34, 13, 14]. Indeed, Passeriformes, Apodiformes and some Columbiformes (e.g. *Nesoenas mayeri* holds the highest value in our dataset) are found at the higher end of the glucose level continuum, in accordance with their relatively small body mass and powered flight, while groups of larger birds such as Phoenicopteriformes, Anseriformes, Rheiformes and Suliformes tend to show low glycaemia levels. This pattern is similar for glycation, with some groups of large birds (such as Phoenicopteriformes, Anseriformes and Suliformes) showing the lowest levels of glycated albumin, while small birds (such as Apodiformes) had the highest values. Nevertheless, glycation remains high in some birds in relation to their glucose levels, as in Rheiformes, or low as in Psittaciformes or Passeriformes. The case of Procellariiformes, which are typically long-lived birds, is particularly striking, with some species exhibiting some of the highest glycation levels (*Calonectris diomedea* and *Macronectes giganteus*) and others some of the lowest (*Procellaria aequinoctialis*). This suggests that, if birds are protected against the deleterious effects of high glycaemia, certain species may have evolved mechanisms to efficiently prevent proteins to be glycated at high rates while others may have evolved mechanisms to resist the consequences of protein glycation. We should also bear in mind that some taxonomic groups may be underrepresented in our study, or biases in species selection due to availability contingencies (e.g. species common in zoos or present in European countries) may exist, so further studies should target this underrepresented groups in order to confirm our predictions.

As expected for a non-enzymatic reaction, just by the law of mass action, bird albumin glycation rates increase with plasma glucose concentration. However, the logarithmic nature of the relationship, and the fact that the slope is lower than one, suggest that species with higher plasma glucose levels exhibit relatively greater resistance to glycation. This finding aligns with previous research indicating that *in vitro* glycation levels of chicken albumin increase at a slower rate than those of human albumin when glucose concentration is elevated [83]. Moreover, these levels are consistently lower than those of bovine serum albumin regardless of glucose concentration and exposition time [37]. As discussed in previous studies comparing chicken to bovine serum albumin [37], or zebra finch to human haemoglobin [53], the lower glycation rates observed in bird proteins may result from a lower number of glycatable amino acids (e.g. lysines) in their sequence and/or their lesser exposure at the protein surface.

Our analyses do not succeed in indicating a significant positive relationship between average glycation levels and the number of glycatable lysines in the albumin sequence. This may be attributed to the limited number of species employed or the weak variation in the number of glycatable lysine residues, which are mostly ranging from 33 to 39. An interesting exception are the 18 glycatable lysine residues of flamingos (*Phoenicopterus ruber*), which also shows very low glycation levels (mean=10.1%). However, the exceptions of 44 in zebra finches and 20 in godwits (*Limosa lapponica*, used in place of *Limosa limosa*) are nevertheless associated with very similar average glycation levels of 20.2% and 20.7% respectively.

### Plasma glucose relates with longevity and may be influenced by reproductive strategies

Our results were only in minor agreement with predictions from POLS theory: it holds for body mass, but not for the other life history variables tested. The only previous comparative study of glycaemia and life history traits to our knowledge [14] shows no relationship with mass-adjusted maximum lifespan in passerines. However, our study over 88 bird species on 22 orders revealed an increase in glucose with mass-adjusted longevity up to a plateau (see **Figure 2.B**). Thus, the relationship between glucose and maximum lifespan may depend on differences between bird orders or be tied to specific species particularities not explored in our study. Such species particularities might involve additional undetermined ecological factors that modify the relationship of glycaemia with longevity. Further exploration of glucose metabolism in relation with lifestyle will bring further light on species-specific life-history adaptations concerning glucose. For example, the species with the lower mass-adjusted maximum lifespan here was a cormorant (*Leucocarbo verrucosus*), which have quite low glucose values for birds.

Regarding reproductive investment (i.e. clutch mass), our results show no relationship with glycation (see **Table 2**), while previous studies reported positive relationships with glycaemia in passerines [14, 83]. Interestingly, most of the species with high clutch mass included in our study belong to the Anseriformes (**Figure ESM2.1**). While these species exhibit very low glycaemia and albumin glycation rates, they are also characterized by a particular reproductive strategy compared to passerines, for whom clutch mass does not imply the same in terms of parental investment. For instance, Anseriformes, unlike Passeriformes, are precocial and their offspring are not very dependent on parental care. Furthermore, they are capital breeders, accumulating the energetic resources required for reproduction in advance rather than collecting them during reproduction. These species typically have large clutch sizes and incubation is usually carried out by females, who store internal fat resources to support this main part of the parental investment. In addition, ducklings are born highly developed, reducing the amount of care required post-hatching (see e.g. [85–86]). Consequently, their dependence on glucose as a rapid energy source for reproduction may be lower, with investment in this activity likely more closely linked to lipid accumulation. This could explain why we did not detect previously reported effects of clutch mass on glucose levels.

### Terrestrial carnivores show a paradoxically increased albumin glycation rate without increased plasma glucose levels

Contrary to our expectation of finding differences across dietary groups, plasma glucose did not significantly vary with species diet. This aligns with results previously reported by [14] for Passeriformes or by [55] for 160 species of vertebrates among which 48 were birds, but not with [15], that showed glycaemia to be increased with the proportion of fruits/seeds in the diet, or with [56], that showed higher (mass-adjusted) glucose levels for terrestrial carnivores and insectivores (those included in our terrestrial carnivores’ category) and lower for frugivorous/nectarivorous. On the other hand, intraspecific data indicates that changes in diet composition hardly affects glycaemia in birds [87]. These studies suggest that glycaemia is tightly regulated independently of dietary composition within species, while it probably varies across species depending on the diet they are adapted to, although depending on the bird groups included in the analyses and the way of assessing it.

Although terrestrial carnivore species did not have significantly higher glycaemia levels in our study, they nonetheless demonstrated significantly higher albumin glycation rates, suggesting a potential susceptibility to protein glycation. Besides unique structural features, this phenomenon could be due to lower albumin turnover rates in terrestrial carnivores. Studies in humans have linked higher oxidative damage to albumin with lower turnover rates (reviewed in [88]), which may extend to other species and post-translational modifications of albumin, such as bird’s albumin glycation. However, the contrary outcome would be anticipated if protein intake were high as seen in carnivorous species [88–89].

Interestingly, the contrast between terrestrial and aquatic predators, which did not show such high glycation rates, suggests the beneficial influence of a substantial difference in food composition or the pre-eminence of another ecological factor, yet to be determined, which may explain lower glycation levels in aquatic predators. In terms of food composition, the proportion of essential n-3 PUFA (polyunsaturated fatty acids), particularly long-chain ones such as DHA and EPA compared with n-6 PUFA and short-chain n-3 PUFA, is different in aquatic and terrestrial environments [90] and therefore in the diet of predators in each environment (e.g. [91]). For instance, low n-6 PUFA/n3-PUFA ratio have been shown to decrease insulin levels and improve insulin resistance in humans [92]. On the other hand, while some studies report that an increase in dietary n3-PUFA reduces the levels of glycated haemoglobin (HbA1c) in humans, this effect was not detected in many other studies and therefore remains inconclusive (reviewed in [93]). The detailed study of the fat composition of the diets of terrestrial and aquatic predators, in particular the ratio between different types of PUFA, merits more attention. In addition, micronutrient content, such as circulating antioxidant defences, should also be taken into consideration. Indeed, a positive relationship between oxidative stress and glycated levels of bovine serum albumin has been reported [94]. One hypothesis is that terrestrial predators have higher systemic oxidative stress levels compared to other species, which may be explained by defects in their antioxidant defences. Uric acid is one of the main non-enzymatic antioxidants in birds [95–97], and uric acid levels are especially high in carnivore species that have a rich protein diet [96, 98–101]. We should therefore take into account other important antioxidants such as vitamin E, vitamin A and carotenoids, which may be less abundant in the diets of terrestrial carnivores (e.g. [102], but see [52]). The question of whether the diet of aquatic carnivores provides a better intake of antioxidants would therefore requires a more detailed description of dietary habits. The herbivorous diet, meanwhile, despite the expected possibility contributing to lower glycaemia and glycation levels due to higher levels of PUFA compared to SFA (saturated fatty acids) (effects reviewed in [93] for humans) and higher fibre content (see [103]), did not lead to significantly lower levels of either of these two parameters. However, the relatively low sample size in that group or the presence of outliers as *Nesoenas mayeri*, with the highest glucose levels of this dataset, makes the interpretation of the obtained results somehow limited. Therefore, further research should be carried out on these species to determine if the expected pattern would emerge with a better sampling.

Finally, differences between captive and wild populations could be considered as a source of variation in glucose or glycation levels, due to a more sedentary lifestyle in captivity, with lower activity levels and a higher food intake likely to lead to increased circulating glucose levels in captive individuals. Additionally, differences in nutrient intake, such as antioxidants, specific fatty acids or amino acids, between captive and wild populations could contribute to variations in glycation levels, as we saw above. Nevertheless, we think that this factor is unlikely to significantly affect our findings regarding diet categories, because our study encompasses species from both captive and wild populations across various diet groups, particularly those exhibiting significant differences (e.g., omnivores versus terrestrial carnivores).

## Conclusion and perspectives

In conclusion, the avian plasma glucose levels measured here are generally higher or in the high range of values recorded for mammalian species [28]. Our study also concludes that there is considerable variation in plasma glucose levels and albumin glycation rates among bird species, with those with the highest glucose levels showing greater resistance to glycation. The correlation between plasma glucose and life history traits are primarily influenced by its inverse association with body mass, along with non-linear, mass-independent effects of longevity. Finally, although diet does not explain plasma glucose levels in our study, terrestrial carnivores have higher albumin glycation rates than omnivores. Whether these intriguing results are explained by specific effects of certain dietary components, such as PUFAs and/or antioxidants, remains to be determined. Differences in plasma glucose levels, albumin glycation rates and glycation resistance (as glycation levels adjusting for plasma glucose concentration) across bird species do not seem consistent with predictions according to the POL hypothesis, except for body mass. Further investigation is needed to elucidate the correlation between these traits and specific life conditions, such as reproductive strategy, migration patterns, flight mode or more detailed diet composition. In addition, more in-depth exploration of glycation levels within high glycaemic groups such as the Passeriformes and other small birds, which make up a significant proportion of avian species, could provide valuable new insights. Similarly, investigating groups such as the Phoenicopteriformes or Anseriformes, which are at the other end of the glycaemic-glycation spectrum, could shed light on the origin of differences between avian orders. Furthermore, notable variations were observed between species within orders, as in the Procellariiformes, which with their high mass-adjusted longevity, and particularly given that some of the age known individuals in our dataset (indeed, two specimens of snow petrels, *Pagodroma nivea*, showed ages higher than the maximum lifespan reported in the sources we explored, determining a new record) have still relatively low levels of glycation, may suggest that some species (the ones that do show higher glycation levels) are not intrinsically resistant to glycation, but rather to its adverse consequences for health. As the main limitations from this study, the usage of individuals from both wild and captive populations, sampled at different periods of the year, and the low sample size for some species, due to logistic constraints, may have introduced noise in the values reported that should be addressed in future studies by implementing a stricter sampling protocol. Also, a more thorough and accurate report and compilation of life history traits from multiple species would allow to increase the number of species included in this kind of analyses. Future research should also focus on specific species to unravel the physiological mechanisms mediating the effects of blood glucose and protein glycation on life-history trade-offs, in particular mechanisms that may vary between taxa and contribute to characteristic adaptations in birds to mitigate the potentially negative effects of comparatively high glycaemia.

## Supporting information

ESM1-4 & 6

## Conflicts of interest

None declared.

## Data and code accessibility

All data used, including references for the values taken from bibliography or databases, will be available as Electronic Supplementary Material (ESM5). Code is made available as Electronic Supplementary Material (ESM7).

## Ethics statement

This study followed all the legal considerations, with the ethic authorisations from the French Ministry of Secondary Education and Research, n°32475 for the zebra finches sampling, the Swiss Veterinary Office (FSVO) n° 34497 for the Alpine swifts, the Ethics Committee of the University of Extremadura (licenses112//2020 and 202//2020) and the Government of Extremadura (licenses CN0012/22/ACA and CN0063/21/ACA) for the godwits and terns from Spain, and Sampling in Terres Australes et Antarctic Françaises was approved by a Regional Animal Experimentation Ethical Committee (French Ministry of Secondary Education and Research permit APAFIS #31773-2019112519421390 v4 and APAFIS#16465–2018080111195526 v4) and by the Comité de l’Environnement Polaire and CNPN (A-2021-55). The samples from the Mulhouse zoo were taken by its licensed veterinary with capacity number: 2020-247-SPAE-162.

## Acknowledgments

This research was funded by an ANR (AGEs – ANR21-CE02-0009). We thank Charles-André Bost (CEBC CNRS) for providing data on some species. We would like to thank the SYLATR Association for collecting samples of wild Passeriformes as part of the MIGROUILLE program, and Manuela Forero and Frederic Angelier (CEBC CNRS) for providing some samples of certain Procellariiformes. We thank the French Parc des Oiseaux (Villars les Dombes), the bird keepers and veterinarians at Mulhouse zoo for their contribution to the collection of blood samples and access to individual data from their captive birds. Data on seabirds from the French Southern Territories was collected within the framework of the ECONERGY (119), ECOPATH (1151) and ORNITHOECO (109) programs of the French Polar Institute (IPEV). These studies are part of the long-term Studies in Ecology and Evolution (SEE-Life) program of the CNRS. We are grateful to Mathilde Lejeune, Natacha Garcin and Camille Lemonnier for their help in collecting those samples. We are thankful to Orsolya Vincze for producing Figure 1, Adrien Brown for determining the number of lysines exposed in albumin sequences, Claire Saraux for her help with the statistics and F. Stephen Dobson for reading and commenting on the manuscript. Finally, we are thankful to Ascensia Diabetes care® for their generous donation of glucometers and strips for glucose measurement.

## Authors’ contributions

. Criscuolo and F. Bertile conceived the idea, directed most of the sample collection and logistics and contributed significantly to the writing. A. Moreno-Borrallo contributed to the development of the questions, gathered the diet and life-history data, performed the statistical analyses, and led the writing. S. Jaramillo Ortiz and C. Schaeffer performed the mass spectrometry analyses for protein glycation measurements, and S. Jaramillo Ortiz contributed to the glucose measures and commented on the manuscript. T. Boulinier. and V. A. Viblanc coordinated the ECONERGY and ECOPATH polar programs, organised the collection of samples on subantarctic seabirds, and commented on the manuscript. Olivier Chastel collaborated collecting marine bird samples and commented on the manuscript. B. Rey contributed with the collection of samples from Parc des Oiseaux (Villars-les-Dombes, France) and commented on the manuscript. P. Bize leads the monitoring of Alpine swifts’ populations from which the samples were obtained, he helped collecting them and commented on the manuscript. J. S Gutiérrez and J. A. Masero contributed with samples from Spain and part of the statistic scripts and commented on the manuscript. B. Quintard leads the health monitoring of Mulhouse zoo bird collection, organized and realized most of Mulhouse zoo samplings and commented on the manuscript. All authors gave their approval for publication.

## Notes

### Competing Interest Statement

The authors have declared no competing interest.

### Summary of Updates

A model and a few little corrections have been added following reviewers' recommendations.

