## Supplementary material for "Variation in albumin glycation rates in birds suggests resistance to relative hyperglycaemia rather than conformity to the pace of life syndrome hypothesis": ESM1-4 & 6: ESM1.pdf

### Electronic Supplementary Material 1 (ESM1)

#### Models with species averages

##### Glucose model

```
 Iterations = 10001:5999901
 Thinning interval  = 100
 Sample size  = 59900 

 DIC: -106.6529 

 G-structure:  ~animal

       post.mean l-95% CI u-95% CI eff.samp
animal   0.03626 0.000596   0.1426    25802

               ~Method_Glu

           post.mean l-95% CI u-95% CI eff.samp
Method_Glu    0.8756 0.000616   0.9388    59900

 R-structure:  ~units

      post.mean  l-95% CI u-95% CI eff.samp
units   0.02191 0.0006759  0.07767    48120

 Location effects: logGlucose ~ Diet + Centered_logBM + Procedence 

                          post.mean l-95% CI u-95% CI eff.samp  pMCMC  
(Intercept)                 2.37100  0.82502  3.98545    59071 0.0103 
*
DietTerrestrial_carnivore   0.02034 -1.05933  1.14510    59900 0.9732  
DietAquatic_predator        0.01828 -1.37382  1.41625    59149 0.9776  
DietHerbivore              -0.02100 -1.17104  1.13783    59900 0.9680  
DietFrugivore_granivore    -0.04185 -1.39315  1.24717    59001 0.9539  
Centered_logBM             -0.11365 -0.84646  0.64960    59130 0.7698  
ProcedenceCaptive           0.03627 -1.29070  1.40114    59900 0.9589  
---
Signif. codes:  0 ‘***’ 0.001 ‘**’ 0.01 ‘*’ 0.05 ‘.’ 0.1 ‘ ’ 1
```

##### Glucose model with life history traits

```
Iterations = 10001:5999901
 Thinning interval  = 100
 Sample size  = 59900 

 DIC: -74.58572 

 G-structure:  ~animal

       post.mean  l-95% CI u-95% CI eff.samp
animal   0.05226 0.0006624   0.2157    29869

               ~Method_Glu

           post.mean  l-95% CI u-95% CI eff.samp
Method_Glu     1.268 0.0006281    1.323    59900

 R-structure:  ~units

      post.mean  l-95% CI u-95% CI eff.samp
units   0.02908 0.0007187   0.1087    51386
```

```
 Location effects: logGlucose ~ Diet + Centered_logBM + poly(ML, 2, ra
w = TRUE) + CM + DT + Procedence

                           post.mean  l-95% CI u-95% CI eff.samp pMCMC
(Intercept)                 2.334820  0.399411 4.337178 59900 0.0257 *
DietTerrestrial_carnivore  -0.004026 -1.528592 1.541982 59900 0.9949
DietAquatic_predator       -0.010700 -1.596257 1.640390 60642 0.9921
DietHerbivore              -0.036708 -1.761151 1.630230 58817 0.9657
DietFrugivore_granivore    -0.030716 -7.794374 7.477946 59900 0.9931
Centered_logBM             -0.078237 -1.000564 0.890303 56515 0.8719
poly(ML, 2, raw = TRUE)1    0.176471 -3.238775 3.599515 60759 0.9254
poly(ML, 2, raw = TRUE)2  -0.393328 -14.806955 14.379736 59900 0.9541
CM                         -0.168559 -3.204873 2.815380 59967 0.9111
DT                         -0.040236 -4.049867 4.067674 59900 0.9832
ProcedenceCaptive           0.031314 -1.430862 1.537092 59900 0.9665
---
Signif. codes:  0 ‘***’ 0.001 ‘**’ 0.01 ‘*’ 0.05 ‘.’ 0.1 ‘ ’ 1
```

#### Glycation model

```
Iterations = 10001:5999901
 Thinning interval  = 100
 Sample size  = 59900

 DIC: -106.7

 G-structure:  ~animal

       post.mean  l-95% CI u-95% CI eff.samp
animal   0.03487 0.0007631   0.1392    25819

               ~Method_Glu

           post.mean  l-95% CI u-95% CI eff.samp
Method_Glu    0.3444 0.0005727   0.3467    59900

 R-structure:  ~units

      post.mean  l-95% CI u-95% CI eff.samp
units   0.02111 0.0007395  0.07448    49146

 Location effects: logGlycation ~ Diet + Centered_logBM + Centered_log
Glucose

                          post.mean  l-95% CI  u-95% CI eff.samp pMCMC
(Intercept)                1.255901  0.344806  2.134884    59900 0.012
*
DietTerrestrial_carnivore  0.052502 -1.044091  1.185074    59900 0.925
DietAquatic_predator       0.018310 -1.290381  1.300686    59900 0.982
DietHerbivore             -0.053868 -1.405728  1.257560    58559 0.937
DietFrugivore_granivore   -0.018507 -1.102124  1.105271    59900 0.974
Centered_logBM             0.008006 -0.696443  0.699297    61227 0.979
Centered_logGlucose        0.339051 -4.654194  5.325153    59900 0.893
---
Signif. codes:  0 ‘***’ 0.001 ‘**’ 0.01 ‘*’ 0.05 ‘.’ 0.1 ‘ ’ 1
```

#### Glycation model with life history traits

```
Iterations = 10001:5999901
 Thinning interval  = 100
 Sample size  = 59900

 DIC: -75.40128

 G-structure:  ~animal
```

```
       post.mean  l-95% CI u-95% CI eff.samp
animal   0.04792 0.0005362   0.1982    32101

               ~Method_Glu

           post.mean  l-95% CI u-95% CI eff.samp
Method_Glu    0.2793 0.0005303   0.4877    59900

 R-structure:  ~units

      post.mean  l-95% CI u-95% CI eff.samp
units   0.02719 0.0007491   0.1021    52282

 Location effects: logGlycation ~ Diet + Centered_logBM + Centered_log
Glucose + ML + CM + DT

                          post.mean l-95% CI  u-95% CI eff.samp pMCMC
(Intercept)                1.227159 -0.058905  2.496171  59900 0.0595 .
DietTerrestrial_carnivore 0.077119 -1.379997  1.520340  59900 0.9173
DietAquatic_predator       0.008690 -1.495711  1.518194  59900 0.9930
DietHerbivore             -0.052138 -1.807832  1.645961  61691 0.9542
DietFrugivore_granivore  -0.078247 -2.194939  1.920607  59900 0.9352
Centered_logBM             0.016661 -0.841361  0.858227  59900 0.9681
Centered_logGlucose        0.179413 -6.579065  6.883513  59900 0.9562
ML                         0.002993 -3.083707  3.134078  60638 0.9977
CM                        -0.093355 -2.629818  2.406782  59900 0.9330
DT                        -0.130439 -4.199502  3.852793  59900 0.9494
---
Signif. codes:  0 ‘***’ 0.001 ‘**’ 0.01 ‘*’ 0.05 ‘.’ 0.1 ‘ ’ 1
```

#### Glycation model with life history traits without glucose

```
Iterations = 10001:5999901
 Thinning interval  = 100
 Sample size  = 59900

 DIC: -75.51184

 G-structure:  ~animal

       post.mean  l-95% CI u-95% CI eff.samp
animal     0.045 0.0007373   0.1844    31737

 R-structure:  ~units

      post.mean  l-95% CI u-95% CI eff.samp
units   0.02685 0.0006839   0.1004    50556

 Location effects: logGlycation ~ Diet + Centered_logBM + ML + CM + DT

                          post.mean l-95% CI  u-95% CI eff.samp pMCMC
(Intercept)                1.217639  0.054541  2.310440 60601 0.0346 *
DietTerrestrial_carnivore 0.083666 -1.321588  1.489212 59900 0.9051
DietAquatic_predator       0.004772 -1.469288  1.512990 61482 0.9914
DietHerbivore             -0.059856 -1.696896  1.572570 59900 0.9431
DietFrugivore_granivore  -0.074390 -2.035350  1.894847 59900 0.9432
Centered_logBM             0.007357 -0.732469  0.739157 59900 0.9796
ML                         0.026304 -2.905055  3.003113 59900 0.9877
CM                        -0.108279 -2.658967  2.356612 59900 0.9308
DT                        -0.132416 -4.002316  3.945679 59900 0.9465
```

### Models with individuals

#### Glucose model

```
Iterations = 10001:5999901
 Thinning interval  = 100
 Sample size  = 59900

 DIC: -992.896

 G-structure:  ~animal

       post.mean l-95% CI u-95% CI eff.samp
animal  0.008021 0.002293  0.01494    59597

               ~species

        post.mean l-95% CI u-95% CI eff.samp
species  0.002995 0.001171 0.005079    59900

 R-structure:  ~units

      post.mean l-95% CI u-95% CI eff.samp
units  0.003922 0.003325 0.004533    59900

 Location effects: logGlucose ~ Diet + Centered_logBM + Procedence

                       post.mean l-95% CI  u-95% CI eff.samp pMCMC
(Intercept)               2.358384 2.252259 2.459482 60833 < 2e-05 ***
DietTerrestrial_carnivore 0.037311 -0.038582 0.111453 59900 0.32174
DietAquatic_predator      0.042473 -0.041770 0.127272 59900 0.31720
DietHerbivore             0.002547 -0.101022 0.101821 59900 0.96551
DietFrugivore_granivore  -0.053312 -0.132924 0.024621 59900 0.17843
Centered_logBM          -0.066964 -0.105964 -0.028517 59148 0.00134 **
ProcedenceCaptive         0.057926 -0.007559 0.124586 59900 0.08344 .
---
Signif. codes:  0 ‘***’ 0.001 ‘**’ 0.01 ‘*’ 0.05 ‘.’ 0.1 ‘ ’ 1
```

#### Glucose model with life history traits

```
Iterations = 10001:5999901
 Thinning interval  = 100
 Sample size  = 59900

 DIC: -822.3621

 G-structure:  ~animal

       post.mean l-95% CI u-95% CI eff.samp
animal  0.008097 0.001737  0.01628    59900

               ~species

        post.mean l-95% CI u-95% CI eff.samp
species  0.003087 0.001147 0.005393    59900

 R-structure:  ~units

      post.mean l-95% CI u-95% CI eff.samp
units  0.004091 0.003425   0.0048    59900

 Location effects: logGlucose ~ Diet + Centered_logBM + poly(ML, 2, ra
w = TRUE) + CM + DT + Procedence

                        post.mean l-95% CI u-95% CI eff.samp pMCMC
```

```
(Intercept)                  2.38663  2.26809  2.50047  59900 < 2e-05 ***
DietTerrestrial_carnivore 0.02035 -0.05557  0.09867  59900 0.59205
DietAquatic_predator       0.03412 -0.04719  0.11187  59900 0.39309
DietHerbivore             -0.05276 -0.17692  0.07030  59900 0.39309
DietFrugivore_granivore   -0.05527 -0.23923  0.13503  59900 0.55790
Centered_logBM            -0.06103 -0.10638 -0.01516  59900 0.00908 **
poly(ML, 2, raw = TRUE)1   0.10739 -0.03536  0.25323  59900 0.14227
poly(ML, 2, raw = TRUE)2 -0.61582 -1.16638 -0.09541  59900 0.02611 *
CM                        -0.09497 -0.26479  0.06912  59900 0.25823
DT                         0.01082 -0.18470  0.21196  59900 0.91606
ProcedenceCaptive          0.03443 -0.03902  0.10580  59900 0.34648
---
Signif. codes:  0 ‘***’ 0.001 ‘**’ 0.01 ‘*’ 0.05 ‘.’ 0.1 ‘ ’ 1
```

#### Glycation model

```
Iterations = 10001:5999901
 Thinning interval  = 100
 Sample size  = 59900

 DIC: -933.1616

 G-structure:  ~animal

       post.mean l-95% CI u-95% CI eff.samp
animal  0.007986 0.001979  0.01522    59900

              ~species

        post.mean l-95% CI u-95% CI eff.samp
species  0.004466  0.00202 0.007134    59900

 R-structure:  ~units

      post.mean l-95% CI u-95% CI eff.samp
units  0.004241   0.0036 0.004937    59900

 Location effects: logGlycation ~ Diet + Centered_logBM + Centered_log
Glucose

                        post.mean  l-95% CI  u-95% CI eff.samp pMCMC
(Intercept)                1.255213 1.161645 1.356336 59900 < 2e-05 ***
DietTerrestrial_carnivore 0.079586  0.000112 0.158918 59900 0.04938 *
DietAquatic_predator       0.021792 -0.064410 0.111384 59900 0.62651
DietHerbivore              0.005237 -0.104032 0.113390 59900 0.92511
DietFrugivore_granivore   -0.010220 -0.090716 0.066216 59900 0.79736
Centered_logBM             0.003794 -0.035404 0.041457 59900 0.84197
Centered_logGlucose        0.150789  0.042859 0.258714 59900 0.00621 **
---
Signif. codes:  0 ‘***’ 0.001 ‘**’ 0.01 ‘*’ 0.05 ‘.’ 0.1 ‘ ’ 1
```

#### Glycation model with life history traits

```
Iterations = 10001:5999901
 Thinning interval  = 100
 Sample size  = 59900

 DIC: -757.4936

 G-structure:  ~animal

       post.mean l-95% CI u-95% CI eff.samp
animal  0.009425 0.001817  0.01963    59900

              ~species

        post.mean l-95% CI u-95% CI eff.samp
```

```
species  0.005169 0.002237 0.008639    59900

 R-structure:  ~units

      post.mean l-95% CI u-95% CI eff.samp
units  0.004572 0.003795  0.00536    59900

 Location effects: logGlycation ~ Diet + Centered_logBM + Centered_log
Glucose + ML + CM + DT

                         post.mean l-95% CI u-95% CI eff.samp pMCMC
(Intercept)               1.231839  1.111789 1.351048 59900 <2e-05 ***
DietTerrestrial_carnivore 0.100920  0.017273 0.187125 59900 0.0214 *
DietAquatic_predator      0.026869 -0.062448 0.117517 59900 0.5491
DietHerbivore            -0.018824 -0.160894 0.119239 59936 0.7812
DietFrugivore_granivore   0.095183 -0.097946 0.291806 59900 0.3288
Centered_logBM            0.003664 -0.043033 0.050278 58765 0.8760
Centered_logGlucose       0.137154  0.011637 0.255134 59900 0.0273 *
ML                        0.036577 -0.121966 0.195499 59043 0.6482
CM                        0.150587 -0.030387 0.346258 59900 0.1139
DT                        0.039982 -0.187558 0.265946 59303 0.7248
```

**Glycation model with life history traits without glucose**

```
Iterations = 10001:5999901
 Thinning interval  = 100
 Sample size  = 59900

 DIC: -756.3145

 G-structure:  ~animal

       post.mean l-95% CI u-95% CI eff.samp
animal    0.0105 0.001971  0.02166    59900

               ~species

        post.mean l-95% CI u-95% CI eff.samp
species  0.005329 0.002197 0.008948    59900

 R-structure:  ~units

      post.mean l-95% CI u-95% CI eff.samp
units    0.0046 0.003839  0.00541    59186

 Location effects: logGlycation ~ Diet + Centered_logBM + ML + CM + DT

                         post.mean l-95% CI u-95% CI eff.samp pMCMC
(Intercept)               1.227931  1.106099 1.357043 59900 <2e-05 ***
DietTerrestrial_carnivore 0.106991  0.018900 0.192773 60415 0.0162 *
DietAquatic_predator      0.031684 -0.058239 0.127370 59900 0.4962
DietHerbivore            -0.028291 -0.171719 0.118113 59900 0.6928
DietFrugivore_granivore   0.098888 -0.100352 0.297249 59900 0.3198
Centered_logBM           -0.003569 -0.051527 0.044211 59900 0.8783
ML                        0.048297 -0.113219 0.208916 58878 0.5488
CM                        0.153343 -0.038424 0.347077 60758 0.1137
DT                        0.042588 -0.190902 0.276599 59900 0.7191
```

### Age & Sex models

#### Glucose

```
 Iterations = 10001:5999901
 Thinning interval  = 100
 Sample size  = 59900

 DIC: -635.2492
```

```
 G-structure:  ~animal

       post.mean l-95% CI u-95% CI eff.samp
animal  0.008703 0.001214  0.01909    59900

               ~species

        post.mean  l-95% CI u-95% CI eff.samp
species  0.002923 0.0006328 0.005658    59900

 R-structure:  ~units

      post.mean l-95% CI u-95% CI eff.samp
units  0.003799 0.003072 0.004595    59900

 Location effects: logGlucose ~ Centered_logBM + poly(logit(Age_relati
ve, FALSE), 2, raw = TRUE) + Sex

                                                 post.mean  l-95% CI
u-95% CI
(Intercept)                                       2.391844  2.278127
2.506021
Centered_logBM                                   -0.052561 -0.097117
-0.006228
poly(logit(Age_relative, FALSE), 2, raw = TRUE)1  0.001298 -0.013674
0.016000
poly(logit(Age_relative, FALSE), 2, raw = TRUE)2 -0.001157 -0.004889
0.002654
SexM                                              0.005705 -0.012474
0.023691
                                                 eff.samp  pMCMC
(Intercept)                                         59186 <2e-05 ***
Centered_logBM                                      59101 0.0275 *
poly(logit(Age_relative, FALSE), 2, raw = TRUE)1    59900 0.8645
poly(logit(Age_relative, FALSE), 2, raw = TRUE)2    59900 0.5414
SexM                                                60869 0.5336
---
Signif. codes:  0 ‘***’ 0.001 ‘**’ 0.01 ‘*’ 0.05 ‘.’ 0.1 ‘ ’ 1
```

#### Glycation

```
 Iterations = 10001:5999901
 Thinning interval  = 100
 Sample size  = 59900

 DIC: 1145.016

 G-structure:  ~animal

       post.mean l-95% CI u-95% CI eff.samp
animal      19.3    3.365    38.51    59900

               ~species

        post.mean l-95% CI u-95% CI eff.samp
species      5.82    1.536    10.92    59900

 R-structure:  ~units

      post.mean l-95% CI u-95% CI eff.samp
units     6.026    4.861    7.266    59742

 Location effects: Glycation ~ Centered_logBM + Centered_logGlucose +
poly(logit(Age_relative, FALSE), 2, raw = TRUE) + Sex

                                                 post.mean l-95% CI u-
95% CI eff.samp
```

```
(Intercept)                                      19.26559 14.02977 24
.55452    59900
Centered_logBM                                    1.12330 -0.88744  3
.12563    63546
Centered_logGlucose                               9.74032  4.27174 15
.04954    59900
poly(logit(Age_relative, FALSE), 2, raw = TRUE)1  -0.07541 -0.66195  0
.53149    59900
poly(logit(Age_relative, FALSE), 2, raw = TRUE)2  -0.05413 -0.20650  0
.09595    60681
SexM                                              0.44836 -0.28507  1
.20100    59900
                                                   pMCMC
(Intercept)                                      < 2e-05 ***
Centered_logBM                                   0.269850
Centered_logGlucose                              0.000267 ***
poly(logit(Age_relative, FALSE), 2, raw = TRUE)1 0.806010
poly(logit(Age_relative, FALSE), 2, raw = TRUE)2 0.482771
SexM                                             0.233790
---
Signif. codes:  0 ‘***’ 0.001 ‘**’ 0.01 ‘*’ 0.05 ‘.’ 0.1 ‘ ’ 1
```

### Lysines model

```
Iterations = 10001:5999951
 Thinning interval  = 50
 Sample size  = 119800

 DIC: 111.8866

 G-structure:  ~animal

       post.mean l-95% CI u-95% CI eff.samp
animal     14.21   0.5989    45.07   111860

 R-structure:  ~units

      post.mean l-95% CI u-95% CI eff.samp
units     18.31    4.944    35.13   116731

 Location effects: Glycation ~ Lysines

            post.mean l-95% CI u-95% CI eff.samp pMCMC
(Intercept)   10.3914  -3.7127  24.9929   119800 0.144
Lysines        0.2462  -0.1438   0.6290   119800 0.196
```

### Orders models

#### Glucose averages

```
 Iterations = 10001:5999901
 Thinning interval  = 100
 Sample size  = 59900

 DIC: -209.2166

 G-structure:  ~species

        post.mean  l-95% CI u-95% CI eff.samp
species  0.001705 0.0003464 0.003511    59900

               ~Method_Glu

           post.mean  l-95% CI u-95% CI eff.samp
Method_Glu  0.004672 0.0001915  0.01309    59900
```

```
 R-structure:  ~units

      post.mean l-95% CI u-95% CI eff.samp
units  0.003713 0.001608 0.005851    59900

 Location effects: logGlucose ~ order 

                        post.mean  l-95% CI  u-95% CI eff.samp pMCMC    
(Intercept)               2.442097  2.317939 2.562344 59900 <2e-05 ***
orderAnseriformes       -0.136101 -0.229771 -0.045736 59900 0.00367 ** 
orderApodiformes          0.104208 -0.063956  0.269026 59900 0.21910    
orderBucerotiformes       0.045225 -0.085697  0.177252 59900 0.49646    
orderCariamiformes        0.084108 -0.078716  0.255211 59900 0.31706    
orderCasuariiformes      -0.163069 -0.337731  0.011609 60636 0.06614 .  
orderCharadriiformes      0.005304 -0.095223  0.101372 59900 0.91780    
orderCiconiiformes       -0.039864 -0.150798  0.069105 59900 0.47349    
orderColumbiformes        0.062913 -0.047089  0.173563 59900 0.26331    
orderCoraciiformes        0.086470 -0.080365  0.253869 59900 0.30067    
orderGalliformes         -0.014638 -0.122414  0.088507 59900 0.78227    
orderGruiformes          -0.086754 -0.191388  0.019906 60809 0.10554    
orderMusophagiformes     -0.032880 -0.149817  0.083853 59900 0.57723    
orderPasseriformes        0.100976  0.004481  0.199953 59900 0.04374 *  
orderPelecaniformes      -0.078898 -0.182340  0.022862 59900 0.12725    
orderPhoenicopteriformes -0.196479 -0.328292 -0.059938 59900 0.00491 *
*
orderProcellariiformes   -0.033725 -0.138918  0.071199 59900 0.52568    
orderPsittaciformes      -0.033701 -0.136006  0.070392 59900 0.51730    
orderRheiformes          -0.202339 -0.370220 -0.036105 59900 0.01930 *  
orderSphenisciformes     -0.131626 -0.249327 -0.012622 58358 0.02945 *  
orderStrigiformes         0.058533 -0.073821  0.193863 59900 0.38818    
orderSuliformes          -0.206606 -0.371552 -0.039227 60514 0.01603 *  
---
Signif. codes:  0 ‘***’ 0.001 ‘**’ 0.01 ‘*’ 0.05 ‘.’ 0.1 ‘ ’ 1
```

#### Glycation averages

```
 Iterations = 10001:5999901
 Thinning interval  = 100
 Sample size  = 59900 

 DIC: 444.2672 

 G-structure:  ~species

        post.mean l-95% CI u-95% CI eff.samp
species     2.873   0.6036    5.908    59900

               ~Method_Glu

           post.mean l-95% CI u-95% CI eff.samp
Method_Glu     8.967   0.3913    25.25    59900

 R-structure:  ~units

      post.mean l-95% CI u-95% CI eff.samp
units     6.234    2.745    9.778    59900

 Location effects: Glycation ~ order 

                        post.mean l-95% CI u-95% CI eff.samp pMCMC    
(Intercept)                23.8174  18.6864 29.0157 59085 0.000267 ***
orderAnseriformes          -9.0998 -12.8132 -5.3161 59900  < 2e-05 ***
orderApodiformes            1.6941  -5.1762  8.5942 59900 0.625309    
orderBucerotiformes        -3.8560  -9.1864  1.6821 59900 0.161536    
orderCariamiformes         -1.6412  -8.5153  5.1824 59900 0.638765    
orderCasuariiformes       -13.0636 -20.0693 -5.6892 59900 0.000401 ***
orderCharadriiformes       -3.4750  -7.4881  0.5848 59900 0.091386 .  
orderCiconiiformes         -2.3910  -6.9647  2.0819 60604 0.299299    
orderColumbiformes         -3.5363  -8.0847  1.0178 59900 0.127579    
```

```
orderCoraciiformes          0.7004  -5.8813  7.7817 59900 0.841002
orderGalliformes           -5.2965  -9.6262 -0.9362 59900 0.018030 *
orderGruiformes            -0.2166  -4.5451  4.1320 58975 0.917262
orderMusophagiformes       -1.3584  -6.2273  3.3938 58473 0.573823
orderPasseriformes         -4.5115  -8.4707 -0.4609 59900 0.028815 *
orderPelecaniformes        -4.9655  -9.1079 -0.7669 59900 0.020568 *
orderPhoenicopteriformes  -12.6524 -18.1743 -7.1537 59900  < 2e-05 ***
orderProcellariiformes     -3.5527  -7.7734  0.8770 59900 0.105275
orderPsittaciformes        -8.4290 -12.7042 -4.2682 59900 0.000134 ***
orderRheiformes             0.5901  -6.4606  7.2461 59900 0.868381
orderSphenisciformes       -6.4491 -11.4353 -1.7623 59900 0.009683 **
orderStrigiformes           1.7468  -3.7739  7.2189 59900 0.528013
orderSuliformes           -10.0020 -16.8119 -3.1082 59900 0.005376 **
---
Signif. codes:  0 ‘***’ 0.001 ‘**’ 0.01 ‘*’ 0.05 ‘.’ 0.1 ‘ ’
```

#### Glucose individuals

```
 Iterations = 10001:5999901
 Thinning interval  = 100
 Sample size  = 59900

 DIC: -989.3556

 G-structure:  ~species

        post.mean l-95% CI u-95% CI eff.samp
species   0.00332  0.00165 0.005173    59900

 R-structure:  ~units

      post.mean l-95% CI u-95% CI eff.samp
units  0.003992 0.003375 0.004628    59900

 Location effects: logGlucose ~ order

                          post.mean  l-95% CI  u-95% CI eff.samp pMCMC
(Intercept)                2.441807  2.363471 2.518260 59900 <2e-05 ***
orderAnseriformes         -0.134276 -0.218401 -0.046738 59900 0.003272 **
orderApodiformes           0.101579 -0.041054 0.236021 59600 0.144674
orderBucerotiformes        0.040000 -0.080951 0.157990 60680 0.507980
orderCariamiformes         0.080218 -0.066951 0.227669 59900 0.279399
orderCharadriiformes       0.009452 -0.083092 0.105417 59900 0.840534
orderCiconiiformes        -0.040935 -0.142712 0.060512 59900 0.416561
orderColumbiformes         0.065267 -0.066617 0.193642 59036 0.326811
orderCoraciiformes         0.082562 -0.078494 0.248061 61463 0.313222
orderGalliformes          -0.001747 -0.102287 0.099458 58770 0.978030
orderGruiformes           -0.089967 -0.189241 0.007686 59900 0.072287 .
orderMusophagiformes      -0.085619 -0.272281 0.096474 59900 0.365810
orderPasseriformes         0.086554 -0.003241 0.175723 59900 0.057763 .
orderPelecaniformes       -0.074397 -0.171168 0.018949 59900 0.121736
orderPhoenicopteriformes -0.291942 -0.439996 -0.144614 59900 0.000301
***
orderProcellariiformes    -0.039027 -0.134402 0.057526 60610 0.413523
orderPsittaciformes       -0.038279 -0.136047 0.057135 61190 0.431419
orderRheiformes           -0.206340 -0.349504 -0.061964 59900 0.006511 **
orderSphenisciformes      -0.136905 -0.243893 -0.031810 59900 0.012721 *
orderStrigiformes          0.034768 -0.119034 0.189827 59900 0.656528
orderSuliformes           -0.212732 -0.359269 -0.062097 61467 0.006344 **
---
Signif. codes:  0 ‘***’ 0.001 ‘**’ 0.01 ‘*’ 0.05 ‘.’ 0.1 ‘ ’ 1
```

#### Glycation individuals

```
 Iterations = 10001:5999901
 Thinning interval  = 100
 Sample size  = 59900
```

```
 DIC: 2391.883

 G-structure:  ~species

        post.mean l-95% CI u-95% CI eff.samp
species     8.851    5.417    12.71    59900

 R-structure:  ~units

      post.mean l-95% CI u-95% CI eff.samp
units     8.099    6.994    9.261    59900

 Location effects: Glycation ~ order

                         post.mean l-95% CI u-95% CI eff.samp pMCMC
(Intercept)                23.3396  19.5523 27.2678 58787  < 2e-05 ***
orderAnseriformes          -9.4202 -13.6002 -5.1701 59146  < 2e-05 ***
orderApodiformes            1.5038  -5.4759  8.6581 59049 0.669516
orderBucerotiformes        -4.0335 -10.1095  1.8861 59082 0.185543
orderCariamiformes         -1.8113  -9.1594  5.6689 59900 0.629149
orderCasuariiformes       -11.9476 -20.6864 -2.7884 59900 0.009182 **
orderCharadriiformes       -2.4454  -7.0388  1.9826 59129 0.285576
orderCiconiiformes         -3.0298  -8.0497  2.0927 59187 0.237930
orderColumbiformes         -2.4171  -7.4663  2.7121 59900 0.347112
orderCoraciiformes          0.5462  -7.5072  8.6354 56042 0.894124
orderGalliformes           -5.4805 -10.3152 -0.5489 59900 0.029649 *
orderGruiformes            -0.8043  -5.8130  3.9945 61375 0.745643
orderMusophagiformes       -1.3513  -6.6584  3.7670 59900 0.604875
orderPasseriformes         -5.2095  -9.6273 -0.7066 58879 0.023306 *
orderPelecaniformes        -5.7022 -10.3614 -1.0444 58508 0.017596 *
orderPhoenicopteriformes  -12.8877 -18.7001 -7.0605 59900 6.68e-05 ***
orderProcellariiformes     -1.9987  -6.7284  2.7765 59146 0.406912
orderPsittaciformes        -8.4219 -12.9810 -3.6229 58856 0.000835 ***
orderRheiformes             0.4280  -6.8066  7.5473 59900 0.908414
orderSphenisciformes       -6.6127 -11.8293 -1.2290 59900 0.015559 *
orderStrigiformes           1.9480  -4.2668  8.1967 59900 0.533623
orderSuliformes           -10.1649 -17.7348 -2.8516 56705 0.007546 **
---
Signif. codes:  0 ‘***’ 0.001 ‘**’ 0.01 ‘*’ 0.05 ‘.’ 0.1 ‘ ’ 1
```

### VIF models

#### Glucose averages

```
                   GVIF Df GVIF^(1/(2*Df))
Diet           1.713187  4        1.069610
Centered_logBM 1.494981  1        1.222694
Procedence     1.728314  1        1.314654
```

#### Glucose averages life history

```
                           GVIF Df GVIF^(1/(2*Df))
Diet                   3.923876  4        1.186354
Centered_logBM         1.991121  1        1.411071
poly(ML, 2, raw = TRUE) 1.822946  2        1.161966
CM                     2.657595  1        1.630213
DT                     1.392595  1        1.180083
Procedence             1.924601  1        1.387300
```

#### Glycation averages

```
                        GVIF Df GVIF^(1/(2*Df))
Diet                1.295990  4        1.032940
Centered_logBM      1.641464  1        1.281196
Centered_logGlucose 1.620827  1        1.273117
```

#### Glycation averages life-history

| | GVIF | Df | GVIF^(1/(2*Df)) |
|---|---|---|---|
| Diet | 3.151870 | 4 | 1.154306 |
| Centered_logBM | 1.971214 | 1 | 1.403999 |
| Centered_logGlucose | 2.188302 | 1 | 1.479291 |
| ML | 1.380756 | 1 | 1.175056 |
| CM | 2.098269 | 1 | 1.448540 |
| DT | 1.295233 | 1 | 1.138083 |

#### Glycation averages life-history without glucose

| | GVIF | Df | GVIF^(1/(2*Df)) |
|---|---|---|---|
| Diet | 2.617272 | 4 | 1.127797 |
| Centered_logBM | 1.354690 | 1 | 1.163912 |
| ML | 1.314165 | 1 | 1.146370 |
| CM | 2.095152 | 1 | 1.447464 |
| DT | 1.276962 | 1 | 1.130027 |

#### Glucose individuals

| | GVIF | Df | GVIF^(1/(2*Df)) |
|---|---|---|---|
| Diet | 3.216096 | 4 | 1.157221 |
| Centered_logBM | 2.237036 | 1 | 1.495672 |
| Procedence | 2.252484 | 1 | 1.500828 |

#### Glucose individuals life-history

| | GVIF | Df | GVIF^(1/(2*Df)) |
|---|---|---|---|
| Diet | 21.782694 | 4 | 1.469819 |
| Centered_logBM | 5.093717 | 1 | 2.256927 |
| poly(ML, 2, raw = TRUE) | 1.966756 | 2 | 1.184234 |
| CM | 3.876582 | 1 | 1.968904 |
| DT | 1.621283 | 1 | 1.273296 |
| Procedence | 3.362333 | 1 | 1.833666 |

#### Glycation individuals

| | GVIF | Df | GVIF^(1/(2*Df)) |
|---|---|---|---|
| Diet | 1.643911 | 4 | 1.064106 |
| Centered_logBM | 2.189558 | 1 | 1.479715 |
| Centered_logGlucose | 1.946567 | 1 | 1.395194 |

#### Glycation individuals life-history

| | GVIF | Df | GVIF^(1/(2*Df)) |
|---|---|---|---|
| Diet | 6.218989 | 4 | 1.256652 |
| Centered_logBM | 3.413201 | 1 | 1.847485 |
| Centered_logGlucose | 2.074540 | 1 | 1.440326 |
| ML | 1.296298 | 1 | 1.138551 |
| CM | 3.114929 | 1 | 1.764916 |
| DT | 1.446495 | 1 | 1.202703 |

#### Glycation individuals life-history

| | GVIF | Df | GVIF^(1/(2*Df)) |
|---|---|---|---|
| Diet | 5.454035 | 4 | 1.236203 |
| Centered_logBM | 2.682914 | 1 | 1.637960 |
| ML | 1.291340 | 1 | 1.136371 |
| CM | 3.111995 | 1 | 1.764085 |
| DT | 1.433119 | 1 | 1.197129 |

#### Glucose age & sex

| | GVIF | Df | GVIF^(1/(2*Df)) |
|---|---|---|---|
| Centered_logBM | 1.148783 | 1 | 1.071813 |
| poly(logit(Age_relative, FALSE), 2, raw = TRUE) | 1.146260 | 2 | 1.034715 |
| Sex | 1.008919 | 1 | 1.004450 |

#### Glycation age & sex

```
                                              GVIF Df GVIF^(1/(2*Df))
Centered_logBM                            1.949521  1        1.396252
Centered_logGlucose                       2.042604  1        1.429197
poly(logit(Age_relative, FALSE), 2, raw = TRUE) 1.200827  2        1.046815
Sex                                       1.029088  1        1.014440
```

### Post-hoc comparisons for dietary categories

A=Omnivores; B=Terrestrial carnivores; C=Aquatic predators; D=Herbivores; E=Frugivores/Granivores

#### Glucose averages

```
> HPDinterval(AG_Dif_BC)
         lower    upper
var1 -1.414069 1.466687
attr(,"Probability")
[1] 0.95
> HPDinterval(AG_Dif_BD)
         lower    upper
var1 -1.222618 1.308603
attr(,"Probability")
[1] 0.95
> HPDinterval(AG_Dif_BE)
         lower    upper
var1 -1.393724 1.500328
attr(,"Probability")
[1] 0.95
> HPDinterval(AG_Dif_CD)
         lower    upper
var1 -1.551009 1.566369
attr(,"Probability")
[1] 0.95
> HPDinterval(AG_Dif_CE)
         lower    upper
var1 -1.717679 1.821919
attr(,"Probability")
[1] 0.95
> HPDinterval(AG_Dif_DE)
         lower    upper
var1 -1.446564 1.520309
attr(,"Probability")
[1] 0.95
```

#### Glucose averages life history

```
> HPDinterval(AGLH_Dif_BC)
        lower    upper
var1 -1.54855 1.531426
attr(,"Probability")
[1] 0.95
> HPDinterval(AGLH_Dif_BD)
         lower    upper
var1 -2.140114 2.182302
attr(,"Probability")
[1] 0.95
> HPDinterval(AGLH_Dif_BE)
         lower    upper
var1 -7.737039 7.508915
attr(,"Probability")
[1] 0.95
> HPDinterval(AGLH_Dif_CD)
         lower    upper
var1 -2.146583 2.220799
attr(,"Probability")
[1] 0.95
```

```
> HPDinterval(AGLH_Dif_CE)
         lower    upper
var1 -7.488458 7.862649
attr(,"Probability")
[1] 0.95
> HPDinterval(AGLH_Dif_DE)
         lower    upper
var1 -7.751623 7.691805
attr(,"Probability")
[1] 0.95
```

#### Glycation averages

```
> HPDinterval(AGly_Dif_BC)
       lower    upper
var1 -1.2729 1.341971
attr(,"Probability")
[1] 0.95
> HPDinterval(AGly_Dif_BD)
         lower   upper
var1 -1.305844 1.52542
attr(,"Probability")
[1] 0.95
> HPDinterval(AGly_Dif_BE)
         lower    upper
var1 -1.056829 1.224983
attr(,"Probability")
[1] 0.95
> HPDinterval(AGly_Dif_CD)
         lower    upper
var1 -1.433015 1.563042
attr(,"Probability")
[1] 0.95
> HPDinterval(AGly_Dif_CE)
         lower    upper
var1 -1.282198 1.396203
attr(,"Probability")
[1] 0.95
> HPDinterval(AGly_Dif_DE)
        lower    upper
var1 -1.40408 1.318576
attr(,"Probability")
[1] 0.95
```

#### Glycation averages life-history

```
> HPDinterval(AGlyLH_Dif_BC)
        lower    upper
var1 -1.36308 1.540043
attr(,"Probability")
[1] 0.95
> HPDinterval(AGlyLH_Dif_BD)
         lower    upper
var1 -2.008801 2.234876
attr(,"Probability")
[1] 0.95
> HPDinterval(AGlyLH_Dif_BE)
         lower    upper
var1 -1.774074 2.117975
attr(,"Probability")
[1] 0.95
> HPDinterval(AGlyLH_Dif_CD)
         lower    upper
var1 -2.048012 2.107497
attr(,"Probability")
[1] 0.95
> HPDinterval(AGlyLH_Dif_CE)
```

```
         lower    upper
var1 -2.005551 2.19016
attr(,"Probability")
[1] 0.95
> HPDinterval(AGlyLH_Dif_DE)
         lower     upper
var1 -2.183145 2.284387
attr(,"Probability")
[1] 0.95
```

#### Glycation averages life-history without glucose

```
> HPDinterval(AGlyLH_NG_Dif_BC)
         lower     upper
var1 -1.369687 1.504995
attr(,"Probability")
[1] 0.95
> HPDinterval(AGlyLH_NG_Dif_BD)
         lower     upper
var1 -1.810076 2.055283
attr(,"Probability")
[1] 0.95
> HPDinterval(AGlyLH_NG_Dif_BE)
         lower     upper
var1 -1.678796 2.054028
attr(,"Probability")
[1] 0.95
> HPDinterval(AGlyLH_NG_Dif_CD)
         lower     upper
var1 -1.917022 2.053911
attr(,"Probability")
[1] 0.95
> HPDinterval(AGlyLH_NG_Dif_CE)
         lower     upper
var1 -1.923162 2.114277
attr(,"Probability")
[1] 0.95
> HPDinterval(AGlyLH_NG_Dif_DE)
         lower     upper
var1 -2.117618 2.130004
attr(,"Probability")
[1] 0.95
```

#### Glucose individuals

```
> HPDinterval(IG_Dif_BC)
           lower      upper
var1 -0.07163346 0.05929215
attr(,"Probability")
[1] 0.95
> HPDinterval(IG_Dif_BD)
           lower     upper
var1 -0.08404549 0.1570657
attr(,"Probability")
[1] 0.95
> HPDinterval(IG_Dif_BE)
            lower     upper
var1 -0.006626182 0.1846572
attr(,"Probability")
[1] 0.95
> HPDinterval(IG_Dif_CD)
           lower     upper
var1 -0.08960038 0.1648752
attr(,"Probability")
[1] 0.95
> HPDinterval(IG_Dif_CE)
           lower     upper
var1 -0.01340042 0.2039278
```

```
attr(,"Probability")
[1] 0.95
> HPDinterval(IG_Dif_DE)
           lower    upper
var1 -0.06618755 0.176413
attr(,"Probability")
[1] 0.95
```

#### Glucose individuals life-history

```
> HPDinterval(IGLH_Dif_BC)
           lower      upper
var1 -0.08234745 0.05308559
attr(,"Probability")
[1] 0.95
> HPDinterval(IGLH_Dif_BD)
           lower     upper
var1 -0.06588702 0.2218879
attr(,"Probability")
[1] 0.95
> HPDinterval(IGLH_Dif_BE)
          lower     upper
var1 -0.1173287 0.2604363
attr(,"Probability")
[1] 0.95
> HPDinterval(IGLH_Dif_CD)
           lower     upper
var1 -0.05760069 0.2320285
attr(,"Probability")
[1] 0.95
> HPDinterval(IGLH_Dif_CE)
          lower    upper
var1 -0.1169439 0.282375
attr(,"Probability")
[1] 0.95
> HPDinterval(IGLH_Dif_DE)
          lower     upper
var1 -0.2299858 0.2335929
attr(,"Probability")
[1] 0.95
```

#### Glycation individuals

```
> HPDinterval(IGly_Dif_BC)
           lower     upper
var1 -0.01230547 0.1285872
attr(,"Probability")
[1] 0.95
> HPDinterval(IGly_Dif_BD)
           lower     upper
var1 -0.05044503 0.2006516
attr(,"Probability")
[1] 0.95
> HPDinterval(IGly_Dif_BE)
           lower     upper
var1 -0.01012096 0.1853377
attr(,"Probability")
[1] 0.95
> HPDinterval(IGly_Dif_CD)
          lower     upper
var1 -0.1140737 0.1468223
attr(,"Probability")
[1] 0.95
> HPDinterval(IGly_Dif_CE)
           lower     upper
var1 -0.07078775 0.1368791
attr(,"Probability")
[1] 0.95
```

```
> HPDinterval(IGly_Dif_DE)
          lower     upper
var1 -0.1086503 0.1426304
attr(,"Probability")
[1] 0.95
```

#### Glycation individuals life-history

```
> HPDinterval(IGlyLH_Dif_BC)
            lower     upper
var1 -0.001756023 0.1502896
attr(,"Probability")
[1] 0.95
> HPDinterval(IGlyLH_Dif_BD)
           lower     upper
var1 -0.03802143 0.2812876
attr(,"Probability")
[1] 0.95
> HPDinterval(IGlyLH_Dif_BE)
          lower     upper
var1 -0.1907319 0.2172327
attr(,"Probability")
[1] 0.95
> HPDinterval(IGlyLH_Dif_CD)
         lower     upper
var1 -0.117811 0.2090437
attr(,"Probability")
[1] 0.95
> HPDinterval(IGlyLH_Dif_CE)
          lower     upper
var1 -0.2768476 0.1317622
attr(,"Probability")
[1] 0.95
> HPDinterval(IGlyLH_Dif_DE)
         lower     upper
var1 -0.359544 0.1331044
attr(,"Probability")
[1] 0.95
```

#### Glycation individuals life-history without glucose

```
> HPDinterval(IGlyLH_NG_Dif_BC)
             lower     upper
var1 -0.0008724393 0.1537172
attr(,"Probability")
[1] 0.95
> HPDinterval(IGlyLH_NG_Dif_BD)
           lower     upper
var1 -0.03474923 0.2953013
attr(,"Probability")
[1] 0.95
> HPDinterval(IGlyLH_NG_Dif_BE)
          lower     upper
var1 -0.1988301 0.2156449
attr(,"Probability")
[1] 0.95
> HPDinterval(IGlyLH_NG_Dif_CD)
          lower     upper
var1 -0.1067734 0.2297418
attr(,"Probability")
[1] 0.95
> HPDinterval(IGlyLH_NG_Dif_CE)
          lower     upper
var1 -0.2779184 0.1386907
attr(,"Probability")
[1] 0.95
> HPDinterval(IGlyLH_NG_Dif_DE)
          lower     upper
```

```
var1 -0.3831259 0.1231232
attr(,"Probability")
[1] 0.95
```

### Within species repeatability

#### Glucose

```
Linear mixed model fit by REML ['lmerMod']
Formula: logGlucose ~ (1 | species)
   Data: Bird.caracIndGlu

REML criterion at convergence: -875.1

Scaled residuals: 
    Min      1Q  Median      3Q     Max 
-3.7868 -0.5688 -0.0078  0.5683  2.8026 

Random effects:
 Groups   Name        Variance Std.Dev.
 species  (Intercept) 0.009768 0.09883 
 Residual             0.003869 0.06220 
Number of obs: 389, groups:  species, 75

Fixed effects:
            Estimate Std. Error t value
(Intercept)  2.39913    0.01214   197.7

Repeatability estimation using the lmm method

Repeatability for species
R  = 0.716
SE = 0.042
CI = [0.619, 0.785]
P  = 8.57e-80 [LRT]
     NA [Permutation]
```

#### Glycation

```
Linear mixed model fit by REML ['lmerMod']
Formula: Glycation ~ (1 | species)
   Data: Bird.caracIndGly

REML criterion at convergence: 2521.1

Scaled residuals: 
    Min      1Q  Median      3Q     Max 
-4.3646 -0.4484 -0.0123  0.4749  3.4483 

Random effects:
 Groups   Name        Variance Std.Dev.
 species  (Intercept) 18.898   4.347   
 Residual              7.994   2.827   
Number of obs: 471, groups:  species, 88

Fixed effects:
            Estimate Std. Error t value
(Intercept)  18.4571     0.4921   37.51


Repeatability estimation using the lmm method

Repeatability for species
R  = 0.703
SE = 0.042
CI = [0.603, 0.767]
P  = 3.53e-86 [LRT]
     NA [Permutation]
```

### Variances explained by the tree

#### Glucose averages

```
0.3060753
             lower     upper
var1 9.658695e-07 0.8365601
attr(,"Probability")
[1] 0.95
```

#### Glucose averages life history

```
0.31353
             lower     upper
var1 1.769981e-06 0.8672609
attr(,"Probability")
[1] 0.95
```

#### Glycation averages

```
0.3324061
            lower     upper
var1 1.64774e-06 0.8480055
attr(,"Probability")
[1] 0.95
```

#### Glycation averages life-history

```
0.333346
             lower     upper
var1 2.934731e-06 0.8685776
attr(,"Probability")
[1] 0.95
```

#### Glycation averages life-history without glucose

```
0.5373031
          lower     upper
var1 0.07157126 0.9932748
attr(,"Probability")
[1] 0.95
```

#### Glucose individuals

```
0.5161435
         lower     upper
var1 0.2761292 0.7415861
attr(,"Probability")
[1] 0.95
```

#### Glucose individuals life-history

```
0.5025716
         lower     upper
var1 0.2420428 0.7546559
attr(,"Probability")
[1] 0.95
```

#### Glycation individuals

```
0.4590051
         lower    upper
```

```
var1 0.2151646 0.696498
attr(,"Probability")
[1] 0.95
```

#### Glycation individuals life-history

```
0.46431
         lower     upper
var1 0.1987901 0.7324513
attr(,"Probability")
[1] 0.95
```

#### Glycation individuals life-history without glucose

```
0.485408
         lower     upper
var1 0.2080762 0.7528297
attr(,"Probability")
[1] 0.95
```

### Traces and posterior distributions

#### Glucose averages

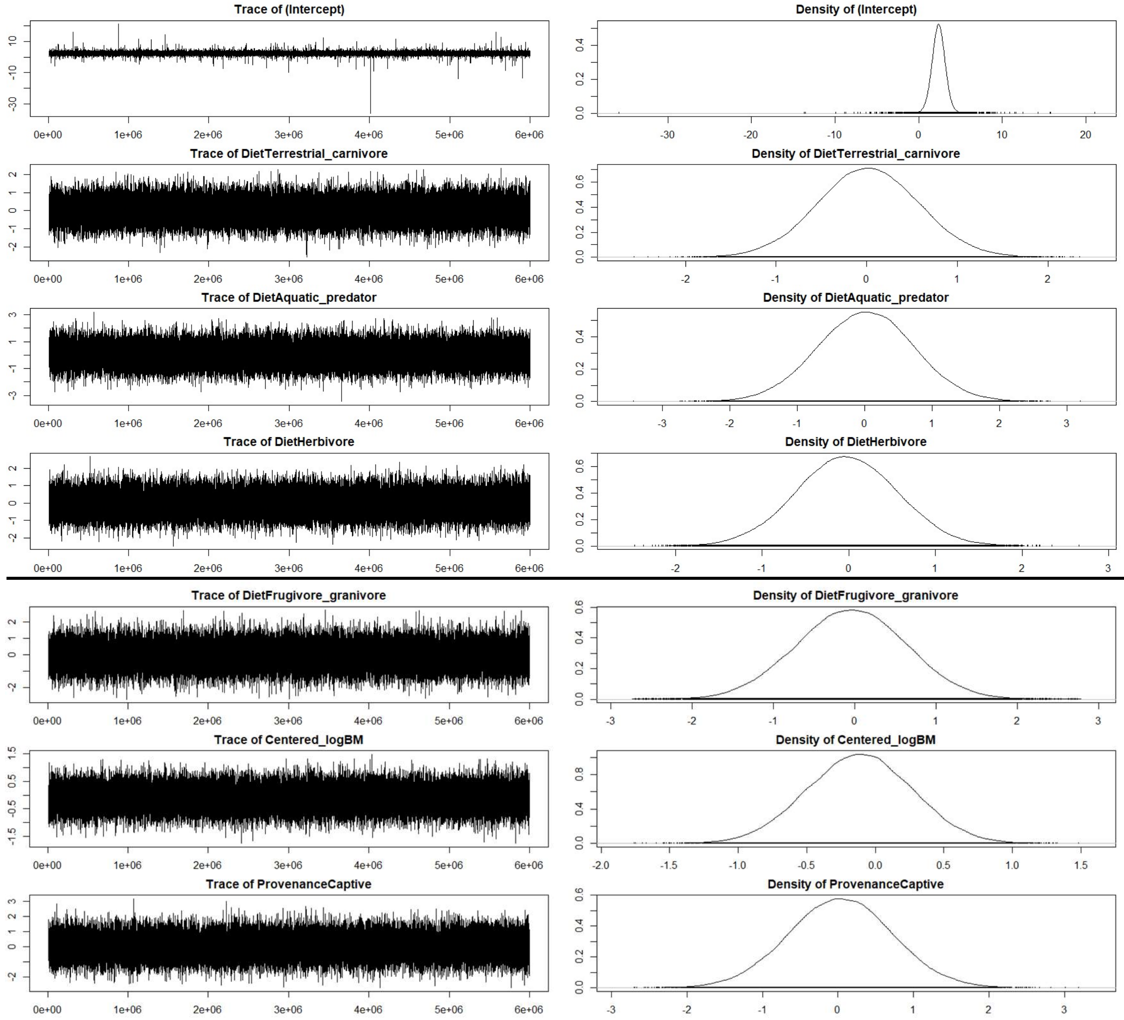

### Glucose averages life history

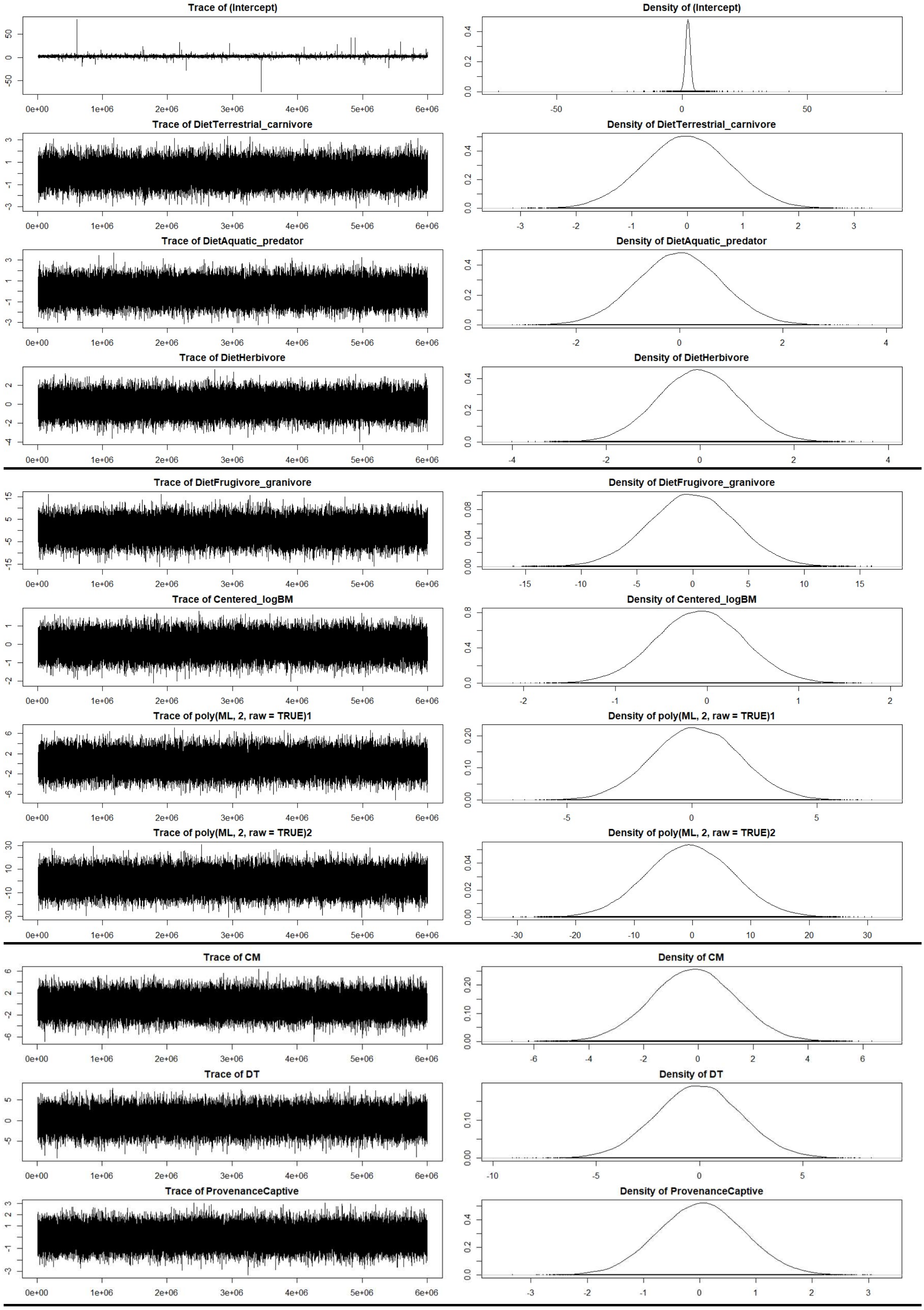

### Glycation averages

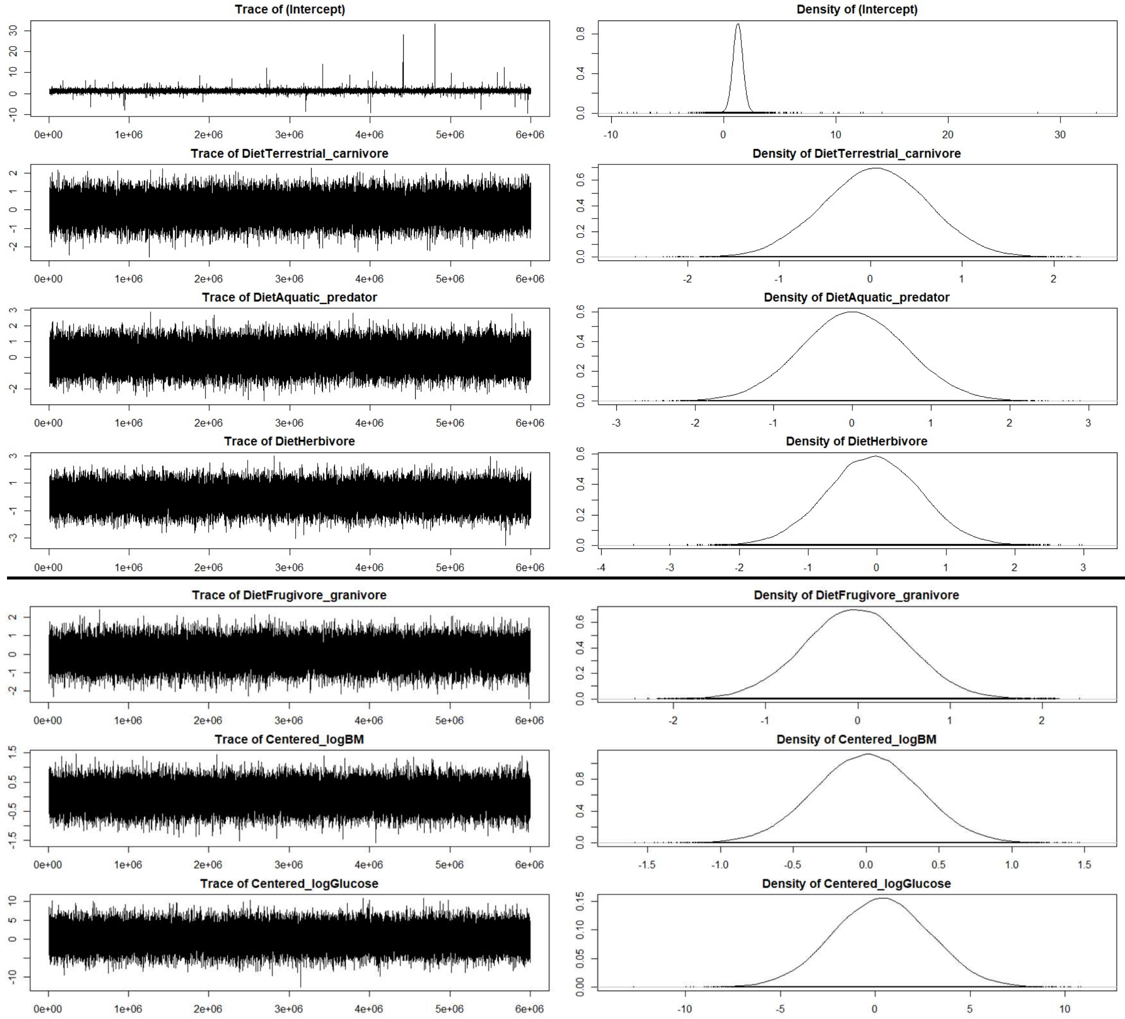


### Glycation averages life-history

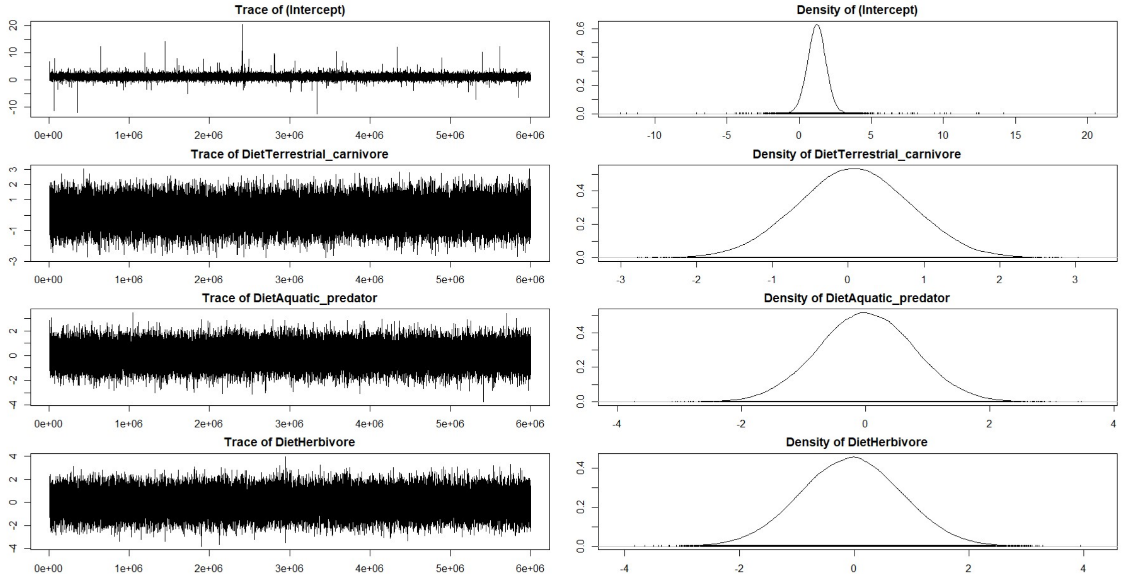

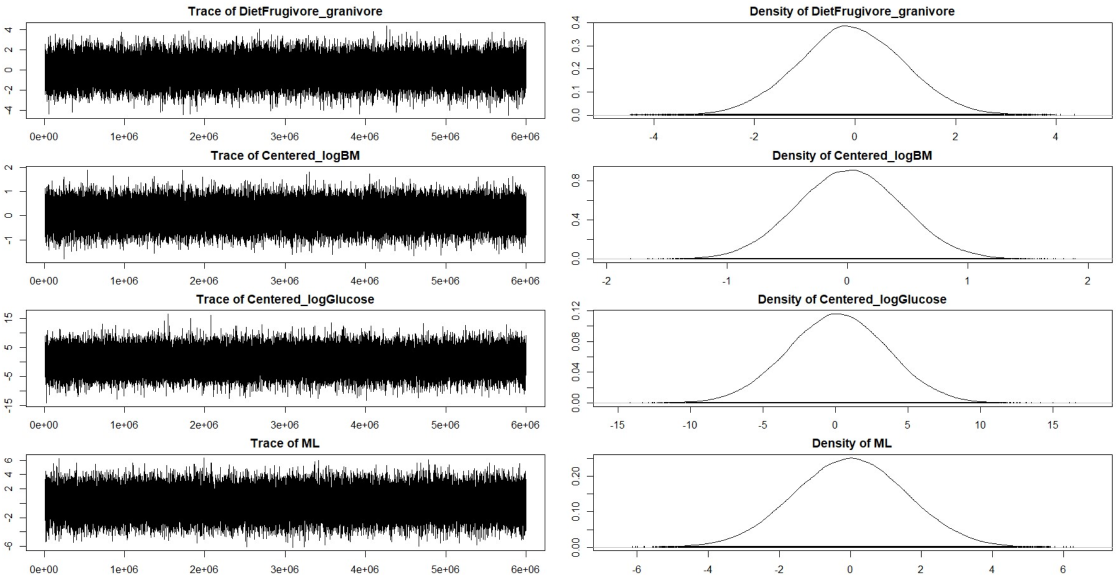


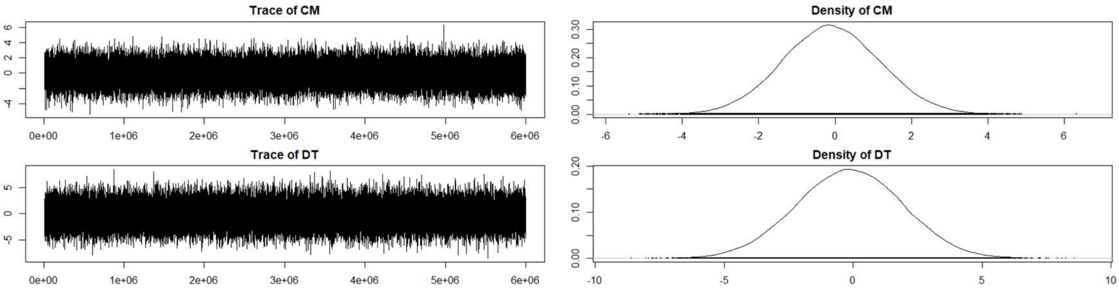


### Glycation averages life-history without glucose

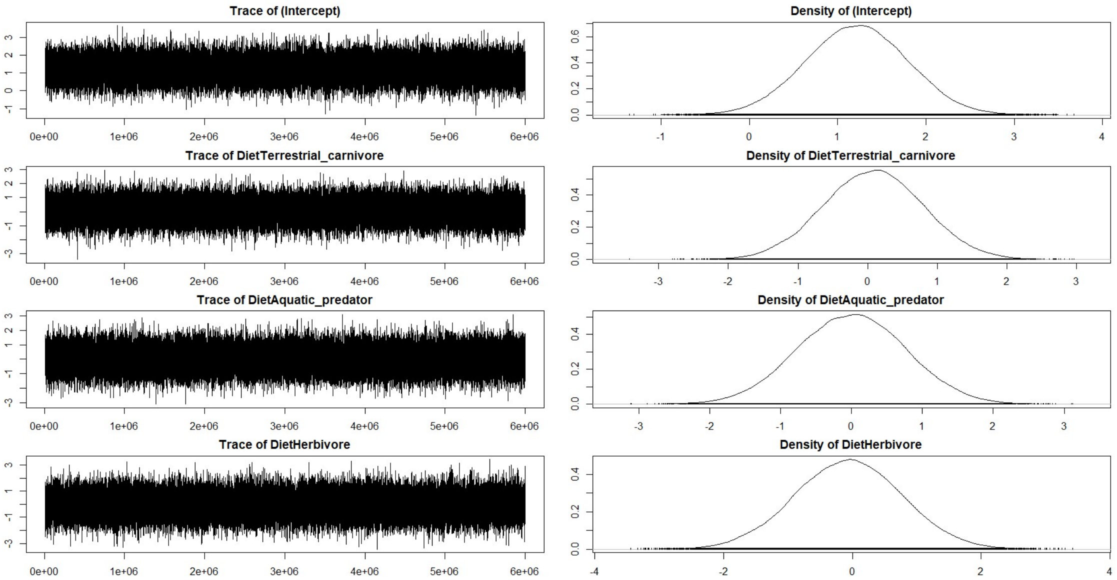

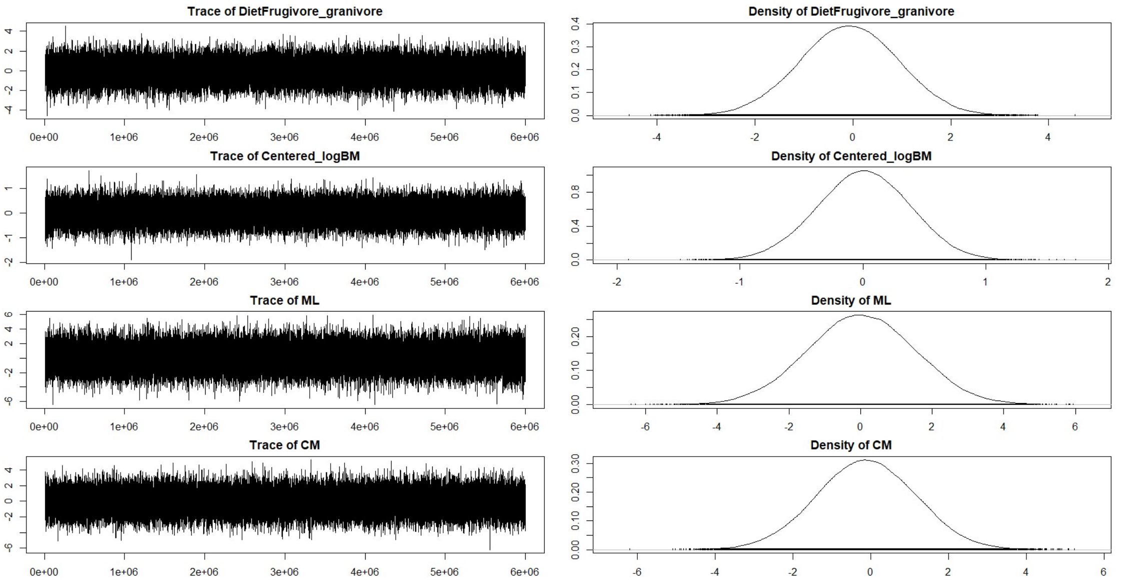


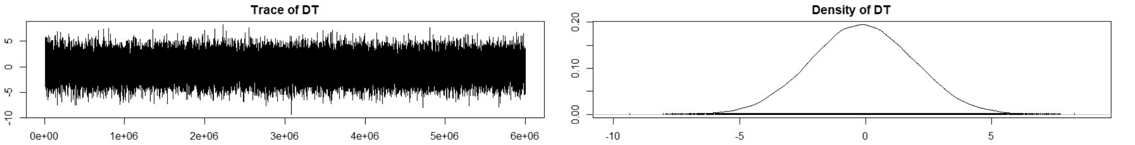


#### Glucose individuals

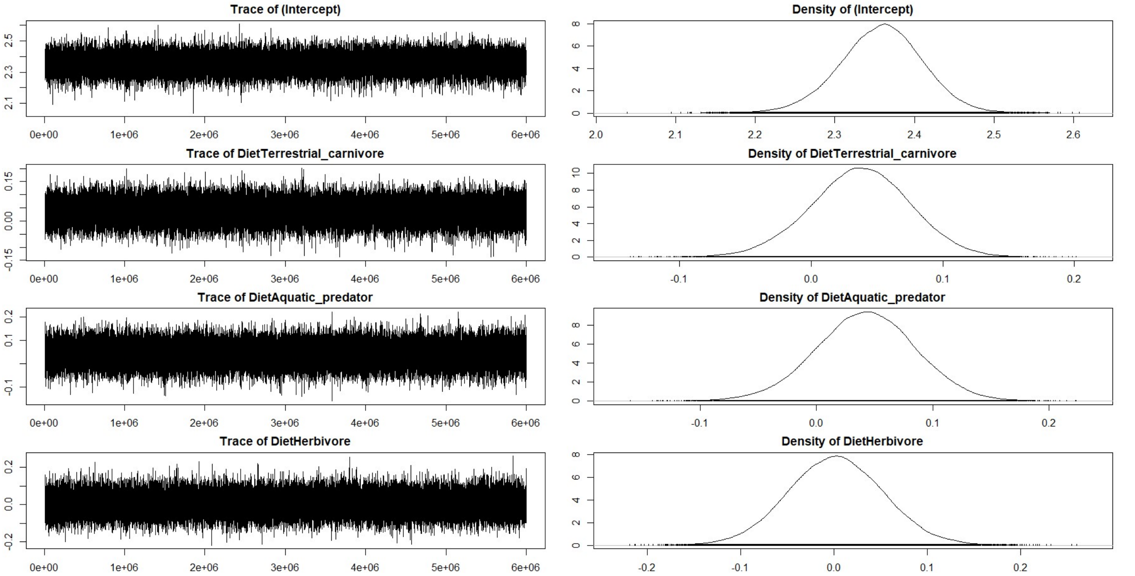

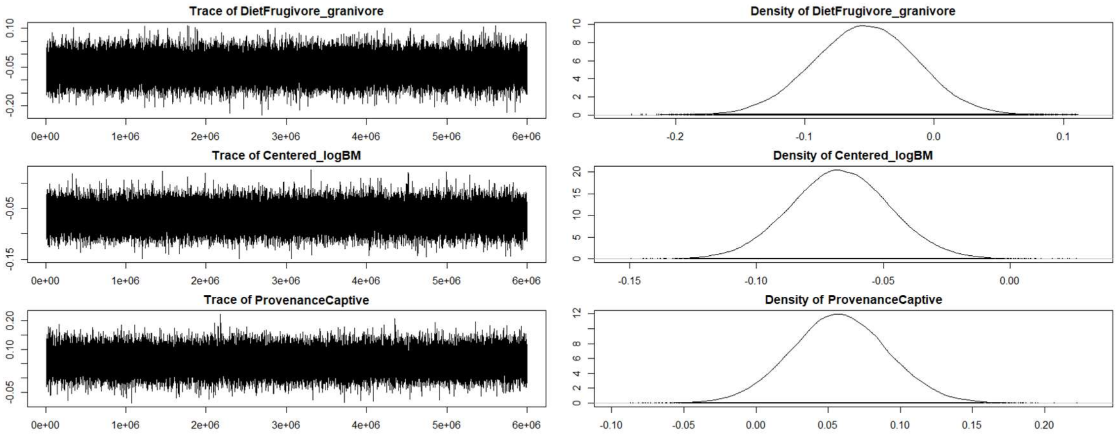


#### Glucose individuals life-history

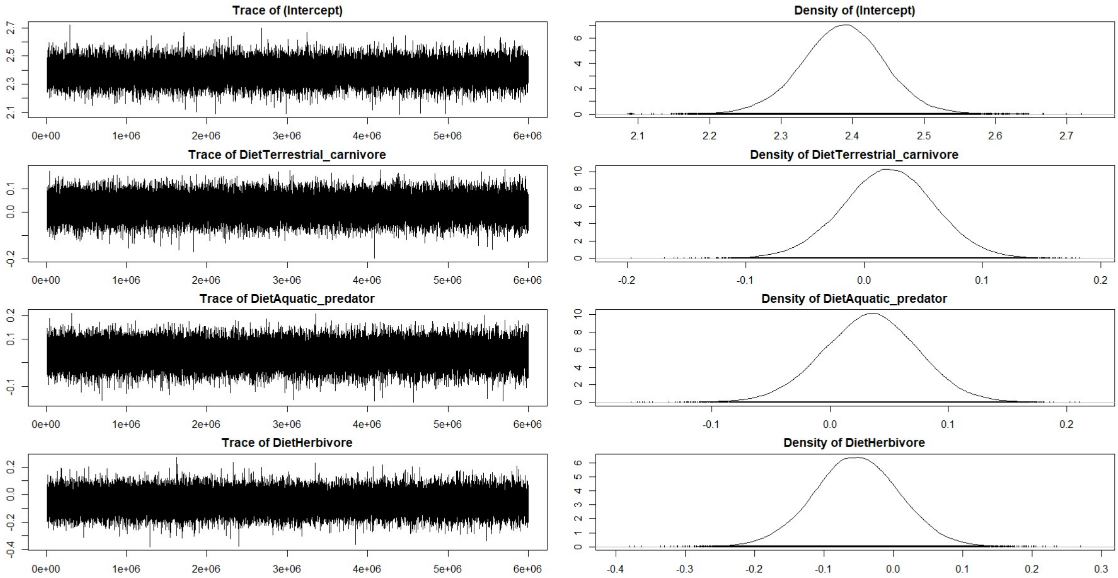


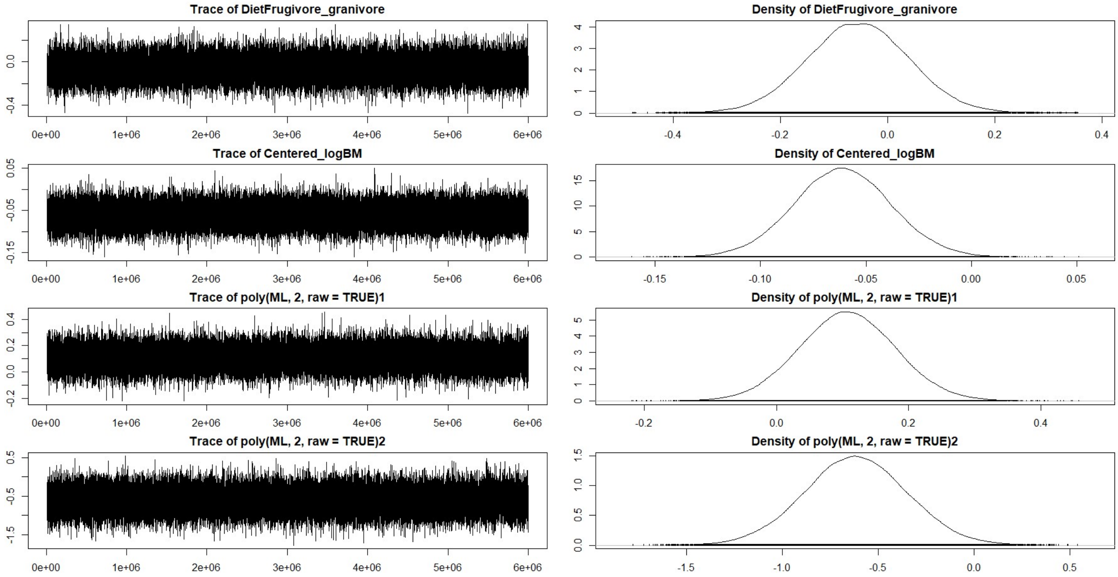

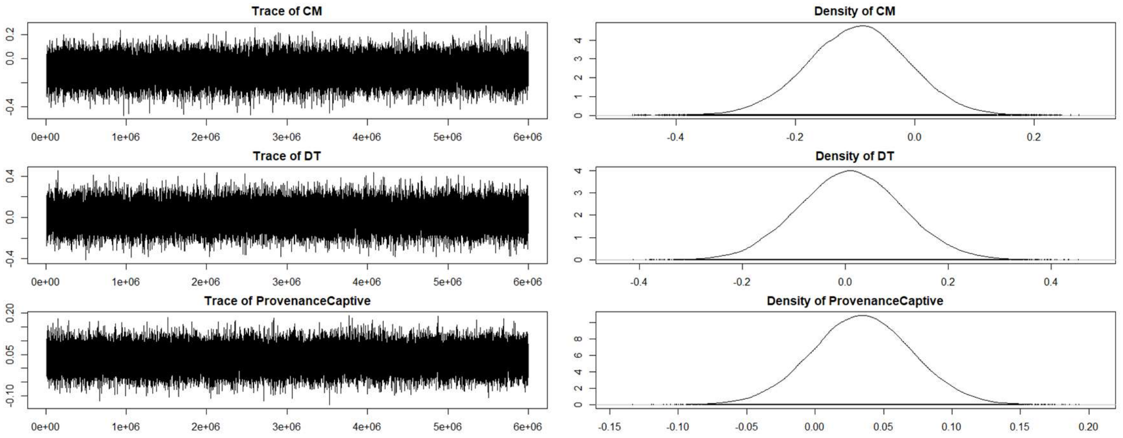


#### **Glycation individuals**

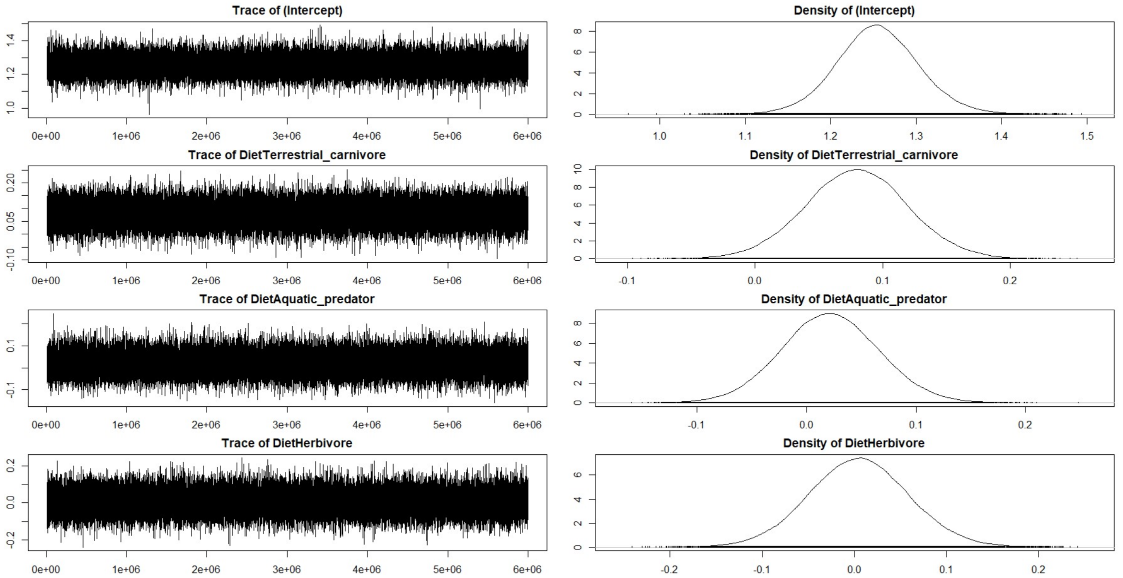


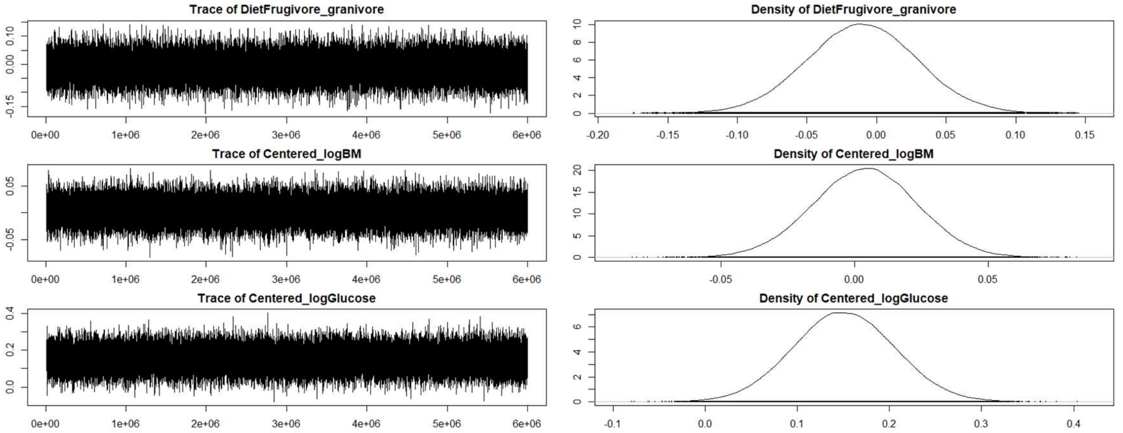

### Glycation individuals life-history

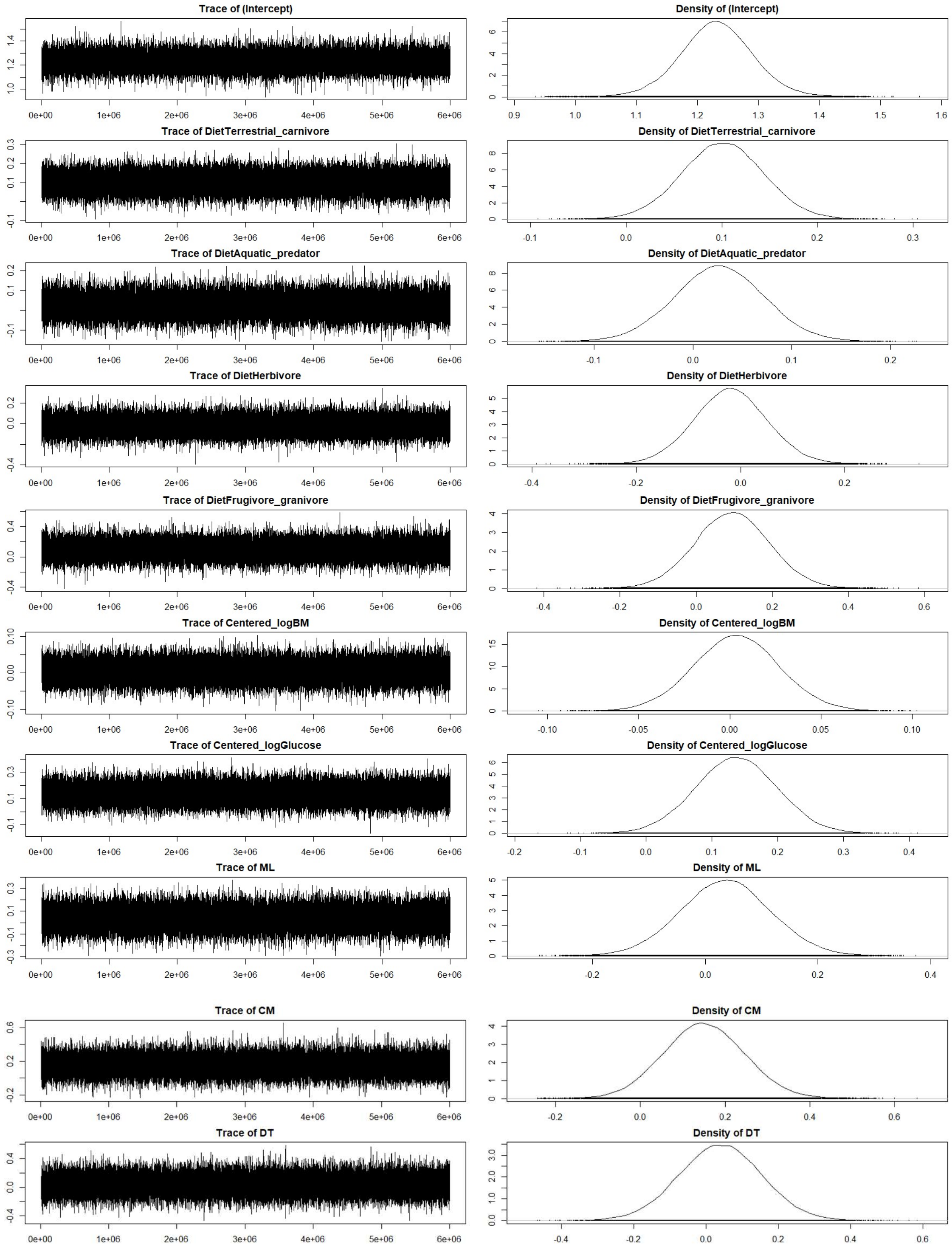

#### Glycation individuals life-history without glucose

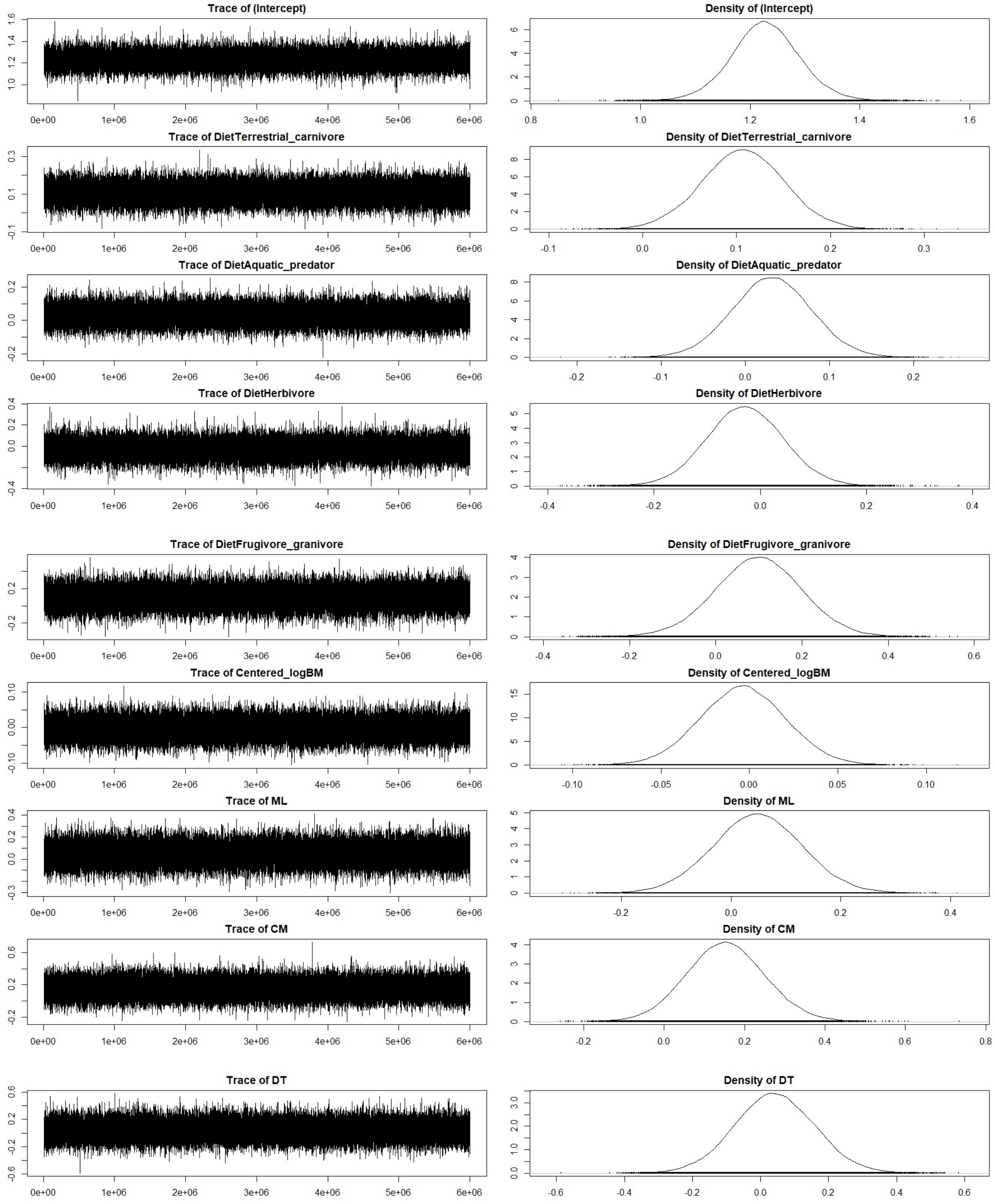


**Figure ESM1.1** Trace showing convergence of the models and posterior density plots of the parameters estimated.

### Supplementary analyses of stress on glucose

#### Glucose repeatability

```
Linear mixed model fit by REML. t-tests use Satterthwaite's method ['l
merModLmerTest']
Formula: Glucose ~ (1 | SpeciesBirdTree/Individual)
   Data: Tomasek2022

REML criterion at convergence: 44335.9
```

```
Scaled residuals:
    Min      1Q  Median      3Q     Max 
-4.6683 -0.5772  0.0246  0.5585  4.0989 

Random effects:
 Groups                      Name        Variance Std.Dev.
 Individual:SpeciesBirdTree (Intercept) 5.188    2.278   
 SpeciesBirdTree            (Intercept) 2.222    1.490   
 Residual                               6.204    2.491   
Number of obs: 8862, groups:  Individual:SpeciesBirdTree, 1705; Specie
sBirdTree, 158

Fixed effects:
            Estimate Std. Error       df t value Pr(>|t|)    
(Intercept)  13.8677     0.1496 146.9083   92.67   <2e-16 ***


Repeatability estimation using the lmm method
```

<u>Repeatability for SpeciesBirdTree</u>

```
R  = 0.163
SE = 0.024
CI = [0.117, 0.209]
P  = 0 [LRT]
     NA [Permutation]
```

**<u>For individuals within species:</u>**

```
> 5.188/(5.188+2.222+6.204)
[1] 0.3810783
```

#### Glucose repeatability (with stress effects)

```
Linear mixed model fit by REML. t-tests use Satterthwaite's method ['l
merModLmerTest']
Formula: Glucose ~ Time + (1 | SpeciesBirdTree/Individual)
   Data: Tomasek2022

REML criterion at convergence: 40911.9

Scaled residuals:
    Min      1Q  Median      3Q     Max 
-5.0618 -0.5405 -0.0221  0.5398  4.6158 

Random effects:
 Groups                      Name        Variance Std.Dev.
 Individual:SpeciesBirdTree (Intercept) 5.499    2.345   
 SpeciesBirdTree            (Intercept) 2.267    1.506   
 Residual                               3.864    1.966   
Number of obs: 8862, groups:  Individual:SpeciesBirdTree, 1705; Specie
sBirdTree, 158

Fixed effects:
             Estimate Std. Error        df t value Pr(>|t|)    
(Intercept) 1.211e+01  1.525e-01 1.573e+02   79.41   <2e-16 ***
TimeG15_    2.936e+00  5.458e-02 7.341e+03   53.79   <2e-16 ***
TimeG30_    2.905e+00  4.976e-02 7.271e+03   58.37   <2e-16 ***
---
Signif. codes:  0 '***' 0.001 '**' 0.01 '*' 0.05 '.' 0.1 ' ' 1

Correlation of Fixed Effects:
         (Intr) TmG15_
TimeG15_ -0.140       
```

```
TimeG30_ -0.153  0.442
```

```
Levene's Test for Homogeneity of Variance (center = median)
        Df F value    Pr(>F)    
group    2  455.14 < 2.2e-16 ***
      8859                      
```

```
Kruskal-Wallis rank sum test

data:  Glucose by Time
Kruskal-Wallis chi-squared = 1347.7, df = 2, p-value < 2.2e-16
```

```
Repeatability estimation using the lmm method
```

**Repeatability for SpeciesBirdTree**

```
R  = 0.195
SE = 0.027
CI = [0.141, 0.248]
P  = 0 [LRT]
     NA [Permutation]
```

**For individuals within species:**

```
> 5.499/(5.499+2.267+3.864)
[1] 0.4728289
```

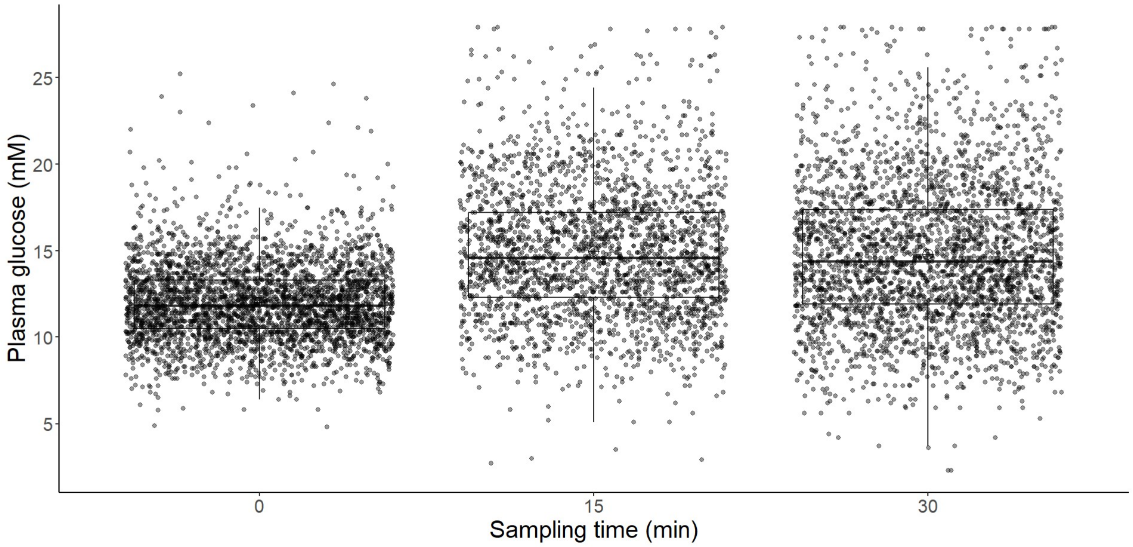


**Figure ESM1.2** Glucose variation (in mM) with sampling time at 0, 15 and 30 min, showing clear heteroskedasticity. Data publicly available from Tomasek et al. 2022 (see ESM6).
