## Supplementary material for "Variation in albumin glycation rates in birds suggests resistance to relative hyperglycaemia rather than conformity to the pace of life syndrome hypothesis": ESM1-4 & 6: ESM2.pdf

### Electronic Supplementary Material 2 (ESM2)

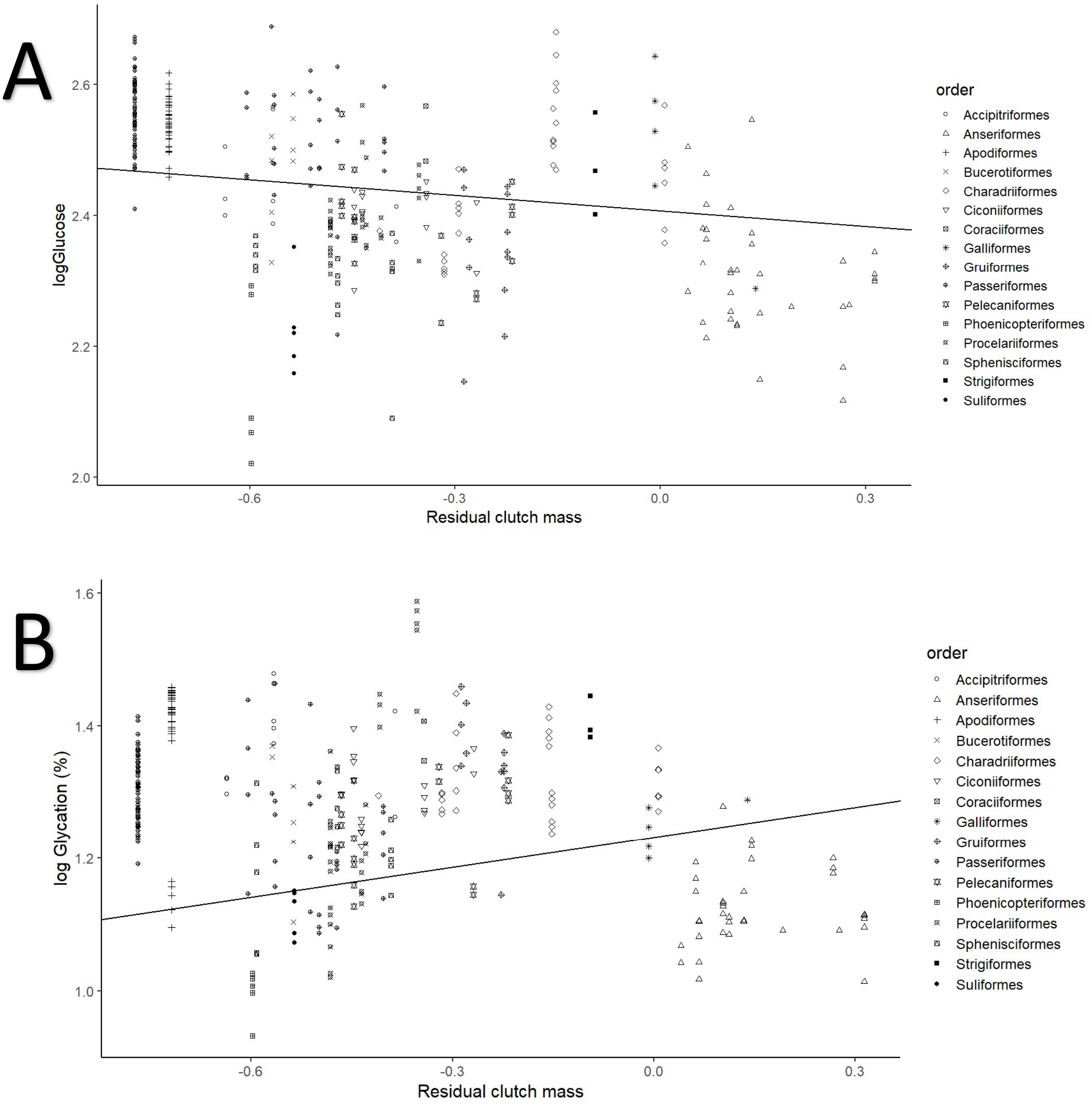


**Figure ESM2.1.** Variation in **A** $log_{10}$(plasma glucose) in mg/dl and **B** $log_{10}$(albumin glycation) levels (%) with respect to residuals of body mass-adjusted clutch mass of the species. This is calculated by multiplying egg mass times clutch size, and then performing a pGLS (see ESM6) of the decimal logarithm of this value against the decimal logarithm of body mass, to finally extract the residuals of such regression. Glycation levels are reported as a percentage of glycated albumin versus total albumin. The legends at right show the different orders to which our species belong.

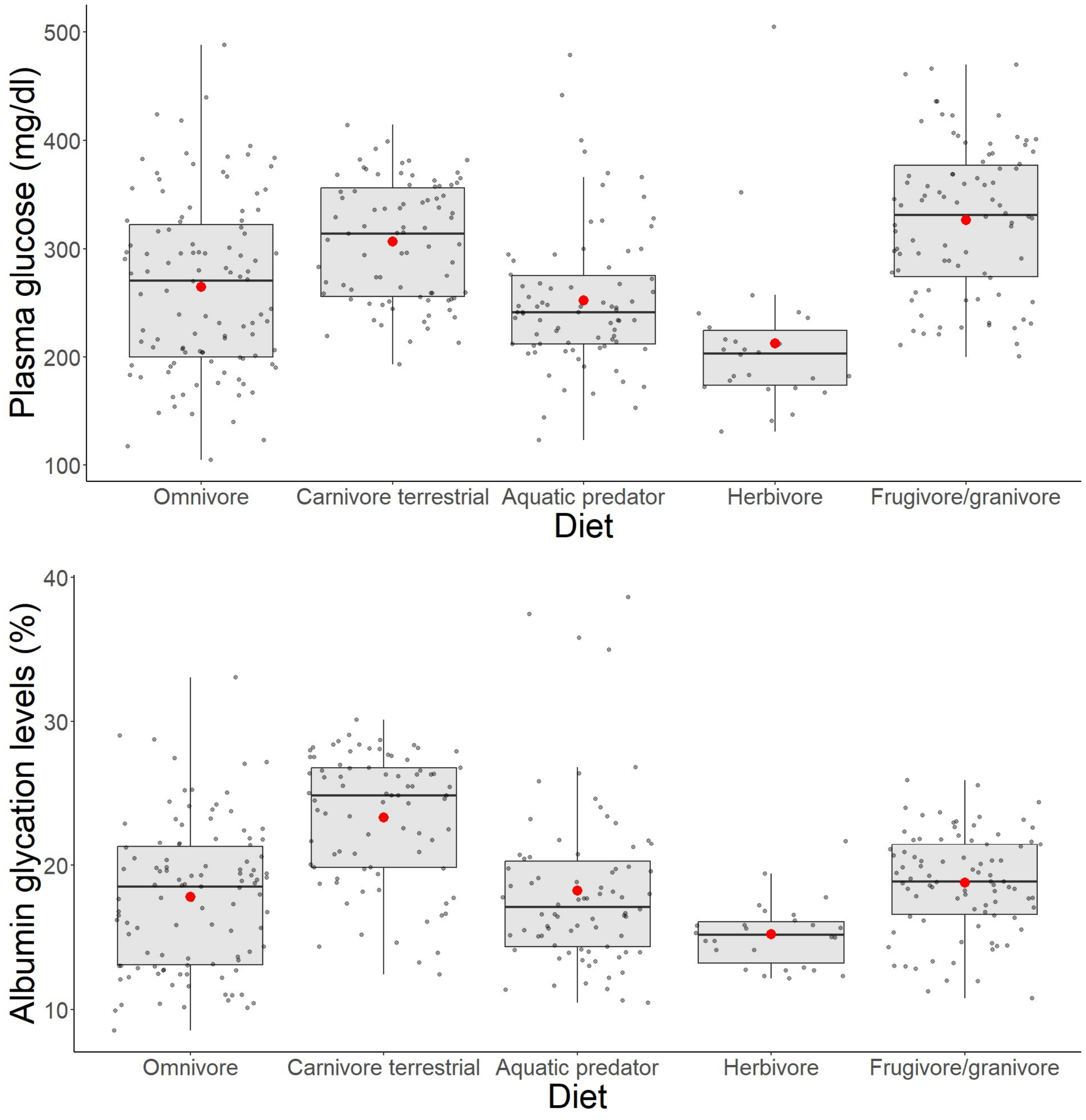


**Figure ESM2.2.** Representation of raw individual data on (**A**) plasma glucose levels and (**B**) albumin glycation in function of birds' diet. Glucose levels are given in mg/dl, while glycation levels are a percentage of total plasma albumin which is found to be glycated.

A1

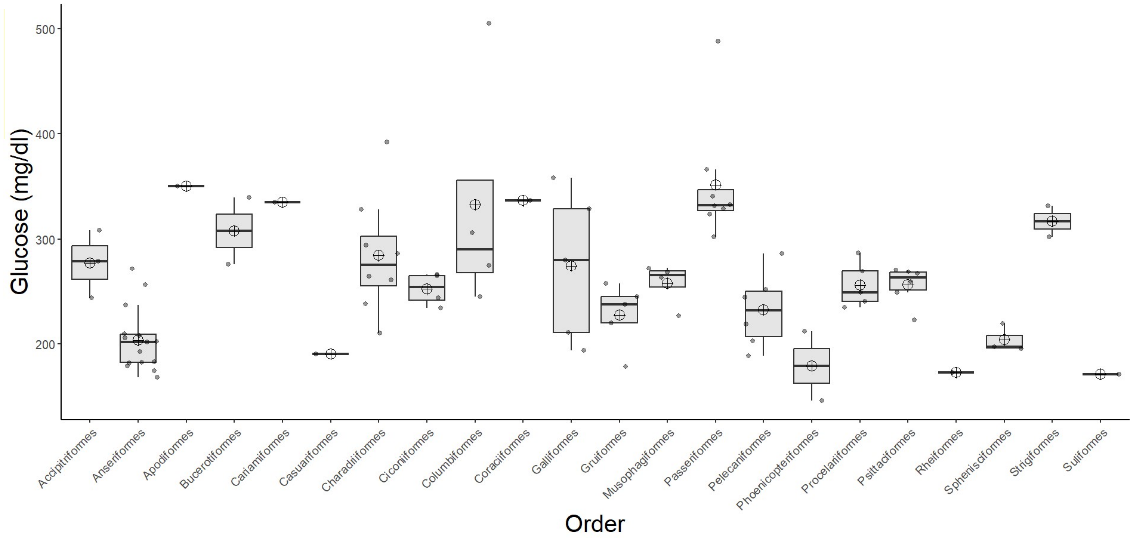

Glucose (mg/dl)
500
400
300
200
Accipitriformes
Anseriformes
Apodiformes
Bucerotiformes
Cariamiformes
Casuariformes
Charadriiformes
Ciconiiformes
Columbiformes
Coraciiformes
Galliformes
Gruiformes
Musophagiformes
Passeriformes
Pelecaniformes
Phoenicopteriformes
Procelariiformes
Psittaciformes
Rheiformes
Sphenisciformes
Strigiformes
Suliformes
Order


A2

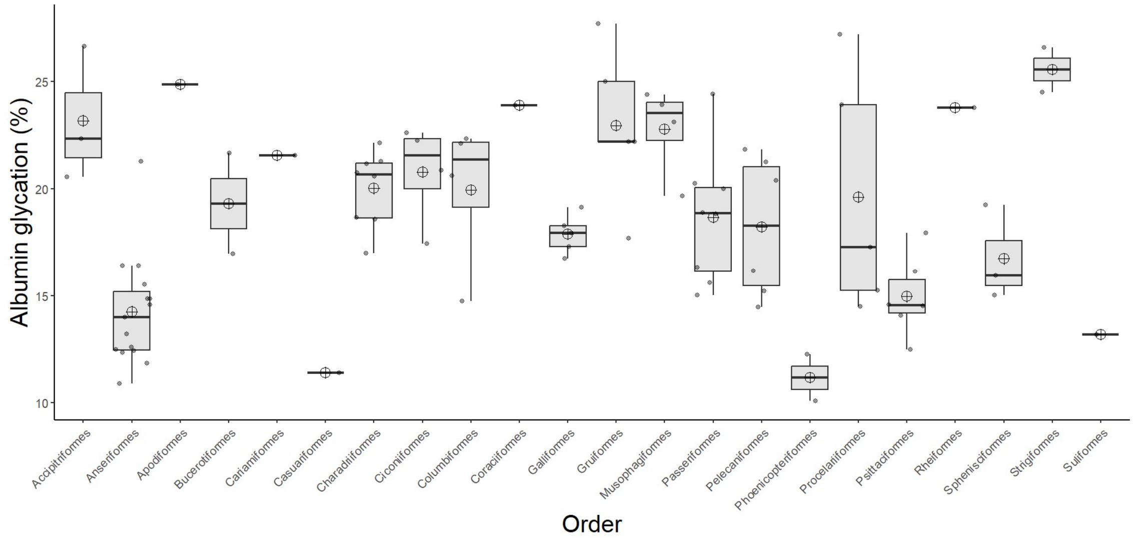

Albumin glycation (%)
25
20
15
10
Accipitriformes
Anseriformes
Apodiformes
Bucerotiformes
Cariamiformes
Casuariformes
Charadriiformes
Ciconiiformes
Columbiformes
Coraciiformes
Galliformes
Gruiformes
Musophagiformes
Passeriformes
Pelecaniformes
Phoenicopteriformes
Procelariiformes
Psittaciformes
Rheiformes
Sphenisciformes
Strigiformes
Suliformes
Order


B1

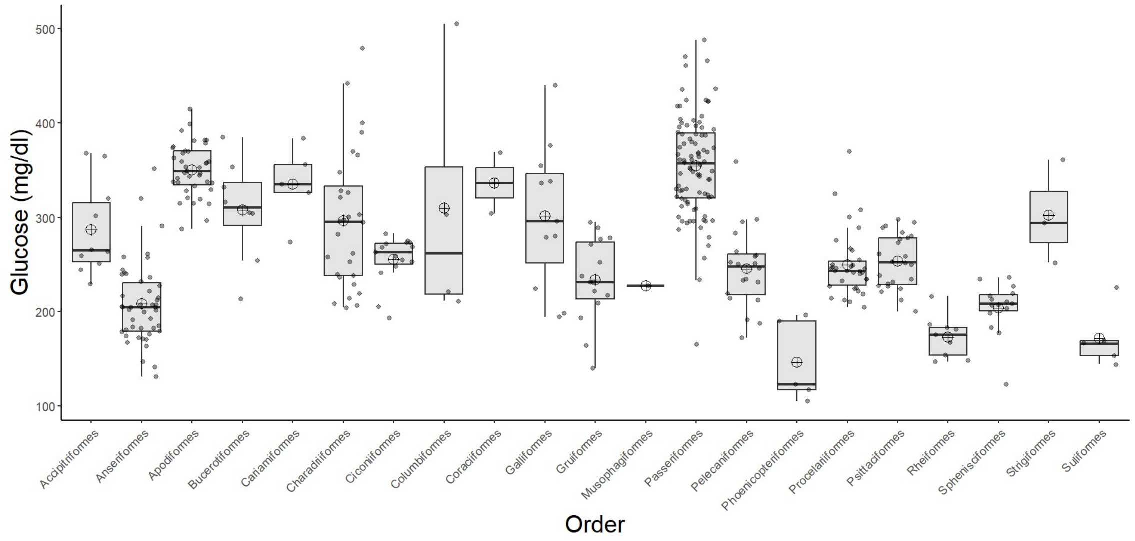

Glucose (mg/dl)
500
400
300
200
100
Accipitriformes
Anseriformes
Apodiformes
Bucerotiformes
Cariamiformes
Charadriiformes
Ciconiiformes
Columbiformes
Coraciiformes
Galliformes
Gruiformes
Musophagiformes
Passeriformes
Pelecaniformes
Phoenicopteriformes
Procelariiformes
Psittaciformes
Rheiformes
Sphenisciformes
Strigiformes
Suliformes
Order

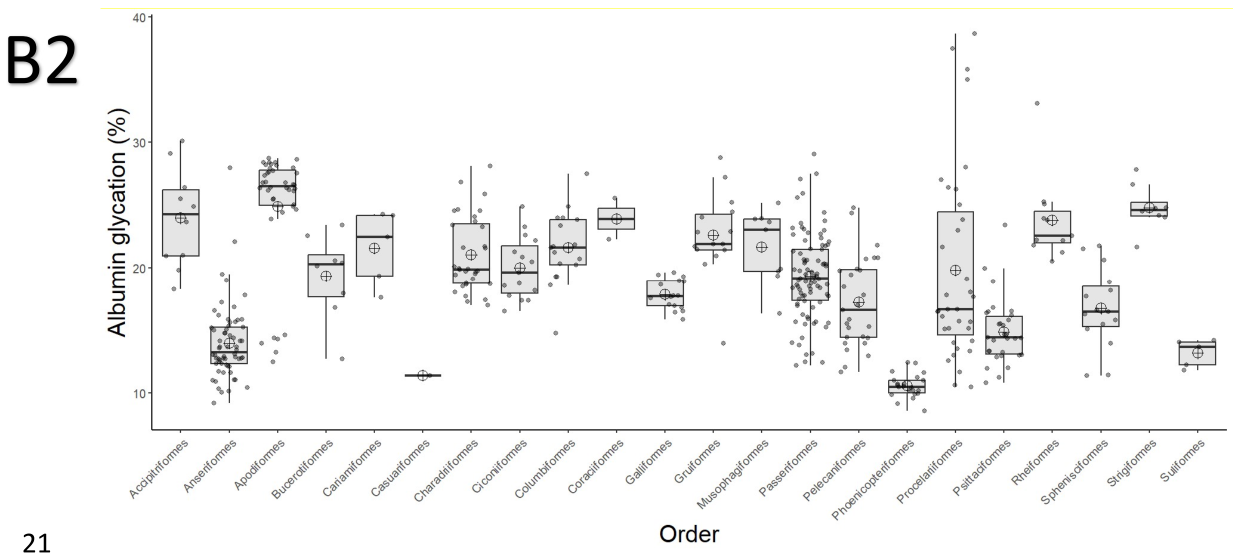


**Figure ESM2.3.** Variation in (A1-B1) plasma glucose levels in mg/dl and (A2-B2) albumin glycation rates across birds' orders included in our study. Panel A represents average species values and B includes individual values.
