## Supplementary material for "Variation in albumin glycation rates in birds suggests resistance to relative hyperglycaemia rather than conformity to the pace of life syndrome hypothesis": ESM1-4 & 6: ESM4.pdf

### <u>Electronic Supplementary Material 4 (ESM4)</u>


#### General

- Bennett, P. M. (1986). Comparative studies of morphology life history and ecology among birds (Doctoral dissertation, University of Sussex).

##### *Aythya baeri*

- Carboneras, C. and G. M. Kirwan (2020). Baer's Pochard (*Aythya baeri*), version 1.0. In Birds of the World (J. del Hoyo, A. Elliott, J. Sargatal, D. A. Christie, and E. de Juana, Editors). Cornell Lab of Ornithology, Ithaca, NY, USA. https://doi.org/10.2173/bow.baepoc1.01

- Kear, J., Editor (2005). Ducks, Geese and Swans. Volume 1: General chapters, and Species accounts (Anhima to Salvadorina). Oxford University Press, Oxford, UK.

##### *Casuarius casuarius*

- Bentrupperbäumer, J. M. (1998). Reciprocal ecosystem impact and behavioural interactions between cassowaries, Casuarius casuarius and humans, Homo sapiens exploring the natural human environment interface and its implications for endangered species recovery in north Queensland, Australia. PhD thesis, James Cook University of North Queensland, Townsville.

- Queensland Parks and Wildlife Service (2001). Recovery plan for the southern cassowary Casuarius casuarius johnsonii 2001–2005. Queensland Parks and Wildlife Service, Brisbane.

- Moore L. (2007). Population ecology of the Southern Cassowary Casuarius casuarius johnsonii, Mission Beach North Queensland. Journal of Ornithology 148(3):357-366.

- Romer, L. (1997). Cassowary Husbandry Manual, Proceedings of February 1996 Workshop. Currumbin Sanctuary, Currumbin.

- Healy, T. P. (2022). Southern Cassowary (Casuarius casuarius), version 2.0. In Birds of the World (S. M. Billerman, Editor). Cornell Lab of Ornithology, Ithaca, NY, USA. https://doi.org/10.2173/bow.soucas1.02

##### *Anas bernieri*

- Carboneras, C. and G. M. Kirwan (2020). Bernier's Teal (Anas bernieri), version 1.0. In Birds of the World (J. del Hoyo, A. Elliott, J. Sargatal, D. A. Christie, and E. de Juana, Editors). Cornell Lab of Ornithology, Ithaca, NY, USA. https://doi.org/10.2173/bow.bertea1.01

##### *Dendrocygna viduata*

- Safford, R. J., and A. F. A. Hawkins, Editors (2013). The Birds of Africa. Volume 8. The Malagasy Region. Christopher Helm, London, UK.

- Carboneras, C. and G. M. Kirwan (2020). White-faced Whistling-Duck (Dendrocygna viduata), version 1.0. In Birds of the World (J. del Hoyo, A. Elliott, J. Sargatal, D. A. Christie, and E. de Juana, Editors). Cornell Lab of Ornithology, Ithaca, NY, USA. https://doi.org/10.2173/bow.wfwduc1.01

##### *Dendrocygna eytoni*

- Carboneras, C. and G. M. Kirwan (2020). Plumed Whistling-Duck (Dendrocygna eytoni), version 1.0. In Birds of the World (J. del Hoyo, A. Elliott, J. Sargatal, D. A. Christie, and E. de Juana, Editors). Cornell Lab of Ornithology, Ithaca, NY, USA. https://doi.org/10.2173/bow.plwduc1.01

***Tachymarptis melba***

- Chantler, P., E. de Juana, G. M. Kirwan, and P. F. D. Boesman (2020). Alpine Swift (Apus melba), version 1.0. In Birds of the World (J. del Hoyo, A. Elliott, J. Sargatal, D. A. Christie, and E. de Juana, Editors). Cornell Lab of Ornithology, Ithaca, NY, USA. https://doi.org/10.2173/bow.alpswi1.01

***Sarkidiornis melanotos***

- Carboneras, C. and G. M. Kirwan (2020). Knob-billed Duck (Sarkidiornis melanotos), version 1.0. In Birds of the World (S. M. Billerman, B. K. Keeney, P. G. Rodewald, and T. S. Schulenberg, Editors). Cornell Lab of Ornithology, Ithaca, NY, USA. https://doi.org/10.2173/bow.comduc2.01

- Demey, R. and Kirwan, G. (2001). Africa Round-up: Duck longevity records. Bull. African Bird Club. 8(1): 6.

***Ara ambiguus***

- Collar, N., P. F. D. Boesman, and C. J. Sharpe (2020). Great Green Macaw (Ara ambiguus), version 1.0. In Birds of the World (J. del Hoyo, A. Elliott, J. Sargatal, D. A. Christie, and E. de Juana, Editors). Cornell Lab of Ornithology, Ithaca, NY, USA. https://doi.org/10.2173/bow.grgmac.01

***Procellaria aequinoctalis***

- Carboneras, C., F. Jutglar, E. de Juana, and G. M. Kirwan (2020). White-chinned Petrel (Procellaria aequinoctialis), version 1.0. In Birds of the World (J. del Hoyo, A. Elliott, J. Sargatal, D. A. Christie, and E. de Juana, Editors). Cornell Lab of Ornithology, Ithaca, NY, USA. https://doi.org/10.2173/bow.whcpet1.01

- Brooke, M. (2004). Albatrosses and Petrels Across the World. Oxford University Press, Oxford, UK.

***Anodorhynchus hyacinthinus***

- Collar, N., P. F. D. Boesman, and C. J. Sharpe (2020). Hyacinth Macaw (Anodorhynchus hyacinthinus), version 1.0. In Birds of the World (J. del Hoyo, A. Elliott, J. Sargatal, D. A. Christie, and E. de Juana, Editors). Cornell Lab of Ornithology, Ithaca, NY, USA. https://doi.org/10.2173/bow.hyamac1.01

***Zenaida graysoni***

- Heck, L. (2020). Socorro Dove (Zenaida graysoni), version 1.0. In Birds of the World (T. S. Schulenberg, Editor). Cornell Lab of Ornithology, Ithaca, NY, USA. https://doi.org/10.2173/bow.socdov1.01

***Aptenodytes patagonicus***

- Martínez, I., F. Jutglar, and E. F. J. Garcia (2020). King Penguin (Aptenodytes patagonicus), version 1.0. In Birds of the World (J. del Hoyo, A. Elliott, J. Sargatal, D. A. Christie, and E. de Juana, Editors). Cornell Lab of Ornithology, Ithaca, NY, USA. https://doi.org/10.2173/bow.kinpen1.01

***Pagodroma nivea***

- Carboneras, C., F. Jutglar, and G. M. Kirwan (2020). Snow Petrel (Pagodroma nivea), version 1.0. In Birds of the World (J. del Hoyo, A. Elliott, J. Sargatal, D. A. Christie, and E. de Juana, Editors). Cornell Lab of Ornithology, Ithaca, NY, USA. https://doi.org/10.2173/bow.snopet1.01

- Amundsen, T. (1995). Egg size and early nestling growth in the Snow Petrel. Condor. 97(2): 345–351.

- Brooke, M. (2004). Albatrosses and Petrels Across the World. Oxford University Press, Oxford, UK.

- Berman et al. (2009). Contrasted patterns of age-specific reproduction in long-lived seabirds. Proceedings of the Royal Society B, 276:375-382

***Cygnus atratus***

- Carboneras, C. and G. M. Kirwan (2020). Black Swan (Cygnus atratus), version 1.0. In Birds of the World (J. del Hoyo, A. Elliott, J. Sargatal, D. A. Christie, and E. de Juana, Editors). Cornell Lab of Ornithology, Ithaca, NY, USA. https://doi.org/10.2173/bow.blkswa.01

***Cacatua moluccensis***

- Rowley, I. and G. M. Kirwan (2020). Salmon-crested Cockatoo (Cacatua moluccensis), version 1.0. In Birds of the World (J. del Hoyo, A. Elliott, J. Sargatal, D. A. Christie, and E. de Juana, Editors). Cornell Lab of Ornithology, Ithaca, NY, USA. https://doi.org/10.2173/bow.saccoc.01

- Juniper, T., and M. Parr (1998). Parrots: A Guide to the Parrots of the World. Pica Press, Robertsbridge, United Kingdom.

***Anser indicus***

- Carboneras, C. and G. M. Kirwan (2020). Bar-headed Goose (*Anser indicus*), version 1.0. In Birds of the World (J. del Hoyo, A. Elliott, J. Sargatal, D. A. Christie, and E. de Juana, Editors). Cornell Lab of Ornithology, Ithaca, NY, USA. https://doi.org/10.2173/bow.bahgoo.01

***Lophura edwardsi***

- McGowan, P. J. K., G. M. Kirwan, and D. A. Christie (2020). Edwards's Pheasant (Lophura edwardsi), version 1.0. In Birds of the World (J. del Hoyo, A. Elliott, J. Sargatal, D. A. Christie, and E. de Juana, Editors). Cornell Lab of Ornithology, Ithaca, NY, USA. https://doi.org/10.2173/bow.edwphe1.01

***Erithacus rubecula***

- Collar, N. (2020). European Robin (Erithacus rubecula), version 1.0. In Birds of the World (J. del Hoyo, A. Elliott, J. Sargatal, D. A. Christie, and E. de Juana, Editors). Cornell Lab of Ornithology, Ithaca, NY, USA. https://doi.org/10.2173/bow.eurrob1.01

***Macronectes halii***

AnAge from ABBBS - Australian Bird and Bat Banding Scheme

***Numida meleagris***

- Martínez, I. and G. M. Kirwan (2020). Helmeted Guineafowl (Numida meleagris), version 1.0. In Birds of the World (J. del Hoyo, A. Elliott, J. Sargatal, D. A. Christie, and E. de Juana, Editors). Cornell Lab of Ornithology, Ithaca, NY, USA. https://doi.org/10.2173/bow.helgui.01

- Urban, E. K., C. H. Fry, and S. Keith, Editors (1986). The Birds of Africa. Volume 2. Academic Press, London, United Kingdom.

***Leptoptilus crumenifer***

- Elliott, A., E. F. J. Garcia, and P. F. D. Boesman (2021). Marabou Stork (Leptoptilos crumenifer), version 1.1. In Birds of the World (Editor not available). Cornell Lab of Ornithology, Ithaca, NY, USA. https://doi.org/10.2173/bow.marsto1.01.1

***Ara glaucogularis***

- Collar, N., P. F. D. Boesman, and C. J. Sharpe (2020). Blue-throated Macaw (Ara glaucogularis), version 1.0. In Birds of the World (J. del Hoyo, A. Elliott, J. Sargatal, D. A. Christie, and E. de Juana, Editors). Cornell Lab of Ornithology, Ithaca, NY, USA. https://doi.org/10.2173/bow.bltmac1.01

- Berkunsky, I., Daniele, G., Kacoliris, F.P., Díaz-Luque, J.A., Silva Frias, C.P., Aramburu, R.M. and Gilardi, J.D. (2014). Reproductive parameters in the critically endangered Blue-throated Macaw: limits to the recovery of a parrot under intensive management. PLOS One. 9(6): e99941. doi:10.1371/journal.pone.0099941.

***Neosenas mayeri***

- Baptista, L. F., P. W. Trail, H. M. Horblit, P. F. D. Boesman, and E. F. J. Garcia (2022). Pink Pigeon (Nesoenas mayeri), version 1.1. In Birds of the World (S. M. Billerman and N. D. Sly, Editors). Cornell Lab of Ornithology, Ithaca, NY, USA. https://doi.org/10.2173/bow.pinpig2.01.1

***Acryllium vulturinum***

- Martínez, I. and G. M. Kirwan (2020). Vulturine Guineafowl (Acryllium vulturinum), version 1.0. In Birds of the World (J. del Hoyo, A. Elliott, J. Sargatal, D. A. Christie, and E. de Juana, Editors). Cornell Lab of Ornithology, Ithaca, NY, USA. https://doi.org/10.2173/bow.vulgui1.01

- Madge, S., and P. McGowan (2002). Pheasants, Partridges and Grouse, including Buttonquails, Sandgrouse and Allies. Christopher Helm, London, United Kingdom.

***Gelochelidon nilotica***

- Molina, K. C., J. F. Parnell, R. M. Erwin, J. del Hoyo, N. Collar, G. M. Kirwan, and E. F. J. Garcia (2020). Gull-billed Tern (Gelochelidon nilotica), version 1.0. In Birds of the World (S. M. Billerman, Editor). Cornell Lab of Ornithology, Ithaca, NY, USA. https://doi.org/10.2173/bow.gubter1.01

- Erwin, R. M., T. B. Eyler, D. B. Stotts and J. S. Hatfield. (1999). Aspects of chick growth in Gull-billed Terns in coastal Virginia. Waterbirds 22 (1):47-53.

- Møller, A. P. (1975c). Ynglebstanden af Sandterne (Gelochelidon nilotica nilotica) Gmel. i 1972 i Europa, Afrika og det vestlige Asien med en oversight over bestandsaendringer i dette arhundrede (The breeding population of Gull-billed Terns Gelochelidon nilotica nilotica Gmel. in 1972 in Europe, Africa and western Asia, with a review of fluctuations during the present century; English summary). Dansk Ornithologisk Forenings Tidsskrift 69:1-8.

***Leucopsar rothschildi***

- Craig, A. J. F., C. J. Feare, and C. J. Sharpe (2020). Bali Myna (Leucopsar rothschildi), version 1.0. In Birds of the World (J. del Hoyo, A. Elliott, J. Sargatal, D. A. Christie, and E. de Juana, Editors). Cornell Lab of Ornithology, Ithaca, NY, USA. https://doi.org/10.2173/bow.balmyn1.01

***Pygoscelis papua***

- Martínez, I., D. A. Christie, F. Jutglar, E. F. J. Garcia, and C. J. Sharpe (2020). Gentoo Penguin (Pygoscelis papua), version 1.0. In Birds of the World (J. del Hoyo, A. Elliott, J. Sargatal, D. A. Christie, and E. de Juana, Editors). Cornell Lab of Ornithology, Ithaca, NY, USA. https://doi.org/10.2173/bow.genpen1.01

***Chionis minor***

- Burger, A. E. and G. M. Kirwan (2020). Black-faced Sheathbill (Chionis minor), version 1.0. In Birds of the World (J. del Hoyo, A. Elliott, J. Sargatal, D. A. Christie, and E. de Juana, Editors). Cornell Lab of Ornithology, Ithaca, NY, USA. https://doi.org/10.2173/bow.blfshe1.01

- Jouventin, P., Bried, J. and Ausilio, E. (1996). Life-history variations of the Lesser Sheathbill Chionis minor in contrasting habitats. Ibis. 138(4): 732–741.

***Mycteria ibis***

- Elliott, A., E. F. J. Garcia, and P. F. D. Boesman (2020). Yellow-billed Stork (Mycteria ibis), version 1.0. In Birds of the World (J. del Hoyo, A. Elliott, J. Sargatal, D. A. Christie, and E. de Juana, Editors). Cornell Lab of Ornithology, Ithaca, NY, USA. https://doi.org/10.2173/bow.yebsto1.01

***Musophaga rossae***

- Turner, D. A., G. M. Kirwan, and P. F. D. Boesman (2021). Ross's Turaco (Musophaga rossae), version 1.1. In Birds of the World (J. del Hoyo, A. Elliott, J. Sargatal, D. A. Christie, and E. de Juana, Editors). Cornell Lab of Ornithology, Ithaca, NY, USA. https://doi.org/10.2173/bow.rostur1.01.1

- Fry, C. H., S. Keith, and E. K. Urban, Editors (1988). The Birds of Africa. Volume 3. Parrots to Woodpeckers. Academic Press, London, UK.

***Acrocephalus scirpaceus***

- Eising, C. M., J. Komdeur, J. Buys, M., Reemer, and D. S. Richardson (2001). Islands in a desert: breeding ecology of the African Reed Warbler Acrocephalus baeticatus in Namibia. Ibis 143(3):482–493.

- Fransson, T., Kolehmainen, T., Moss, D. & Robinson, R. (2023) EURING list of longevity records for European birds.

***Caloenas nicobarica***

- Ricklefs (2000). Intrinsic aging-related mortality in birds. J Avian Biol, 31:103-111.

***Aegypius monachus***

- Salvador, A. (2023). Cinereous Vulture (Aegypius monachus), version 2.0. In Birds of the World (G. M. Kirwan, Editor). Cornell Lab of Ornithology, Ithaca, NY, USA. https://doi.org/10.2173/bow.cinvul1.02

- Eliotout, B., P. Lécuyer, and O. Duriez (2007). Premiers résultats sur la biologie de reproduction du vautour moine Aegypius monachus en France. Alauda 75(3):253–264.

- de la Puente, J., A. Bermejo, J. C. del Moral, and A. Ruiz (2011). Juvenile dispersion, dependence period, phylopatry and breeding maturity age of the cinereous vulture. In Ecología y conservación de las rapaces forestales europeas (I. Zuberogoitia and J. E. Martínez, Editors), Departamento de Agricultura de la Diputación Foral de Bizkaia, Bilbao, Spain. pp. 270–280.

***Vanellus miles***

- del Hoyo, J., P. Wiersma, G. M. Kirwan, and N. Collar (2020). Masked Lapwing (Vanellus miles), version 1.0. In Birds of the World (S. M. Billerman, B. K. Keeney, P. G. Rodewald, and T. S. Schulenberg, Editors). Cornell Lab of Ornithology, Ithaca, NY, USA. https://doi.org/10.2173/bow.maslap1.01

- ABBBS - Australian Bird and Bat Banding Scheme

***Cariama cristata***

- Gonzaga, L. P. and G. M. Kirwan (2020). Red-legged Seriema (Cariama cristata), version 1.0. In Birds of the World (J. del Hoyo, A. Elliott, J. Sargatal, D. A. Christie, and E. de Juana, Editors). Cornell Lab of Ornithology, Ithaca, NY, USA. https://doi.org/10.2173/bow.relser1.01

***Gerontichus eremita***

- Matheu, E., J. del Hoyo, G. M. Kirwan, and E. F. J. Garcia (2020). Northern Bald Ibis (Geronticus eremita), version 1.0. In Birds of the World (J. del Hoyo, A. Elliott, J. Sargatal, D. A. Christie, and E. de Juana, Editors). Cornell Lab of Ornithology, Ithaca, NY, USA. https://doi.org/10.2173/bow.waldra1.01

- Reşit Akçakaya, H. (1990). Bald Ibis Geronticus eremita population in Turkey: An evaluation of the captive breeding project for reintroduction. Biological Conservation, 51(3), 225–237. doi:10.1016/0006-3207(90)90153-g

***Burhinus grallarius***

- Hume, R., G. M. Kirwan, and P. F. D. Boesman (2020). Bush Thick-knee (Burhinus grallarius), version 1.0. In Birds of the World (J. del Hoyo, A. Elliott, J. Sargatal, D. A. Christie, and E. de Juana, Editors). Cornell Lab of Ornithology, Ithaca, NY, USA. https://doi.org/10.2173/bow.butkne1.01

***Balearica regulorum***

- Archibald, G.W., C.D. Meine, and E. F. J. Garcia (2020). Gray Crowned-Crane (Balearica regulorum), version 1.0. In Birds of the World (J. del Hoyo, A. Elliott, J. Sargatal, D. A. Christie, and E. de Juana, Editors). Cornell Lab of Ornithology, Ithaca, NY, USA. https://doi.org/10.2173/bow.grccra1.01

- Ricklefs (2000). Intrinsic aging-related mortality in birds. Journal of Avian Biology, 31:103-111

***Psophia crepitans***

- Potter, A. B. (2020). Gray-winged Trumpeter (Psophia crepitans), version 1.0. In Birds of the World (T. S. Schulenberg, Editor). Cornell Lab of Ornithology, Ithaca, NY, USA. https://doi.org/10.2173/bow.gywtru1.01

- Horning, C.L., Hutchins, M. and English, W. (1988), Breeding and management of the common trumpeter (Psophia crepitans). Zoo Biol., 7: 193-210. https://doi.org/10.1002/zoo.1430070302

- Male, J. P. (1989). Foster parenting, growth and management of the Common trumpeter, Psophia crepitans, at the National Zoological Park, Washington. International Zoo Yearbook.

***Neophron percnopterus***

- García-Ripollés, C., López-López, P. Integrating effects of supplementary feeding, poisoning, pollutant ingestion and wind farms of two vulture species in Spain using a population viability analysis. J Ornithol 152, 879–888 (2011). https://doi.org/10.1007/s10336-011-0671-8

***Rhea pennata***

- Sales, J. (2006). The rhea, a ratite native to South America. Avian and Poultry Biology Reviews 17(4):105–124.

***Dacelo novaeguineae***

- Woodall, P. F. (2020). Laughing Kookaburra (Dacelo novaeguineae), version 1.0. In Birds of the World (J. del Hoyo, A. Elliott, J. Sargatal, D. A. Christie, and E. de Juana, Editors). Cornell Lab of Ornithology, Ithaca, NY, USA. https://doi.org/10.2173/bow.laukoo1.01

***Rhea americana***

- Kirwan, G. M., A. Korthals, and C. E. Hodes (2021). Greater Rhea (Rhea americana), version 2.0. In Birds of the World (B. K. Keeney, Editor). Cornell Lab of Ornithology, Ithaca, NY, USA. https://doi.org/10.2173/bow.grerhe1.02

- Sick, H. (1993). Birds in Brazil: A Natural History. Princeton University Press, Princeton, NJ, USA.

- Bruning, D. F. (1973). Breeding and rearing rheas in captivity. International Zoo Yearbook 13:163–172.

- Navarro, J. L., P. E. Vignolo, M. R. Demaría, N. O. Maceira, and M. B. Martella (2005) Growth curves of farmed Greater Rheas (Rhea americana albescens) from central Argentina. Archives für Geflugelkunde 69:90–93.

***Anthropoides paradiseus***

- Archibald, G.W., C.D. Meine, and E. F. J. Garcia (2020). Blue Crane (Anthropoides paradiseus), version 1.0. In Birds of the World (J. del Hoyo, A. Elliott, J. Sargatal, D. A. Christie, and E. de Juana, Editors). Cornell Lab of Ornithology, Ithaca, NY, USA. https://doi.org/10.2173/bow.blucra2.01

***Anthropoides virgo***

- Archibald, G.W., C.D. Meine, E. F. J. Garcia, and G. M. Kirwan (2020). Demoiselle Crane (Anthropoides virgo), version 1.0. In Birds of the World (J. del Hoyo, A. Elliott, J. Sargatal, D. A. Christie, and E. de Juana, Editors). Cornell Lab of Ornithology, Ithaca, NY, USA. https://doi.org/10.2173/bow.demcra1.01

***Gyps fulvus***

- Salvador, A. (2022). Eurasian Griffon (Gyps fulvus), version 3.0. In Birds of the World (S. M. Billerman and M. A. Bridwell, Editors). Cornell Lab of Ornithology, Ithaca, NY, USA. https://doi.org/10.2173/bow.eurgri1.03

- Sarrazin, F., C. Bagnolini, J. L. Pinna, and E. Danchin (1996). Breeding biology during establishment of a reintroduced Griffon Vulture Gyps fulvus population. Ibis 138: 315–325.

- Adamian, M. S., and D. Klem (1999). Handbook of the Birds of Armenia. American University of Armenia, Yerevan, Armenia.

***Machronectes giganteus***

- Carboneras, C., F. Jutglar, and G. M. Kirwan (2020). Southern Giant-Petrel (Macronectes giganteus), version 1.0. In Birds of the World (J. del Hoyo, A. Elliott, J. Sargatal, D. A. Christie, and E. de Juana, Editors). Cornell Lab of Ornithology, Ithaca, NY, USA. https://doi.org/10.2173/bow.angpet1.01

- Foote et al. (2011). Individual state and survival prospects: age, sex, and telomere length in a long-lived seabird. Behavioral Ecology, 22.

***Tauraco fischeri***

- Turner, D. A. and P. F. D. Boesman (2020). Fischer's Turaco (Tauraco fischeri), version 1.0. In Birds of the World (J. del Hoyo, A. Elliott, J. Sargatal, D. A. Christie, and E. de Juana, Editors). Cornell Lab of Ornithology, Ithaca, NY, USA. https://doi.org/10.2173/bow.fistur1.01

***Tauraco leucolophus***

- Turner, D. A. and P. F. D. Boesman (2020). White-crested Turaco (Tauraco leucolophus), version 1.0. In Birds of the World (J. del Hoyo, A. Elliott, J. Sargatal, D. A. Christie, and E. de Juana, Editors). Cornell Lab of Ornithology, Ithaca, NY, USA. https://doi.org/10.2173/bow.whctur2.01

***Branta ruficollis***

- Carboneras, C., G. M. Kirwan, and C. J. Sharpe (2020). Red-breasted Goose (*Branta ruficollis*), version 1.0. In Birds of the World (J. del Hoyo, A. Elliott, J. Sargatal, D. A. Christie, and E. de Juana, Editors). Cornell Lab of Ornithology, Ithaca, NY, USA. https://doi.org/10.2173/bow.rebgoo1.01

***Larus dominicanus***

- ABBBS - Australian Bird and Bat Banding Scheme
