## Supplementary material for "Variation in albumin glycation rates in birds suggests resistance to relative hyperglycaemia rather than conformity to the pace of life syndrome hypothesis": ESM1-4 & 6: ESM6.pdf

### Electronic Supplementary Material 6 (ESM6)

#### Study species

Of all the species on the study, 65 came from captive populations of the Mulhouse zoo (Mulhouse, France) and the Parc des Oiseaux (Villars-les-Dombes, France). These samples were collected during annual vaccination campaigns in October 2018 and 2021. Zebra finch (*Taeniopygia guttata*) samples were taken in August-September 2022 from individuals kept in captivity in the animal facilities of the DEPE-IPHC (Department of Ecology, Physiology and Ethology, Institut Pluridisciplinaire Hubert Curien, Strasbourg, France). Alpine swift (*Tachymarptis melba*) samples were collected in May and August 2023 from wild individuals breeding in colonies in Switzerland (see e.g. [1-3]). Adult black-tailed godwits (*Limosa limosa*) were sampled in February 2022 in Extremadura rice fields, southwestern Spain. Adult gull-billed terns (*Gelochelidon nilotica*) were sampled in May-June 2022 from a breeding colony located at Villalba de los Barros reservoir in Extremadura, southwestern Spain. All passerine samples, excluding zebra finches and Bali myna (*Leucopsar rothschildii*), were obtained from wild individuals captured using mist nets during a migration monitoring project in a wetland at Sainte-Soline (France) in August 2022. Scopoli's shearwater (*Calonectris diomedea*) samples were collected in 2011 in the Chafarinas Islands, Spain. The remaining samples from Procellariiformes species, penguins (Sphenisciformes), cormorants (Suliformes) and all marine Charadriiformes were collected from wild populations in the French Southern Territories of Crozet and Kerguelen during 2022. Sample sizes for each species measured by our team went from 1 to 60 for glucose (mean=5.32; σ=7.07; median=4.5; mode=5) and from 1 to 55 for glycation (mean=5.19; σ=7.9; median=4; mode=5). Excepting zebra finches (*Taeniopygia guttata*) (60 for glucose, 55 for glycation) and Alpine swifts (*Tachymarptis melba*), (39 for glucose, 40 for glycation) whose sample sizes are bigger because these measurements were originally collected on untreated animals for separate analyses (unpublished data), the values are lower and vary much less in number (1-16 for both glucose and glycation), and are limited in size by logistical reasons, mainly related to mass-spectrometry operation. The values reported here for glycation are counting the individuals included in the statistical analyses, not the ones sampled, as some individuals had glycation values under the limit of detection, that were eliminated from the analyses, as stated in the main text. However, when considering glucose values coming from the 13 species from ZIMs database, the sample sizes were bigger 17-387 individuals, (mean=90.46; σ=26.31; median=55; see main text). The validity of the glucose and glycation values from our dataset to represent the species (as sample size per species was often low) was assessed by performing general linear mixed models (GLMM; *lmer()* function from *lme4* package in R [4]) with either glucose or glycation as dependent variables and "species" as a random factor for the intercept only, and then obtaining the repeatability score with *rptGaussian()* from *rptR* package [5] in order to determine if the variability between species is higher than within species.

#### Possible effects of stress

Although glucose levels are known to be affected by the stress response, and thus by handling time (see e.g. [6-8] for birds), and time between bird capture and blood sampling ('handling time') is not measured here, there are some reasons to consider that our results are still robust. First, most individuals were sampled shortly after captured, so that the stress response could not play a big role on glucose variation. Second, we considered that this change in glucose levels with stress, at least in certain species, may be driven more by an increase in variation than by an increase in average values, which means that our study would remain conservative, so that the probability of finding significant results is lower, but the patterns of variation that we did find are more robust. We base this idea on an analysis we did on publicly available data from 160 species of Passeriformes from [9], in which we performed a LMM on glucose with sampling time in min (0, 15 and 30) as fixed factor (see **Figure ESM1.2**), and individual nested within species as random factor, showing a clear heteroskedasticity across sampling times, and a model without the fixed factor.  This way, we also compared the percentages of variance explained by the species and individual when considering all reported glucose values (i.e. at all sample times) together with the variance when the time effect was accounted for (see ESM1), showing a lower species repeatability when sampling time is not considered. Moreover, their species-explained variance when sampling time is not considered is lower than in our study, probably in relation to our higher taxonomic cover. Given the mentioned heteroskedasticity of the data, a Kruskal-Wallis test (non-parametric) was performed to explore the differences in glucose values across sampling times, reporting a significant result (see ESM1), which indicates differences across the times, that are nevertheless not necessarily associated with different means. Third, preliminary analyses performed by our team on data from Alpine swifts and zebra finches (unpublished results), in which many individuals were sampled during the same session, and therefore differences in stress levels across individuals may be present, show no significant effect of time on glucose variation, particularly when the individuals were overnight fasted, as it is the case for zebra finches.

#### Diet data clarifications

In many cases, adjustments were necessary because the criteria used by [10] to classify species (subsequently used in AVONET) were based on the ecological niche and its relationship with morphology rather than considering the potential association of dietary composition with physiological variables. To address this limitation, we used the “Trophic.Niche” variable from AVONET and merged some categories: “Vertivore”, “Invertivore” and “Scavenger” into “Carnivore terrestrial”, “Frugivore” and “Granivore” into “Frugivore/granivore” and “Herbivore terrestrial” and “Herbivore aquatic” into “Herbivore”, due to lack of sufficient species on each separate category (as in “Herbivore terrestrial” and “Herbivore aquatic”) and sometimes on difficulties to tease apart the diet categories for our species (as in “Frugivore” and “Granivore”, as most of our species classified within these groups had a 50%-50% or similar distribution in their food source between both categories, zebra finches being the only exception being exclusively granivorous). Furthermore, some considered in AVONET as “Omnivores” were reclassified, and only considered “Omnivores” if diet was widely distributed between animal and plant sources, and not only within animal or plant categories (e.g. a species that eats invertebrates, vertebrates and carrion was not considered by us as an “Omnivore”, but as either an “Aquatic predator” or a “Carnivore terrestrial”, depending on its main feeding source).

#### Statistics

##### Life history traits

Given that all the 3 life history parameters (maximum lifespan, clutch mass and developmental time) used in our analyses are widely recognized to be strongly correlated with body mass, and interpretations of the variation in life history traits can change if body mass is not adjusted for [11] we chose to adjust the life history traits by body mass prior to run our analyses, as done by [12], the most similar study on glucose levels in birds in function of life history, so that our results are easily comparable. This was done by $\log_{10}$ transforming all variables and performing a phylogenetically controlled Generalized Least Squares model (pGLS, *gls* function in R; [13]) including a correction for phylogeny (assuming a Brownian model and the same tree as described in the main text for the MCMC GLMMs), from which the residuals were extracted. These residuals were the variables used in the main models when referring to “life history traits”, and they are also the same thing as when referring to mass-adjusted variables across the text. Maximum lifespan was used here as a proxy for longevity. Despite concerns about its use (e.g. [14-15]), it remains the most widely available longevity-related data in many databases for a large number of species. Clutch mass, calculated by multiplying clutch size by egg mass (as in [12], it was used as an indicator of reproductive investment. Finally, developmental time (incubation time plus number of days to fledging) was used as a proxy for growth, given that growth rate was difficult to calculate due to lack of information on fledging mass. These variables were also chosen as they were among the ones available for a greater number of species from the sources we used, and for comparative purposes with [12]. Including more variables as e.g. number of clutches per year (one of the other available variables for a considerable number of species) would have further reduced the sample size without, in our opinion, adding much more information, as reproductive effort is already accounted for in the clutch mass variable.

#### Main models

Models were performed in the following way (see **Table ESM6. 1**): first, in order to account for all the species in the dataset (as individual values were not available for all the species, with no individual at all in some cases; see ***Species and sample collection*** section from main text methods) with average glucose and then with average glycation as dependent variables, and later with the individual values in another set of models. Body mass and diet were always included as independent variables. All of this was repeated also including life history traits but with the 66 species for which data were available (only species for which we had all the selected life-history variables were kept). Glycaemia and body mass were $\log_{10}$ transformed and centred in all the models (centring of glucose only occurred when it was a covariable) in order to better interpret the intercepts [16]. In models predicting glycation, plasma glucose was also added as a covariate. For these particular models, linear (Glycation ~ Centred glucose) and logarithmic ($\log_{10}$Glycation ~ $\log_{10}$(Centred glucose)) relationships were assessed, comparing the DIC (Deviance Information Criteria) of the different models, log-log relationship showing much lower DIC than the linear relationship. Semilogarithmic relationships (both Glycation ~ $\log_{10}$(Centred Glucose) and $\log_{10}$Glycation ~ Centred glucose) were also assessed, but DICs were similar between these and either the linear or the log-log model, with much lower DIC for the ones using $\log_{10}$Glycation as dependent variable. Also, all these models showed similar results regarding the significant effects found, and if glucose and glycation were standardized, the estimates of the slopes were very similar for all of them. Therefore, the log-log model was chosen. Identical models without including glucose as a covariate were also performed (for the glycation models including life-history traits) to determine if glycation itself (not glycation resistance, as defined in the main text), showed a covariation with life-history. Quadratic component of maximum longevity was also included in glucose and glycation models including life-history data, after data exploration suggested such pattern. The potential effects of captivity

were also tested by a factor with two levels indicating the provenance of the samples (captive or wild), in preliminary analyses with a lower number of samplings ($10^4$ iterations, burnin=1000). As no effects were found for this variable in the glycation models, it was only maintained in the glucose models thereafter. Besides, in glucose models, the methodology used for determining it (glucometer or kit) was included as a random factor, in order to control for a potential bias in the results. The species was added as a random factor (given a tree provided within the *pedigree* argument of the *MCMCglmm* function).
Since glucose values were not available for all individuals, our models addressing intraspecific variability were limited to 379 individuals from 75 species when considering only diet and body mass predictors (and glucose when modelling glycation). Similarly, only 316 individuals from 58 species were incorporated into the models when life-history traits were introduced. The species identity was always included as a random factor twice: one to control for the phylogeny and the other to avoid the pseudo-replication effect of having repeated measures within species. The method used to determine plasma glucose levels was not included in this case, since here the glucometer was used in all cases.

Gaussian distribution was always assumed for the models, as even if glycation values are, strictly speaking, proportions, not only they follow a continuous distribution (as their calculation come from the division of area values that vary in a continuous way; see main text), but they vary within a limited range (8.57-38.65 %, mean=18.8 %, $\sigma$=5.3) never sufficiently close to 0 or 1 and also do not represent probabilities given by observations of cases as to be treated as binomial with log link function. Furthermore, we believe that there are some reasons why logit transformation, as previously also recommended for proportional data [17], is not the best option in this case: first, logit calculations are suitable for exploring how probabilities of certain outcomes differ between groups, which is not the case in our analyses; second, exploration of the data suggested this transformation did not markedly alter the variance structure of the variable, which was close to a gaussian curve, while making more difficult the interpretation of the outcomes of the models.

Model convergence and lack of autocorrelation was checked by visual inspection of trace of the model MCMC simulations (see trace and density plots of posteriors in **Figure ESM1.1**).

#### Models on age and sex, on lysine residues and on orders

We performed supplementary analyses only with relative age (when it was known, in most cases for birds born in captivity, or belonging to wild populations with demographic monitoring), including a quadratic term, and sex of the individuals (as provided by the caregivers at the zoos or the person in charge of organizing the sample collection and known by direct phenotypic observation when existent sexual dimorphism allows it or by molecular sexing with PCR when not), as predictive variables of glucose and glycation levels. Relative age was defined as its proportion to maximum lifespan of the species (i.e. divided by it), and logit transformation was performed for this ratio to avoid the particular distribution characteristics of proportion data, as Gaussian distribution was assumed in the models. We included glucose as a covariate for the glycation models, and body mass in every case, when it was available for the individuals. These models included a total of 239 individuals from 49 species (see **Table ESM6. 1.**).

We performed a model (MCMCglmm with species as a random factor) comparing the number of lysines exposed in the albumin of each species with the glycation levels measured in our study. This analysis aimed to assess whether this factor could represent a meaningful mechanism of resistance to glycation, as outlined in [18]. When the albumin amino acids sequence for a species we measured was not available on NCBI or UniprotKB (the sources used to determine the number of lysines of each species' albumin; see below for additional details), we selected a closely related species, typically from the same genus (see **Table ESM6. 2.** for a list of the species selected and their correspondence to ours). In total, 19 species were included in this analysis.

Finally, the models on glucose and glycation values controlling for the bird orders included in the dataset (see ESM5) were also performed as MCMCglmm, with two models considering the species averages (one for glucose and one for glycation) and two including the intraspecific values and therefore the species as a random factor to control for pseudo-replication due to multiple sampling of the same species. The species were allocated to the same orders employed on the classification given by the most updated version of Birds of the World [19]. The orders were organized in the model by alphabetic order, Accipitriformes constituting thus the intercept of all of them.

**Table ESM6. 1.** Main set of models performed with the number of species and individuals included in them. A total of 10 MCMCglmm models were performed, having either plasma glucose or albumin glycation levels as response variable and a different number of species and total datapoints depending on if the life-history (LH) traits were included (i.e. maximum lifespan, clutch mass and developmental time) or only diet and body mass (also glucose in the glycation models). The models considering age and sex did not include life history traits nor diet.

| **Models performed and N for each** | **Species averages** | | | | **Intraspecific variation** | | | | **Age and sex** | |
|---|---|---|---|---|---|---|---|---|---|---|
| | Glucose | | Glycation | | Glucose | | Glycation | | Glucose | Glycation |
| | Non LH | LH | Non LH | LH | Non LH | LH | Non LH | LH | | |
| **Number of species** | 88 | 66 | 88 | 66 | 75 | 58 | 75 | 58 | 49 | 49 |
| **Number of individuals** | | | | | 379 | 316 | 379 | 316 | 239 | 239 |

#### Quantification of exposed lysine residues

The aminoacidic sequences of albumins were searched in UniprotKB and the NCBI database. For twenty species, the sequences were submitted to the PHYRE2 server to generate a PDB structure. Then, using DEPTH, the accessible surface area (ASA) was calculated for each residue in the sequences. We used a custom Python script to determine whether each lysine residue is exposed or not. To achieve this, the ASA of the lysine side chain was divided by 205 angstroms (the maximum observed accessible surface area for a lysine [20-21]). If it exceeded 0.25, the residue was considered exposed. This methodology allowed us to determine the count of exposed lysine residues in our avian albumin sequences.

PHYRE2: http://www.sbg.bio.ic.ac.uk/~phyre2/html/page.cgi?id=index

DEPTH: https://bio.tools/depth ; http://mspc.bii.a-star.edu.sg/depth (Depth web server is actually dead)

**Table ESM6. 2.** List of species used in the models testing the relationship between the number of albumin's exposed lysines and its glycation rates. On the left, the list of considered species and on the right, the list of species used as references for the number of exposed lysines. The ones in bold are those that coincide.

| Species in our dataset | Original species |
|---|---|
| **Anas platyrhynchos** | **Anas platyrhynchos** |
| Anser anser | Anser brachyrhynchus* |
| Anser indicus | Anser cygnoides* |
| **Aptenodytes patagonicus** | **Aptenodytes patagonicus** |
| Tachimarptos melba | Apus apus |
| Aythya baeri | Aythya fuligula |
| **Balearica regulorum** | **Balearica regulorum** |
| **Bubo bubo** | **Bubo bubo** |
| **Cariama cristata** | **Cariama cristata** |
| **Cygnus atratus** | **Cygnus atratus** |
| **Eudyptes chrysolophus** | **Eudyptes chrysolophus** |
| Limosa limosa | Limosa lapponica |
| **Numida meleagris** | **Numida meleagris** |
| Leucocarbo verrucosus | Phalacrocorax carbo |
| **Phoenicopterus ruber** | **Phoenicopterus ruber** |
| **Pygoscelis papua** | **Pygoscelis papua** |
| **Rhea pennata** | **Rhea pennata** |
| **Taeniopygia guttata** | **Taeniopygia guttata** |
| **Tauraco erythrolophus** | **Tauraco erythrolophus** |
| Turdus merula | Turdus rufiventris |

* For these species, as they both belong to the same genus, an average was calculated for both the number of exposed lysines and the glycation levels, considering them in the analyses as one. This way, *Aser anser* and *Anser indicus* were substituted by an *Anser sp.* average glycation value, and *Anser brachyrhynchus* and *Anser cygnoides* by an average number of lysine exposed. The actual values were very similar in both cases, so this average is likely to reflect realistic values.

[12]. Tomášek, Oldřich, Lukáš Bobek, Tereza Kauzálová, Marie Adamkova and Tomáš Albrecht. 2019. Fuel for the pace of life: baseline blood glucose concentration co-

evolves with life-history traits in songbirds. *Functional Ecology*, 33 (2): 239–49. https://doi.org/10.1111/1365-2435.13238
